# Demonstrating and disrupting well-learned habits

**DOI:** 10.1101/745356

**Authors:** Ahmet O. Ceceli, Catherine E. Myers, Elizabeth Tricomi

## Abstract

Researchers have exerted tremendous efforts to empirically study how habits form and dominate at the expense of deliberation, yet we know very little about breaking these rigid habits to restore goal-directed control. In a three-experiment study, we first illustrate a novel approach of studying well-learned habits, in order to effectively demonstrate habit disruption. In Experiment 1, we use a Go/NoGo task with familiar color-response associations to demonstrate outcome-insensitivity when compared to novel, more flexible associations. Specifically, subjects perform more accurately when the required mapping is the familiar association of green–Go/red–NoGo than when it is red–Go/green–NoGo, confirming outcome-insensitive, habitual control. As a control condition, subjects show equivalent performance with unfamiliar color-response mappings (using the colors blue and purple mapped to Go and NoGo responses). Next, in Experiments 2 and 3, we test a motivation-based feedback manipulation in varying magnitudes (i.e., performance feedback with and without monetary incentives) to break the well-established habits elicited by our familiar stimuli. We find that although performance feedback prior to the contingency reversal test is insufficient to disrupt outcome-insensitivity in Experiment 2, a combination of performance feedback and monetary incentive is able to restore goal-directed control in Experiment 3, effectively breaking the habits. As the first successful demonstration of well-learned habit disruption in the laboratory, these findings provide new insights into how we execute and modify habits, while fostering new and translational research avenues that may be applicable to treating habit-based pathologies.

## Introduction

When categorizing motivated behaviors, habits are distinguished from goal-directed actions in that they are performed reflexively in response to a triggering cue, without consideration of the consequences [1]. These habitual behaviors are less cognitively taxing than their goal-directed counterparts, allowing for their utilization in instances where the resource-consuming reflection of potential outcomes may not be ideal [2–4]. For example, looking both ways before crossing a street is an action best elicited habitually, and ideally should persist despite the absence of oncoming traffic. In contrast, the optimal motivational control system for commuting to a new destination would be outcome-reliant, reflective, and thus resource-consuming goal-directed performance.

For decades, the motivational bases of behavioral control (i.e., goal-directed and habitual actions) have been investigated in rodent models. In a typical study examining habitual control, a neutral stimulus (e.g., a visual cue, or the context of the chamber) signals hungry rats to press a lever in pursuit of a food outcome. This behavioral training period is often followed by a devaluation procedure—the rat is allowed free-access to the food, promoting satiation and diminishing the food’s value (hence the term *devaluation*). In a subsequent, unrewarded, extinction phase, the experimenter can then assess whether the trained lever-press action is flexible and goal-directed (i.e., strong responses when animal is hungry but diminished responses when satiated), or rigid and habitual (i.e., persistent responses regardless of satiation) [5]. Generally, over-training of the stimulus—response—outcome association tends to render actions habitual. Thus, an over-trained rat persists in pressing the lever despite a diminished value in outcome, suggesting that the actions are driven by the preceding cue or the chamber context. In contrast, value-driven goal-directed control survives following moderate experience with the stimulus–response–outcome chain [6]. Motivational control testing in humans has followed suit with similar operant conditioning paradigms, in which a primary or a secondary reward is devalued to determine whether actions are cue or value driven [7–12]. Another widely-used example is the sequential decision task, in which subjects respond to probabilistic multi-step associative sequences and recruit model-based (i.e., goal-directed; taking into account the cognitive model of the task environment) or model-free (i.e., similar to habits; actions based solely on history of reward receipt) strategies to maximize gain and minimize loss [13].

These methods have undoubtedly contributed a great deal to our understanding of habits; however, such paradigms are limited in critical aspects. Indeed, the habit experience has been a difficult construct to effectively capture via behavioral paradigms in humans [14–16]. First, in contemporary paradigms, including those based on outcome-devaluation and sequential decision-making, the agent must develop a newly formed habit. Accordingly, the tools at our disposal facilitate the study of novel, lab-developed habits, while leaving incomplete our understanding of well-learned habits that are more representative of daily experiences. For example, especially in outcome-devaluation tasks involving valued and devalued food rewards, testing whether a behavior is habitual relies on several critical factors. The demonstration of a habit may depend on successful over-training of a new cue–response–outcome association that develops a strong enough link between the cue and the response to guide behavior [11]. Furthermore, the effectiveness of the devaluation procedure where a food outcome is selectively fed to diminish its value may become problematic in humans for reasons not encountered in rats, such as demand characteristics, and hesitation to eat copious amounts of junk food in a potentially socially intimidating lab setting. Lastly, the experimenter makes assumptions of comparable food palatability, in that the agent must value the food options similarly prior to selective devaluation for any value-based manipulation to be effective [11]. These lab-generated habits are also arduous to develop via over-training, especially in expensive neuroimaging contexts. More importantly, the strength of the trained habit would be insufficient for a meaningful investigation of the habit-breaking process, in that even multi-day training is often measured in minutes to hours [11,17]. Thus, the current tools provide a costly platform that only captures the unidirectional shift from goal-directed to habitual control [18]. In other words, although these novel, lab-created associations permit the study of habit formation and execution, we are limited in our tools to investigate habit disruption with similar efficacy.

Despite tremendous efforts directed towards understanding habit formation and expression, a wider gap in the literature remains regarding the breaking of habits. Accessing the shift from habitual to goal-directed control may ultimately facilitate interventions that remediate rigid and maladaptive behaviors, yet we are not currently methodologically equipped to tackle this translational research avenue with a rich toolkit. Accordingly, we propose that developing a novel habit from an action–outcome contingency is not a pre-requisite for studying the motivational basis for habits, but that an existing, more robust habit could be examined in the lab with less effort. An effective approach may involve using salient cues that elicit well-established, habit-like behaviors that are impervious to their consequences. For instance, the colors red and green have highly specific “stop” and “go” associations, possibly strengthened in a variety of contexts including traffic lights, visual signals of danger and safety, and childhood games, songs, and stories [19]. The familiar red–stop and green–go contingencies have previously been transformed into Go/NoGo tasks to assess response inhibition via perseverative errors (i.e., NoGo accuracy) [19–21]. Similarly, we can test for behavioral rigidity by assessing performance when these contingencies are congruent with daily experiences versus when adjusted to reflect outcomes incongruent with most real-world scenarios. Thus, instead of devaluing the palatability of a primary reward, we render a well-learned association inappropriate for optimal task performance. The agent must override a prepotent red stimulus–stop response with an incongruent green stimulus–stop response to achieve the intended, correct outcome. A more pronounced accuracy impairment when managing incongruencies within this well-learned color-response mapping, compared to changes in a newly-acquired mapping, would permit us to conclude that these familiar stimuli evoke outcome-insensitive actions, the hallmark of habitual behavior. Upon establishing that these familiar stimuli elicit habitual control, we can then provide the platform to study habit disruption by testing manipulations that protect against mapping-related performance impairments–essentially overriding the habitual response by engaging cognitive control processes. The motivational control framework identifies habits as cue-dependent, and goal-directed behaviors as those contingent on the outcome [1]. Accordingly, a previously goal-directed behavior is rendered habitual when the associative strength of the stimulus–response component governs actions, rendering the outcome inessential for action execution. A promising strategy for restoring goal-directed control may be via boosting the salience of the outcome—for instance, by enhancing the link between the response and outcome.

Providing opportunities for performance tracking and administering other forms of performance-based feedback (e.g., primary and secondary rewards) have been used extensively in enhancing behavioral output [22,23]. For instance, the delivery of performance tracking information combined with a monetary reward successfully improved performance on a visual task [23]. A combination of primary and secondary rewards (e.g., juice and monetary incentives) has also been documented to improve goal-directed performance on a cued task-switching paradigm via motivational enhancement [24]. The promise of a future reward contingent on performance has sufficed in improving performance during task-switching, and accelerating responses during a reaction time task with congruent and incongruent stimuli [25,26]. Furthermore, trial-by-trial, transient monetary incentives (i.e., increasing reward magnitudes from low to high across trials) have served as salient performance boosters in tasks that taxed executive control, as well as visual perception [27]. Taken together with the finding that performance-contingent monetary rewards engage top-down control on task-switching [28], performance tracking and performance-contingent rewards may be prime candidates for enhancing goal-directed behavioral control. Thus, we propose that boosting motivation via performance-contingent feedback (e.g., intrinsic and extrinsic rewards that promote task performance improvements) may serve as a useful tool in restoring flexibility in otherwise rigid behaviors.

To achieve the goal of demonstrating and breaking a well-established habit, we introduce in Experiment 1 our novel Go/NoGo task that capitalizes on the familiar Green–Go, Red–NoGo associations people typically develop throughout the course of their lives. If the red–stop and green–go associations are well-learned, outcome-insensitive habits, there should be within-subject decrements in performance on an incongruent mapping of color to response (green–stop, red–go) compared to the well-learned congruent mapping (red–stop, green–go). That is, if participants are responding habitually, they should be more accurate when withholding responses to the red NoGo cue, and more likely to make errors of commission (e.g., responding to green cue when instructed to withhold responding), than if they are responding in a goal-directed manner. In comparison, there should be no such within-subject differences between novel color– response mappings (e.g. blue–stop, purple–go vs. purple–stop, blue–go). Then, in Experiments 2 and 3, we explore strategies to disrupt the well-learned red-stop, green-go habit by amplifying the salience of the action outcomes. Specifically, we use cumulative performance-contingent feedback to remediate the incongruency-related impairment—in an effort to restore goal-directed control in the face of habit-eliciting stimuli by reducing outcome-insensitive responses.

## Experiment 1

### Methods

#### Participants

We recruited 50 undergraduate students (32 female, 18 male participants; *M*_age_=20.28, *SD*_age_=2.96) from the Rutgers University-Newark campus for course credit. All subjects provided informed consent. Study protocols were approved by the Rutgers University Institutional Review Board. Participants were excluded if they reported having color-blindness.

#### Materials and Procedures

Participants were administered the Barratt Impulsivity Scale (BIS) [29], and randomly assigned to one of two stimulus type conditions (Familiar or Novel stimuli). They underwent a Go/NoGo task in which either Green and Red (Familiar condition) or Purple and Blue (Novel condition) traffic lights comprised Go and NoGo signals. Participants were instructed to respond as quickly and accurately to these stimuli as possible using the keyboard. A second phase followed in which the color-response contingencies were swapped (see Fig 1). Note that in the Familiar condition, the Green-Go/Red–NoGo mapping was considered “congruent” with associations in everyday life, while the Red–Go/Green–NoGo mapping was considered “incongruent.” We assumed that the Novel stimuli have no well-established Go or NoGo associations in daily life. The order in which participants underwent the two phases of the task was counterbalanced to ensure that the results could not be attributed to a specific order of managing the contingencies. Thus, we were able to examine the rigidity of our Familiar behavioral contingencies hypothesized to elicit outcome-insensitive responses in relation to a Novel stimulus set. An exit survey with demographic information concluded the study.

**Fig 1.**
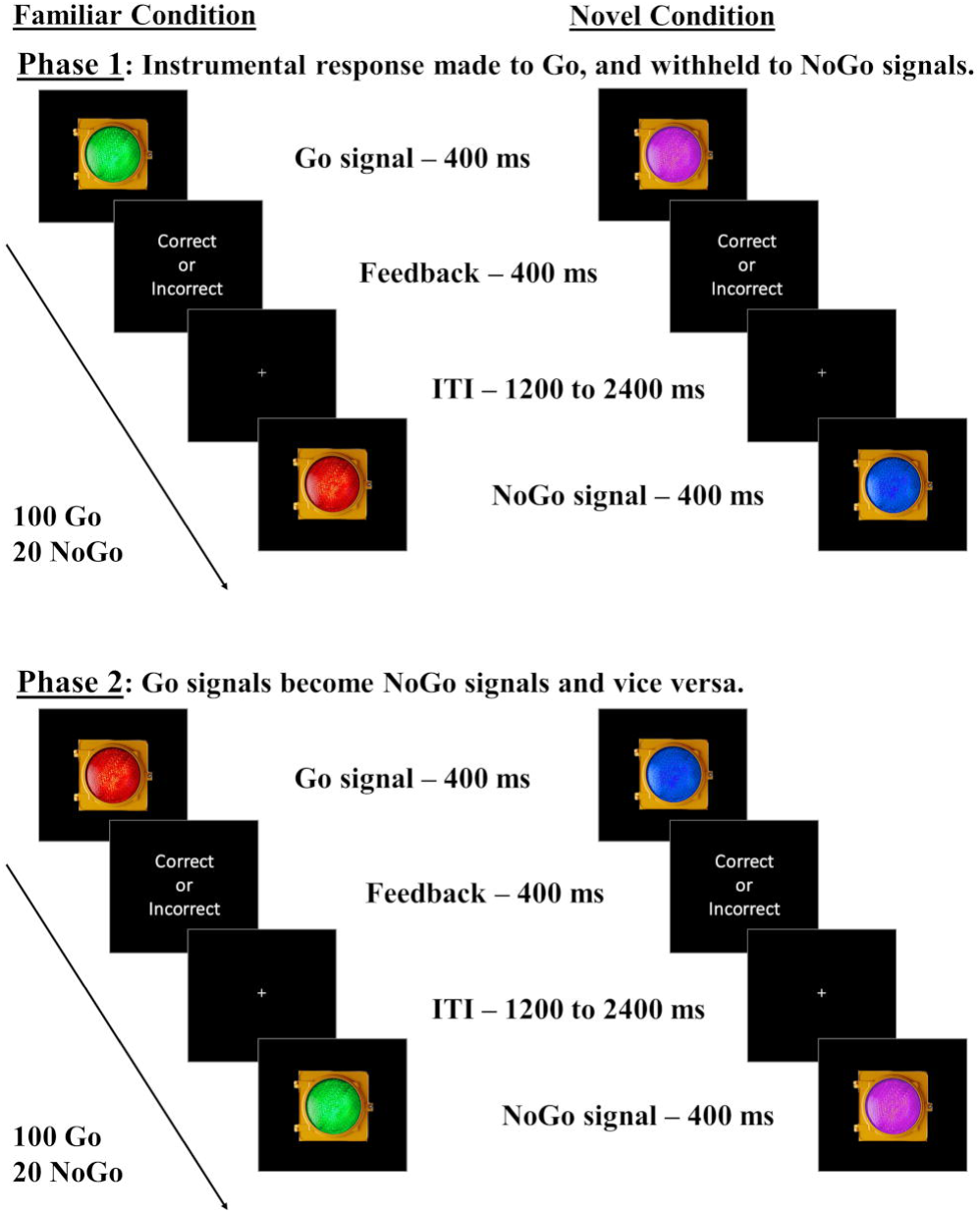
Go/NoGo task with familiar and novel lights. Participants are assigned to Familiar or Novel conditions. In the Familiar condition, subjects complete two phases: one where green signals Go and red signals NoGo (“congruent” mapping) and one where red signals Go and green signals NoGo (“incongruent” mapping). In the Novel condition, participants complete two similar phases, but the colors are blue and purple, for which there should be no strong pre-existing associations with “stop” and “go” responses. We predicted more commission errors in the Familiar condition for incongruent than congruent mappings, indicating outcome insensitivity, with no such within-subject differences expected in the Novel condition. Phase orders were counterbalanced across subjects.

Each phase comprised 100 Go and 20 NoGo trials (5:1 Go-NoGo ratio). The Go/NoGo stimuli remained onscreen for 400 milliseconds (ms), and each response produced a brief “correct” or “incorrect” text slide that offset after 400 ms (e.g., failure to withhold response in a NoGo trial produced the “incorrect” text slide). Go responses had to be performed before stimulus offset to be registered as correct by pressing the “1” key on the keyboard. The inter-trial intervals varied randomly between 1200 and 2400 ms to ensure engagement with the task. All participants completed a brief practice session (six correct Go or NoGo responses) using the same stimuli as the first phase. This practice session was conducted with the experimenter present to ensure the comprehension of instructions.

If these familiar associations elicit habitual, cue-driven behavioral control, subjects in the Familiar condition should experience a significant impairment in NoGo accuracy task demands when incongruent with lifelong experiences (Green–NoGo), compared to when they are congruent with lifelong experiences (Red-NoGo). In contrast, because the blue and purple stimuli are not expected to have strong Go or NoGo associations, participants in the Novel condition should show similar performance levels for both color–response mappings, illustrating the flexibility of responses executed towards the novel stimuli.

#### Data Analysis

Because the moderate ratio of Go to NoGo signals was expected to produce pre-potent Go responses [30], NoGo accuracy served as the primary measure of interest. As a secondary measure of outcome-sensitivity, identical analyses were performed using Go accuracy as dependent variable (DV). A mixed ANOVA with a DV of NoGo accuracy, Condition (Familiar or Novel stimulus conditions) as a between-subjects factor, and Mapping (congruent or incongruent mapping in the Familiar, and arbitrary color–response mapping in the Novel condition) as a within-subjects factor, was performed using Age, Gender, and Impulsivity (BIS score) as covariates. Post-hoc t-tests were employed to detect mapping-related differences in both conditions. We also performed a confirmatory omnibus test containing information from both conditions—a hierarchical multiple regression to test the predictive strength of the Condition variable on mapping-related impairment. We summarize these omnibus regression data below, but refer readers to the supplement for details (S1 and S2 Tables). Similar analyses were performed with Go response time (RT) as DV to further explore the data.

To determine sample size for our study, we performed an *a priori* power analysis using the effect size from an existing study examining Go/NoGo contingency change [31]. A within-group comparison of commission errors due to contingency change—one similar to the primary analyses reported above—determined that 12 participants would be needed per group to reach 80% statistical power. We adjusted this sample size in accordance with our two between-subjects factors that yielded four groups, (two Condition levels and two Order levels – that is, the counterbalanced orders in which participants completed the two phases of the task), warranting a sample size of 50.

### Results

#### Primary index of outcome-sensitivity: NoGo accuracy

To examine whether Condition (Familiar or Novel) predicted outcome-sensitivity, we performed a repeated measures ANOVA using NoGo accuracy as the DV, Condition as a between-subjects factor, Mapping as a within-subjects factor, controlling for Age, Gender, and Impulsivity as covariates. We found no main effect of Condition, *F*(1,45) = 0.99, *p* = .325, η_p_^2^ = .02, or Mapping, *F*(1,45) = 0.10, *p* = .748, η_p_^2^ < .01. but as evident in Fig 2, we found a significant Condition x Mapping interaction. *F*(1,45) = 8.65, *p* = .005, η_p_^2^ = .16. The congruent mapping produced higher accuracy compared to the incongruent mapping, which was not different from performance to the novel stimuli. Post-hoc paired-samples t-tests further revealed a significant difference in NoGo accuracy in the Familiar condition, *t*(24) = 3.53, *p* = .002, suggesting that the congruent “Red–NoGo” mapping elicits fewer errors of commission compared to the incongruent “Green-NoGo” mapping—a difference indicative of outcome-insensitive, habitual control. Contingency change yielded no differences in errors of commission between phases in the Novel condition, supporting the labile nature of newly learned associations, *t*(24) = −0.88, *p* = .387.

**Fig 2.**
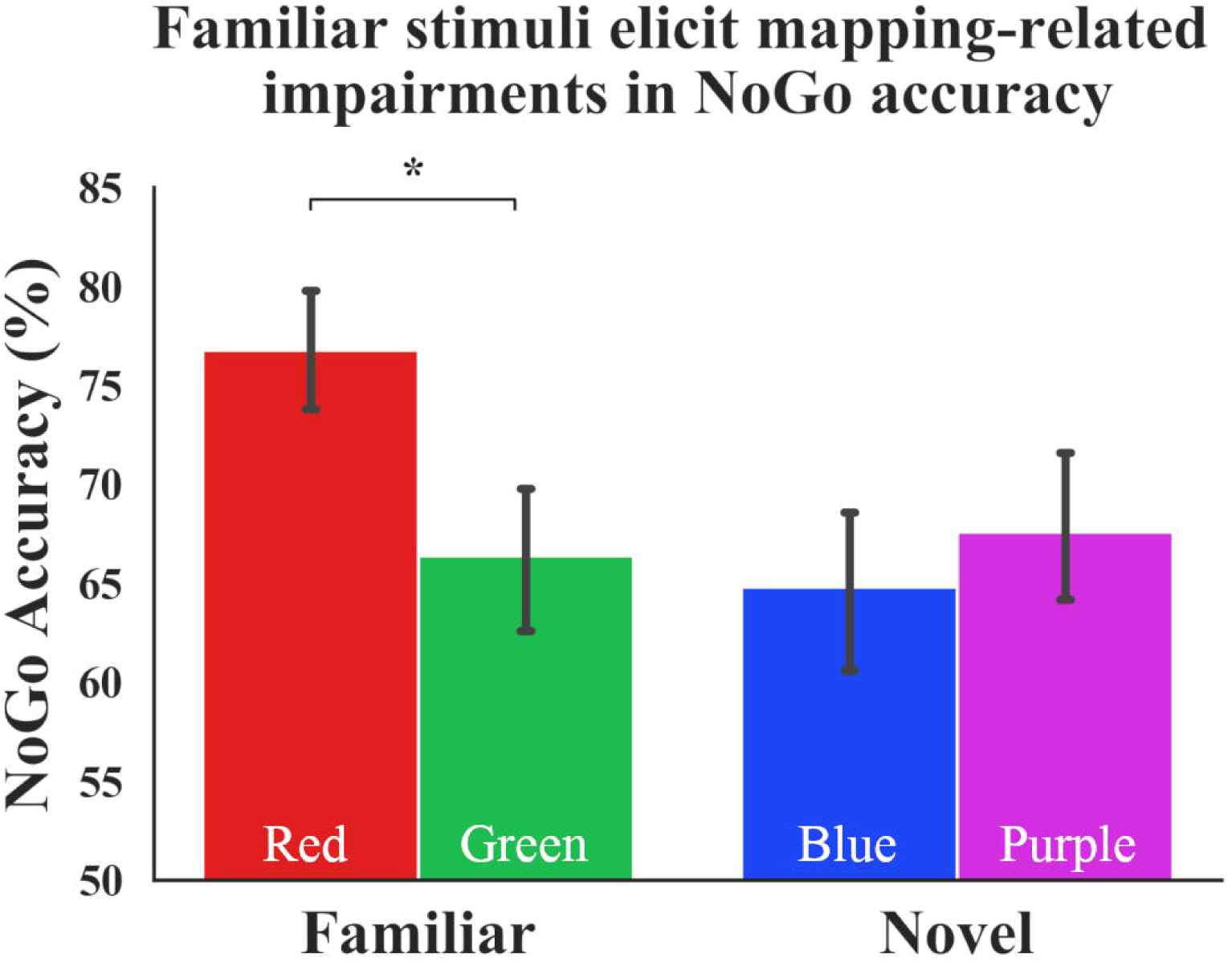
Familiar stimuli elicit mapping-related impairments in NoGo accuracy. The congruent mapping produces higher accuracy compared to the incongruent mapping, which is not different from performance to the novel stimuli. Specifically, participants make significantly fewer errors of commission when the NoGo signal is red compared to green. There is no difference in accuracy in the Novel condition when the NoGo signal is purple vs. blue. Condition x Mapping interaction: *p* = .005. Error bars depict standard error of mean (SEM). Color of bars reflects NoGo stimulus colors.

The omnibus regression test confirmed the significant effect of Condition. When controlling for participants’ age, gender, and self-reported impulsivity, the inclusion of the Condition regressor in the hierarchical multiple regression model explained an additional 15.5% of the variance in outcome-sensitivity: β_Condition_ = −0.40, *p* = .006, ΔR^2^ = .15, indicating differential outcome-sensitivity across Familiar and Novel conditions. The details of this omnibus regression test and beta weights of all model parameters can be found in the supplement (S1 Table).

#### Secondary index of outcome-sensitivity: Go accuracy

A mixed-design ANOVA controlling for Age, Gender, and Impulsivity as covariates, using Go accuracy as the DV revealed no main effect of Condition, *F*(1,45) = 0.19, *p* = .667, η_p_^2^ < .01, or Mapping, *F*(1,45) = 2.93, *p* = 0.094, η_p_^2^ = .06, but a near-significant Condition x Mapping interaction at *F*(1,44) = 3.93, *p* = .054, η_p_^2^ = .08 (Fig 3). Post-hoc paired-samples t-tests suggested a Go accuracy impairment only in the Familiar condition, *t*(24) = 3.10, *p* = .005, whereas no mapping-related Go impairment was observed in the Novel condition, *t*(24) = 0.28, *p* = .785.

**Fig 3.**
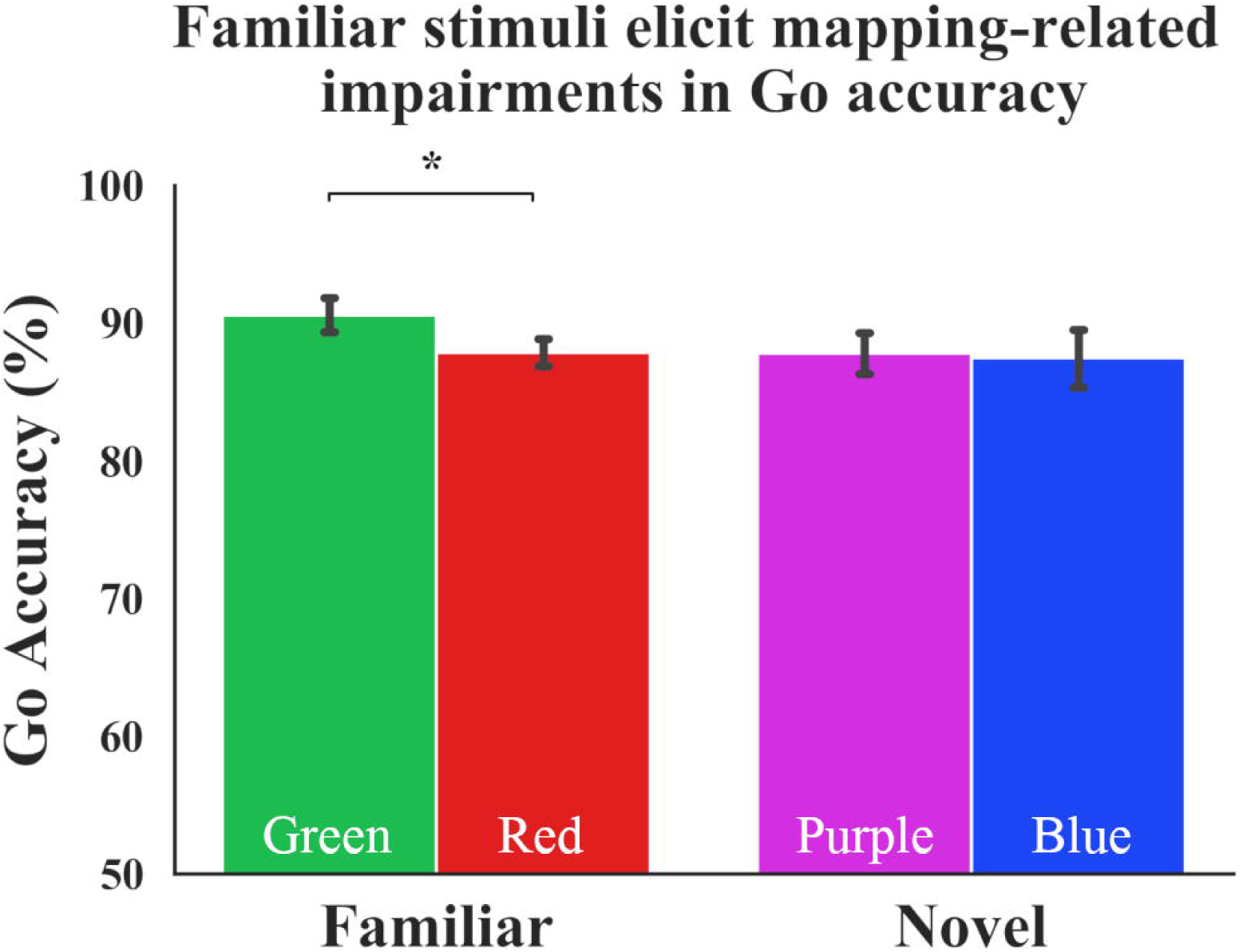
Familiar stimuli elicit mapping-related impairments in Go accuracy. Subjects perform worse when the Go signal is red compared to green. No such differences are seen in the Novel condition, when Go signal is blue vs. purple. Condition x Mapping interaction: *p* = .054. Error bars depict standard error of mean. Color of bars reflects Go stimulus colors.

The omnibus regression test also confirmed the role played by Condition on our secondary assay of outcome-sensitivity, Go accuracy. Controlling for participants’ age, gender, and impulsivity scores, the inclusion of the Condition regressor significantly predicted mapping-related Go accuracy changes: β_Condition_ = −0.27, ΔR^2^ = .07, *p* = .049 (see S2 Table for details). Lastly, similar analyses performed with Go RT as DV yielded no significant results (all *p*s > .05).

These Go accuracy data lend support to the hypothesis that while red and green stimuli are rigid and habitual in triggering stop/go actions, blue and purple stimuli are not strongly associated with either of the stop/go outcomes, but instead are labile and sensitive to the changes in action-outcome contingencies.

### Discussion

This experiment demonstrates that habitual behavior that capitalizes on existing, non-lab-derived associations, can be demonstrated in the lab. By using the strong links between the green–go and red–stop associations in a Go/NoGo task, we were able to quantify the degree of flexibility to well-stamped in cue–response–outcome associations. Importantly, our results suggest that responses are more outcome-insensitive (i.e., habitual) when the stimulus meanings are congruent with our experiences with traffic lights in daily life (i.e., when a traffic light indicating “stop” is red, rather than green, blue or purple). We note that incongruency-related impairments alone are not enough to conclude that a response is habitual; rather this conclusion must be verified by a comparison of the habitual associations (i.e., green–go, red–stop) with the novel control condition Go/NoGo associations (i.e., purple–go, blue–stop). Specifically, these red and green light stimuli triggered outcome-insensitive actions as evidenced by an accuracy impairment when Go and NoGo contingencies were incongruent with their well-established meanings outside of the lab. In contrast, the novel purple–go and blue–stop contingencies are not well-established in one’s daily experiences, and their associative strength is limited to the participant’s brief experience in the lab. Therefore, compared to the familiar stimuli, the actions evoked by the novel stimuli are more flexible to contingency changes, as reflected by similar NoGo and Go accuracy scores for blue vs. purple.

It can be argued that if these familiar red and green stimuli elicit outcome-insensitive habits, one should also display lower accuracy rates to green-NoGo compared to blue or purple-NoGo contingencies. However, our results above suggest that green-NoGo performance is similar to those elicited by the novel stimuli. The comparable performance here may be due to between-subject designs requiring more power compared to within-subject designs [36], thus making it more difficult to detect a potential decrement in green-NoGo accuracy. We further examine the unexpected pattern observed here in Experiments 2 and 3, with the prediction that with sufficient power, the green-NoGo mapping will indeed elicit lower accuracy rates compared to either Novel condition NoGo mapping.

Assessing motivational control, which attributes the source of one’s actions to either a preceding cue or its consequences, has long relied on experimental manipulations of outcome value. Rodent and human studies employing outcome-devaluation procedures of food rewards have depended on the subjects’ comparable palatability of the foods used in the research, as well as the development of an outcome-insensitive habit via over-training of these action-outcome contingencies [11,32]. Other researchers have made use of the instructed devaluation of outcomes, and computational investigations of choice strategy categorizations of model-based and model-free performance [10]. Although tremendously effective in their own avenues, a common area outside of the reach of these tasks is well-learned habits that better represent real world scenarios.

Our Go/NoGo task with familiar and novel stimuli provides new possibilities in studying habits. We demonstrate habits in a lab setting using stimuli that do not require lengthy training sessions to develop strong stimulus–response associations. This time- and cost-effective paradigm can serve as an especially useful tool in studying habits in expensive neuroimaging contexts. Perhaps more importantly, taking advantage of well stamped-in cue–response associations to study habits promises to contribute to translational science via new research avenues. For instance, although contemporary paradigms have proved fruitful in studying the formation and expression of habits, the nature of the tasks do not facilitate the investigation of habit disruption. Novel associations that have become outcome-insensitive following limited, lab-specific experience may not be rigid enough to represent real-world behaviors, and breaking these weak habits may not be translationally valuable.

## Experiment 2

We attempt the breaking of well-learned habits in Experiment 2, in which we boost motivation via cumulative performance feedback prior to contingency reversal. Because the motivational control framework attributes habits to be driven by antecedent cues and goal-directed actions to be guided by resulting outcomes, we hypothesized that amplifying the salience of the outcome may promote goal-directed performance at the expense of habitual control, thus aiding in breaking the well-learned habit.

### Methods

#### Participants

We recruited 100 undergraduate students (67 female and 33 male participants; *M*_Age_=20.26, *SD*_Age_=3.05) from the Rutgers University-Newark campus. All participants provided informed consent and received course credit for their participation. Study protocols were approved by the Rutgers University Institutional Review Board. Participants were excluded if they reported having color-blindness.

#### Procedures

For the Go/NoGo task, participants were randomly assigned to a Feedback Group or No Feedback Group, and within each group, participants were randomly assigned to either Novel or Familiar condition, as in Experiment 1.

Feedback Group: After completing the BIS, participants underwent a similar Go/NoGo task to the one described in Experiment 1. Accordingly, each phase comprised 100 Go and 20 NoGo trials (5:1 Go–NoGo ratio). As reported in Experiment 1, all stimuli remained on the screen for 400 ms, and responses produced brief feedback slides consisting of “correct” or “incorrect” that offset after 400 ms (e.g., failure to withhold response in a NoGo trial produced the “incorrect” text slide). Go responses had to be performed before stimulus offset to be registered as correct by pressing the “1” key on the keyboard. The inter-trial intervals varied randomly between 1200 and 2400 ms to ensure engagement with the task. All subjects completed a brief practice session (six correct Go or NoGo responses) using the same stimuli that comprised the task. This practice session was conducted with the experimenter present to ensure the comprehension of instructions.

In the Familiar condition, participants were instructed to “Go” on green traffic light stimuli as quickly and accurately as possible, and withhold responses to the red traffic light. Next, a cumulative performance feedback manipulation followed, in which we displayed subjects’ percent NoGo accuracy scores on the screen. Participants were informed that the percentage score reflected their performance thus far (they were not informed that the score only reflected NoGo accuracy), and in the next phase of the task, the Go and NoGo signals would be reversed, such that they would need to make a response as quickly and accurately as possible to the red traffic light, and refrain from responding to the green traffic light. Identical feedback and task instructions were provided to the participants in the Novel condition regarding the change in contingencies of the purple–Go and blue–NoGo associations. It should be noted that Experiment 1 reports differential mapping-related impairments across Familiar and Novel conditions regardless of the order in which phases were completed (S1-S2 Tables). Therefore, unlike Experiment 1, the phase orders in Experiment 2 were not counterbalanced, in that all participants in the Familiar condition underwent the congruent (Green–Go, Red–NoGo) mappings first, followed by the incongruent mappings; all participants in the Novel condition underwent the Purple–Go, Blue–NoGo mapping first, and these mappings were reversed in the second phase. This change in experimental protocol enabled rendering the congruent contingency as baseline for participants in the Familiar group, and testing whether the presence of a mid-experiment performance manipulation affected subsequent incongruent task performance. An exit survey consisting of demographic questions concluded the experiment.

No Feedback Group: Participants in the No Feedback group underwent the same procedures as the Feedback group, except that no cumulative performance feedback was provided at any point. This No Feedback group served as a control condition for the Feedback group, as well as an internal replication of Experiment 1.

#### Data Analysis

To examine the role of Feedback, mixed-design ANOVAs with NoGo accuracy as DV, Feedback as a between-subjects and Mapping as a within-subjects factor were performed for each Condition, using the controlled variables Age, Gender, and Impulsivity as covariates. Post-hoc t-tests were carried out to examine mapping-related accuracy differences in both Feedback groups. As a secondary measure of outcome-sensitivity, identical analyses were performed using Go accuracy as a DV. Similar analyses were performed with Go RT as DV to further explore the data. It should be noted that we did not test for a three-way Condition x Feedback x Mapping interaction with any of our DVs, because our primary interest was determining whether cumulative performance feedback has any effect on motivational control, not necessarily whether this effect differs based on the familiarity of the stimuli. For example, we would not expect cumulative feedback to promote accuracy improvements in the Familiar Condition while impairing performance in the Novel condition.

Building from Experiment 1, we performed a confirmatory omnibus hierarchical multiple regression to test the predictive strength of the Condition and Feedback variables on mapping-related impairment. The summary of the omnibus regression test is reported below, and its details can be found in the supplement (S3-S4 Tables).

We performed a power analysis using the effect size of the Condition x Mapping interaction in Experiment 1 (η_p_^2^ = .16) and determined that a sample of 12 participants per group would be sufficient to reach 80% statistical power to detect the effect of differential accuracy rates due to Condition. We opted for this interaction value for our investigation of the role of feedback, because we wanted our feedback-related assertions to be grounded in predictions of a replicated effect of habitual performance to familiar, and goal-directed performance to novel stimuli. To further increase statistical power due to the addition of a Feedback group per condition, we increased our sample size to 25 per group—a total of 100 undergraduate students.

### Results

#### Primary index of outcome-sensitivity: NoGo accuracy

We hypothesized that performance feedback may be a salient factor that can potentially restore goal-directed control when managing these well-established associations. However, cumulative performance feedback did not break the habits elicited by these familiar stimuli. We performed a mixed-design ANOVA using NoGo accuracy as the DV, and Age, Gender, and Impulsivity as covariates. We found no main effect of Feedback, *F*(1,45) = 0.08, *p* = .778, η_p_^2^ <.01, or Mapping, *F*(1,45) = 1.96, *p* = .169, η_p_^2^ = .04, and we also found that no significant Feedback x Mapping interaction exists : *F*(1,45) = 0.08, *p* = .776, η_p_^2^ < .01 (see Fig 4). Post-hoc t-tests revealed significant incongruency-related impairments in both Feedback, *t*(24) = 2.72, *p* = .012, and No Feedback, *t*(24) = 3.16, *p* = .004, groups, indicating that cumulative performance feedback did not prevent habitual control from dominating in the Familiar condition. Although we were unable to break habits as hypothesized here, our findings lend support to the rigidity of these well-learned associations that persevere in the face of an otherwise salient motivational manipulation, performance feedback [33,34].

**Fig 4.**
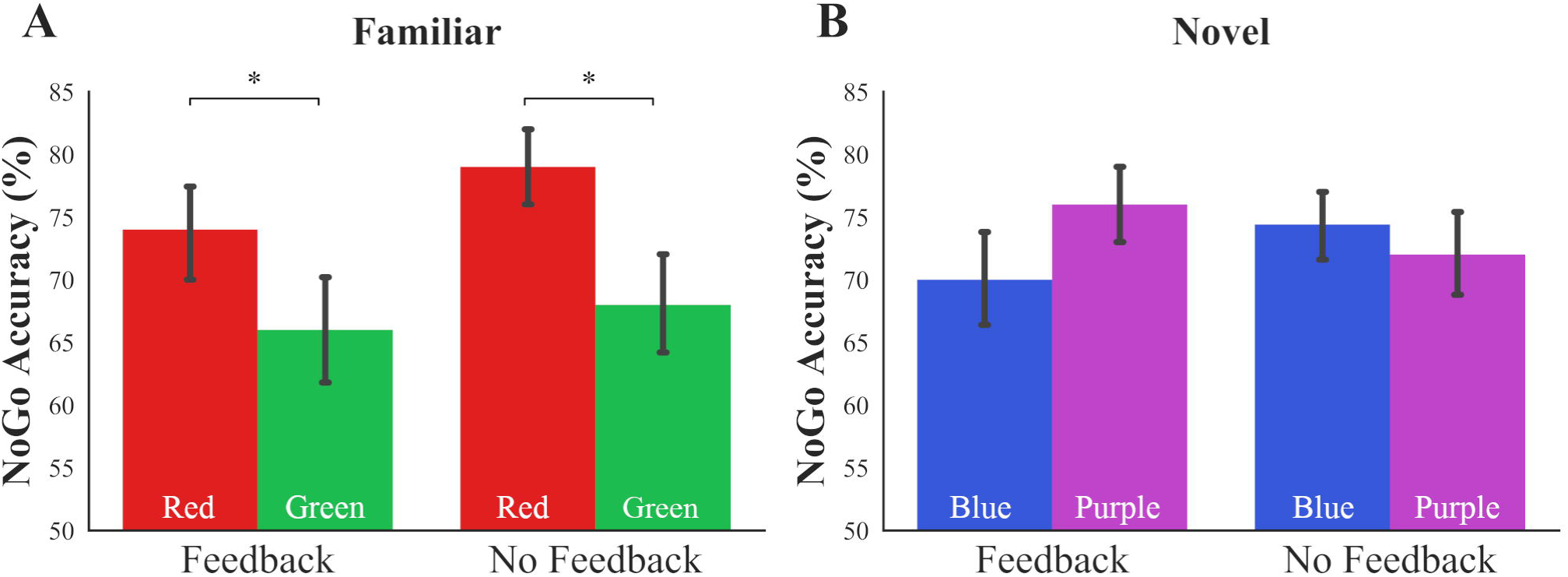
Performance feedback does not significantly disrupt well-established habits. (A) In the Familiar condition, both Feedback and No Feedback groups suffer an incongruency-related impairment (*p* = .776) in NoGo accuracy. (B) NoGo accuracy in the Novel condition is not significantly improved by performance feedback (sig. interaction of *p* = .033, non-sig. post-hoc t-tests: *p* > .05). Error bars denote SEM. Color of bars reflects NoGo stimulus colors.

We performed a similar ANOVA to determine whether cumulative performance tracking improved goal-directed control of novel associations. As seen in Fig 4, we did not find a main effect of Feedback, *F*(1,45) = 0.40, *p* = .528, η_p_^2^ < .01, or Mapping, *F*(1,45) = 0.60, *p* = .442, η_p_^2^ = .01, yet found a Feedback x Mapping interaction on NoGo accuracy in the Novel Condition when controlling for Age, Gender, and Impulsivity as covariates: *F*(1,45) = 4.84, *p* = .033, η_p_^2^ = .10. In sum, these results suggest that performance feedback alone may not be a salient enough manipulation to restore goal-directed control.

#### Secondary index of outcome-sensitivity: Go accuracy

We performed a mixed-design ANOVA of the Familiar condition data using Go accuracy as DV, Feedback as a between-, and Mapping as a within-subjects factor, with Age, Gender, and Impulsivity as covariates. We found no significant main effect of Feedback *F*(1,45) = 0.10, *p* = .751, η_p_^2^ < .01, or Mapping, *F*(1,45) = 0.14, *p* = .705, η_p_^2^ < .01, but found a significant Feedback x Mapping interaction: *F*(1,45) = 4.73, *p* = .035, η_p_^2^ = .09 (Fig 5), suggesting that Go accuracy was affected differentially by performance feedback. Post-hoc paired-samples t-tests of Go accuracy across phases yielded evidence for an incongruency-related impairment in the No-Feedback group, *t*(24) = 3.22, *p* = .004), but not in the Feedback group, *t*(24) = 1.14, *p* = .265. Indeed, the omnibus hierarchical regression model attributes Condition and Feedback regressors a significant role in predicting Go accuracy change (β_Condition_ = −0.32, *p* = .001, β_Feedback_ = 0.28, *p* = .003; ΔR^2^ = .18).

**Fig 5.**
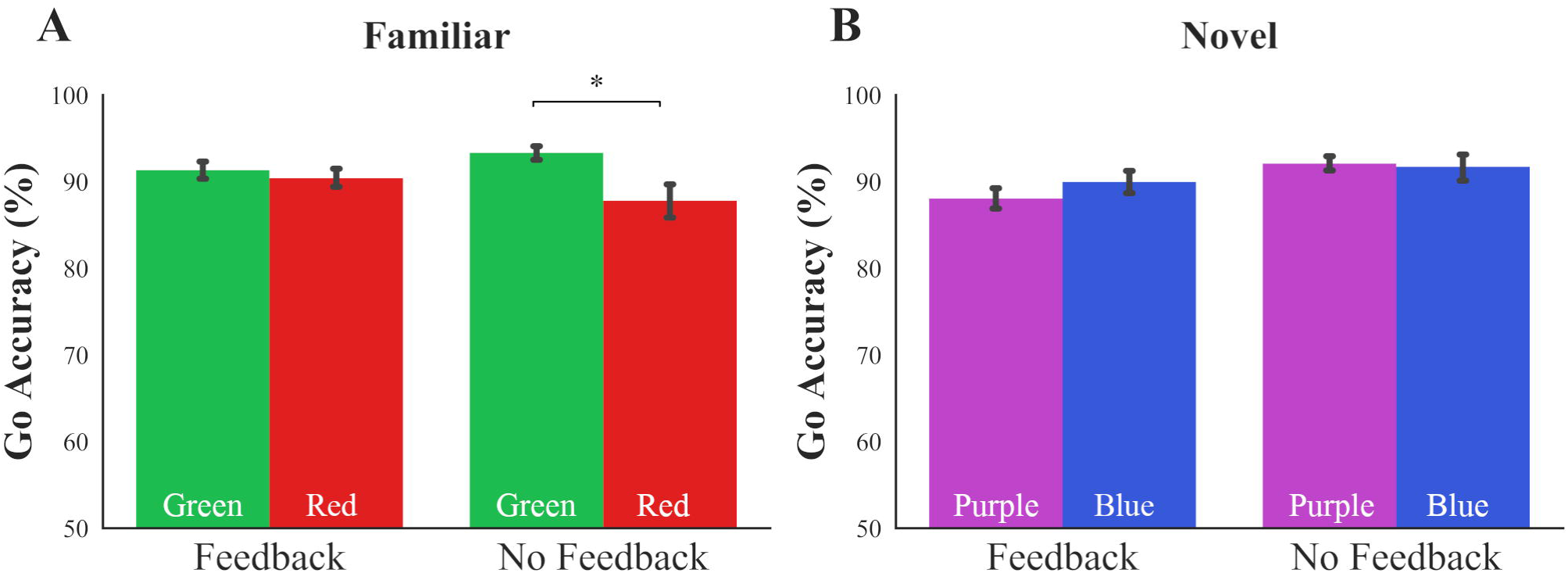
Performance feedback protects against habitual Go actions. (A) When participants received cumulative feedback on their performance, the Go accuracy impairment otherwise observed without feedback was prevented when managing Familiar stimuli (Feedback x Mapping interaction *p* = .035). (B) Performance feedback did not significantly improve Go accuracy in the Novel condition (Feedback x Mapping interaction *p* = .117). Error bars denote SEM. Color of bars reflects Go stimulus colors.

Despite the significant Feedback regressor in the omnibus test, we did not observe a significant improvement effect due to cumulative performance feedback in the Novel condition Go accuracy results. A mixed-design ANOVA using Go accuracy as the DV, Feedback as the between-, and Mapping as the within-subjects factor, with Age, Gender, and Impulsivity as covariates revealed no significant main effect of Feedback, *F*(1,45) = 3.53, *p* = .067, η_p_^2^ = .07, or Mapping, *F*(1,45) = 3.14, *p* = .083, η_p_^2^ = .06, and no significant Feedback x Mapping interaction: *F*(1,45) = 2.56, *p* = .117, η_p_^2^ = .05 (Fig 5). Post-hoc paired-samples t-tests suggest an improvement effect only in the Feedback group: *t*(24) = −2.39, *p* = .025 with feedback, *t*(24) = 0.32, *p* = .749 without feedback. Given the lack of significant Feedback x Mapping interaction in the Novel condition, we refrain from speculating further about the effect of cumulative performance feedback on goal-directed Go responses. Similar analyses performed with Go RT as DV yielded no significant findings (all *p*s > .05).

### Discussion

In sum, we report that cumulative performance feedback is not sufficient to disrupt the well-learned habits elicited by the familiar stimuli used in our task. However, supplementary analyses using accessory measures of behavioral control (i.e., familiar Go accuracy), suggest that feedback may be a useful tool in enhancing behavioral flexibility. Therefore, these patterns warrant further examination of feedback to disrupt habitual control.

We conclude that cumulative performance feedback was not salient enough to break habits according to our primary analyses, yet our findings were valuable in two ways. First, the validity of our Go/NoGo task using well-learned associations to study habits relies on the rigidity of these green–go and red–stop associations. The persistent habitual control exhibited here despite the delivery of performance feedback lends credence to the associative strength of our familiar stimuli. Next, given the modest signs of performance improvement due to the presentation of performance information, early reports of combined (i.e., performance tracking and monetary incentives) feedback’s positive effects on performance, and the beneficial effects of performance-contingent feedback on behavioral flexibility [23–27], we were motivated to enhance the salience of the provided feedback to break well-learned habits. In Experiment 3, we further amplified the salience of the outcome by pairing performance-contingent cumulative feedback with a bonus monetary reward prior to changing Go and NoGo contingencies. We studied the effects of monetary and cumulative performance feedback on Go/NoGo task performance, and whether this amplification of outcome salience resulted in the breaking of a well-learned habit, and improvement of novel, goal-directed performance.

## Experiment 3

The promising but insufficient effect of cumulative performance feedback on the motivational control of action motivated us to examine the combined effect of performance and monetary input. Thus, we implemented in our mid-experiment performance feedback manipulation a cash bonus. We hypothesized that this bonus, combined with performance tracking information, would enhance goal salience and promote cognitive control processes to override habitual control. Experimental procedures were identical to those described in Experiment 2, with the addition of awarding participants in the Feedback group a surprise $5 cash bonus before the change in Go/NoGo mappings.

### Methods

#### Participants

To test the effects of dual feedback, we recruited the same number of participants for Experiment 3 as in Experiment 1. One-hundred participants (76 female, 24 male participants; *M*_age_=19.74, *SD*_age_=2.79) from the Rutgers University-Newark undergraduate research subject pool were recruited for course credit. All participants provided informed consent. Study protocols were approved by the Rutgers University Institutional Review Board. Participants were excluded if they reported having color-blindness.

#### Procedures

After completing BIS, participants underwent a similar Go/NoGo task to the one described in Experiment 2, where they were randomly assigned to Feedback and No Feedback groups, and Familiar and Novel conditions. As in Experiment 2, each phase comprised 100 Go and 20 NoGo trials (5:1 Go–NoGo ratio), and the stimuli remained on the screen for 400 ms. Go and NoGo responses (or lack thereof) produced brief feedback slides consisting of “correct” or “incorrect” that offset after 400 ms (e.g., failure to withhold response in a NoGo trial produced the “incorrect” text slide). Go responses had to be performed before stimulus offset to be registered as correct by pressing the “1” key on the keyboard. The inter-trial intervals varied randomly between 1200 and 2400 ms to ensure engagement with the task. All participants completed a brief practice session prior to the task, similar to the previous two experiments.

Identical to Experiment 2, in the Familiar condition’s first phase, participants were instructed to “Go” on green traffic light stimuli as quickly and accurately as possible, and “NoGo” on red traffic light stimuli. Next, a monetary and cumulative performance feedback manipulation followed, in which we displayed participants’ cumulative NoGo accuracy as a percentage score on the screen. Participants were informed that the percentage score reflected their performance thus far. Additionally, unique to Experiment 3, the experimenter left the room, and returned briefly after with a $5 bill, and informed the participant that this money was earned because of performance thus far in the task. Unbeknownst to the participants, the cash bonus was not actually contingent on performance. The participant was then informed that the Go and NoGo signals would be reversed, such that they would need to make a response as quickly and accurately as possible to the red traffic light, and refrain from responding to the green traffic light. Identical performance and monetary feedback information and reversal instructions were provided to the participants in the Novel condition regarding the reversal of purple–Go and blue– NoGo responses. An exit survey containing demographic questions concluded the experiment.

Participants in the No Feedback group underwent the same procedures as the Feedback group, except for the feedback manipulation, in that participants received no cumulative performance or monetary feedback.

#### Data Analysis

To reveal the potential effect of dual feedback on motivational control, we performed mixed-design ANOVAs with NoGo accuracy as the DV, Feedback as a between- and Mapping as a within-subjects factor for each Condition, using the Age, Gender, and Impulsivity variables as covariates. Post-hoc paired-samples t-tests were carried out when necessary to examine mapping-related accuracy differences in both Feedback groups. As a supplemental measure of outcome-sensitivity, identical tests were performed using Go accuracy as the DV. Similar analyses were performed with Go RT as DV to further explore the data. Identical to Experiment 2, we performed a confirmatory omnibus hierarchical multiple regression to test the predictive strength of the Condition and Feedback variables on outcome-sensitivity. The summary of the omnibus regression test are reported below, and the details can be found in the supplement (S5 and S6 Tables). Lastly, to further explore whether green-NoGo (i.e., the color-response mapping that is incongruent with daily experiences) elicits lower accuracy rates compared to either Novel color-response mapping with sufficient power, we pooled Experiment 2 and 3 data (due to their identical No-Feedback procedures) and performed independent-samples t-tests to compare green-NoGo accuracy to purple- and blue-NoGo accuracy in the No-Feedback conditions.

### Results

#### Primary index of outcome-sensitivity: NoGo accuracy

We tested the role of dual feedback in disrupting habitual control to familiar stimuli by performing a mixed-design repeated measures ANOVA on data from the Familiar condition, using NoGo accuracy as the DV. We found no main effect of Feedback, *F*(1,45) = 0.75, *p* = .390, η_p_^2^ = .10, or Mapping, *F*(1,45) = 1.51, *p* = .225, η_p_^2^ = .03, but found a significant Feedback x Mapping interaction when controlling for Age, Gender, and Impulsivity: *F*(1,45) = 5.24, *p* = .027, η_p_^2^ = .10 (see Fig 6). This interaction suggests differential impairment based on the availability of cumulative performance and monetary feedback, such that the lack of feedback when managing familiar stimuli resulted in a significantly larger incongruency-related decrement in NoGo accuracy. Post-hoc t-tests confirmed a significant impairment in the No-Feedback group, *t*(24) = 5.25, *p* < .001, replicating our findings from Experiments 1 and 2, but no significant effect in the Feedback group *t*(24) = 1.92, *p* = .067.

**Fig 6.**
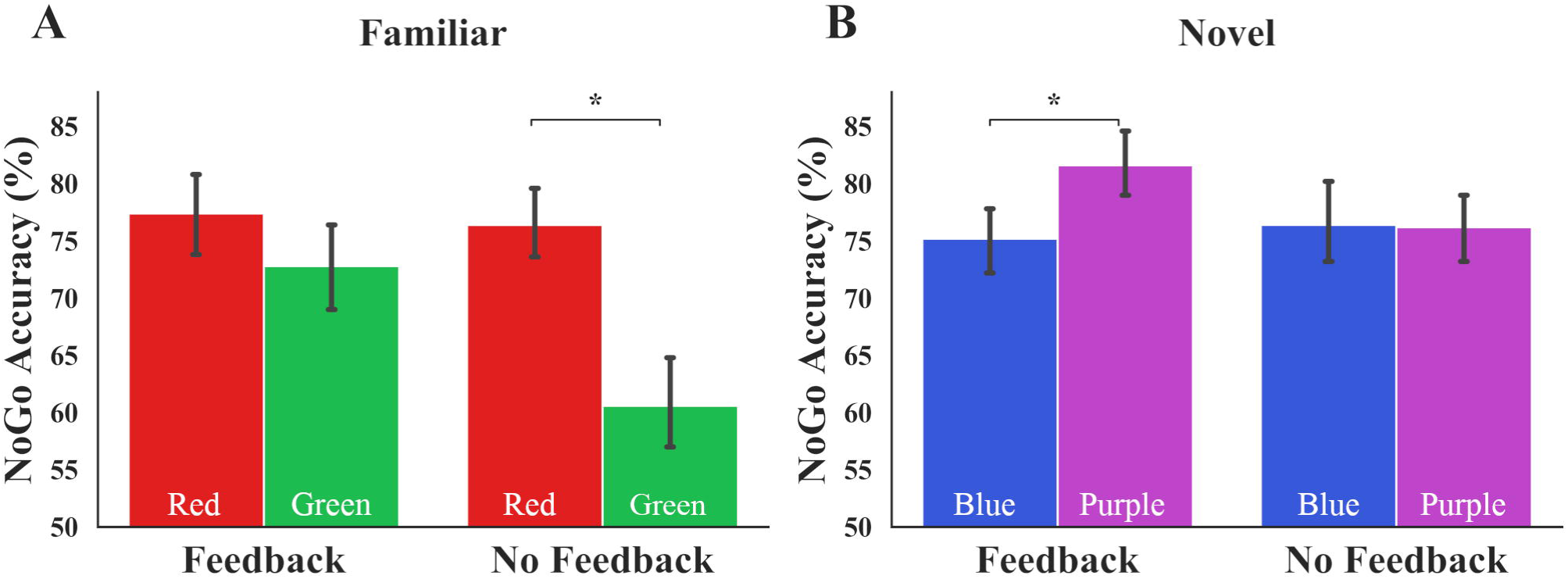
Monetary and performance feedback disrupt habits while improving goal-directed performance to newly-learned stimuli. (A) Providing performance and monetary feedback prevents the incongruency-related impairment normally indicative of habitual control (Feedback x Mapping interaction: *p* = .027). (B) Dual feedback also improves goal-directed control of novel associations significantly (Feedback x Mapping interaction: *p* = .038). Error bars denote SEM. Color of bars reflects NoGo stimulus colors.

To understand whether dual feedback enhanced goal-directed performance to newly-learned associations, we performed similar analyses on the Novel condition data. The mixed-design ANOVA, when controlling for Age, Gender, and Impulsivity as covariates, yielded no main effect of Feedback, *F*(1,45) = 0.10, *p* = .756, η_p_^2^ < .01, or Mapping, *F*(1,45) = 0.42, *p* = .522, η_p_^2^ = .01; however, we found a significant Feedback x Mapping interaction on NoGo accuracy in the Novel condition: *F*(1,45) = 4.55, *p* = .038, η_p_^2^ = .09 (Fig 6). Post-hoc t-tests revealed significant improvement of NoGo accuracy in the Feedback group, *t*(24) = −2.32, *p* = .029, which was not observed in the No-Feedback group, *t*(24) = 0.08, *p* = .938.

Consistent with these significant Feedback x Mapping interactions in both Familiar and Novel conditions, our omnibus hierarchical regression model revealed Condition and Feedback regressors to be significant predictors of outcome-sensitivity. Combined, Condition and Feedback explained 26.6% of the variance in mapping-related NoGo accuracy change (β_Condition_ = −0.43, *p* < .001, β_Feedback_ = 0.28, *p* = .003; ΔR^2^ = .27). These data suggest that the differential mapping-related NoGo impairment observed in Experiment 2 was replicated in Experiment 3, and importantly, that dual feedback is able to significantly predict improvements in performance. The entirety of the omnibus test can be found in the supplement (S5 Table).

#### Secondary index of outcome-sensitivity: Go accuracy

As a supplementary assay of behavioral control, we analyzed Go accuracy using similar statistical procedures. We input Go accuracy as a DV, Feedback as a between-, and Mapping as a within-subjects factor, with Age, Gender, and Impulsivity as covariates into a mixed-design ANOVA. For the Familiar condition, we found no significant main effect of Feedback *F*(1,45) = 2.36, *p* = .131, η_p_^2^ = .05, a significant main effect of Mapping, *F*(1,45) = 4.15, *p* = .048, η_p_^2^ = .08, but no significant Feedback x Mapping interaction: *F*(1,45) = 2.52, *p* = .119, η_p_^2^ = .05 (Fig 7), suggesting that Go accuracy was not significantly affected by dual feedback in the Familiar condition. However, post-hoc paired-samples t-tests revealed incongruency-related impairments in Go actions specific to the No Feedback group: *t*(24) = 2.58, *p* = .017 without feedback vs. *t*(24) = 0.10, *p* = .925 with dual feedback. Given the lack of interaction, we refrain from asserting that dual feedback disrupts habitual Go actions—our secondary assay of outcome-sensitivity.

**Fig 7.**
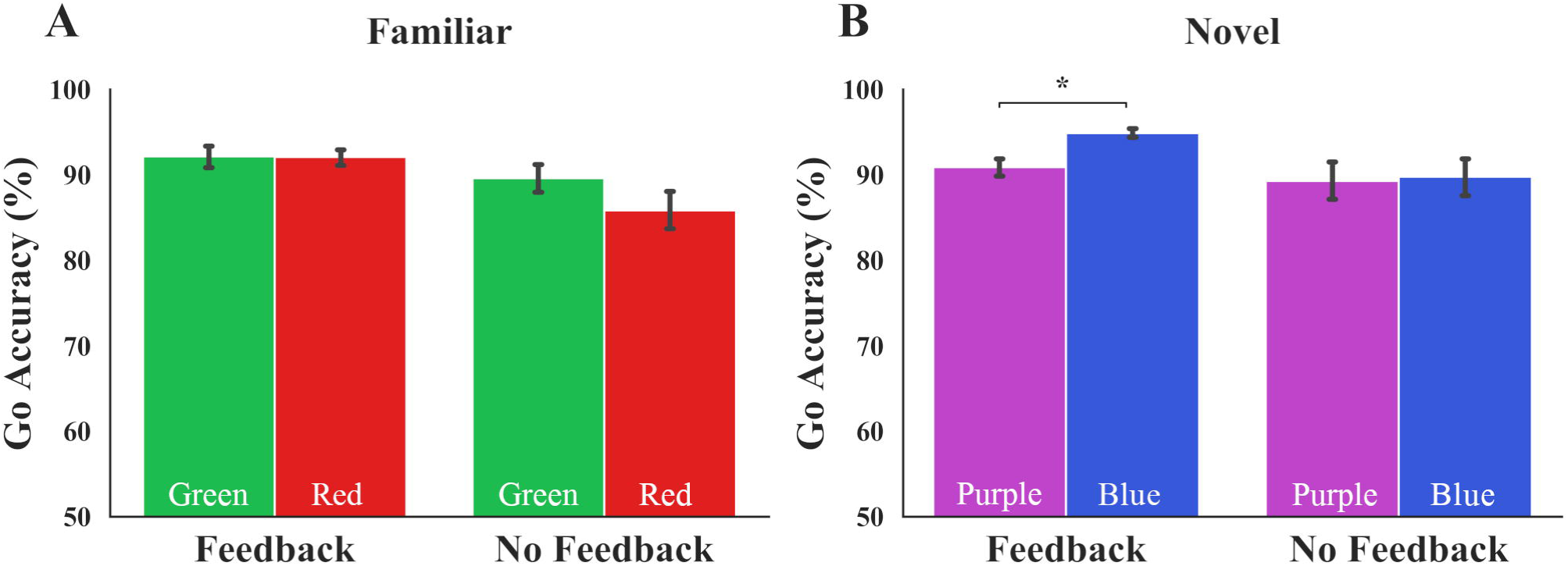
Dual feedback improves goal-directed Go accuracy. (A) Dual feedback did not have a significant effect on the incongruency-related Go accuracy impairment when managing well-learned cues (*p* = .119). (B) Dual feedback improved goal-directed Go responses to novel associations (*p* = .012). Error bars denote SEM. Color of bars reflects Go stimulus colors.

We then tested the effect of dual feedback on Go accuracy in the Novel condition to determine whether our enhanced feedback manipulation improved goal-directed control when managing the contingency changes in newly-learned associations. We performed a mixed-design repeated measures ANOVA using Go accuracy as the DV, Feedback as the between-, and Mapping as the within-subjects factor, with Age, Gender, and Impulsivity as covariates. This ANOVA yielded a significant main effect of Feedback, *F*(1,45) = 5.49, *p* = .024, η_p_^2^ = .11, and no significant effect of Mapping, *F*(1,45) = 0.49, *p* = .488, η_p_^2^ = .01; however, it revealed a significant Feedback x Mapping interaction: *F*(1,45) = 6.93, *p* = .012, η_p_^2^ = .13 (see Fig 7). Post-hoc t-tests of each Feedback group confirms that monetary incentives paired with cumulative performance feedback significantly improved newly-learned Go associations that are executed by the goal-directed system: *t*(24) = −4.86, *p* < .001 with dual feedback, *t*(24) = −0.51, *p* = .616 with no feedback.

Our omnibus hierarchical regression model reveals that Condition and Feedback regressors significantly predict mapping-related Go accuracy changes. These regressors in sum account for 21% of the variance in the DV (β_Condition_ = −.36, *p* < .001, β_Feedback_ = .28, *p* = .004; ΔR^2^ = .21). These values suggest that Go accuracy is selectively impaired in the Familiar condition, and Feedback is able to promote goal-directed Go actions. Due to the non-significant Condition x Mapping interaction in the Familiar condition data, we restrict the scope of our dual feedback assertions on Go accuracy to the Novel condition. Details of the omnibus regression can be found in the supplement (S6 Table). Lastly, similar analyses performed with Go RT as DV yielded no significant findings (all *p*s > .05).

Finally, when we combine No-Feedback groups in Experiments 2 and 3 where participants undergo identical procedures, we find that the green-NoGo mapping (*M*_Green_ = 64.30, *SD*_Green_ = 20.35) yields significantly lower accuracy rates than either novel stimulus (*M*_Blue_ = 75.40, *SD*_Blue_ = 15.87, *M*_Purple_ = 74.10, *SD*_Purple_ = 16.03) despite the between-subjects design (green vs. blue: *t*(98) = 3.04, *p* = .003; green vs. purple: *t*(98) = 2.67, *p* = .009). This result suggests that with sufficient power, we are able to detect that the incongruent color-response mapping yields impaired performance in comparison to the newly-learned color-response contingencies.

### Discussion

Collectively, our Experiment 3 findings suggest that a global motivational boost involving amplified performance and monetary feedback produces a habit-breaking effect that restores goal-directed control. Without feedback, we observe a significant impairment in NoGo and Go accuracy when familiar green and red light stimuli demand responses incongruent with daily experiences. We find that this outcome-insensitive habit (i.e., inflexible, cue-driven behavior that persists despite the outcome) of the green–go and red–stop actions is disrupted when participants are provided dual feedback, such that the significant incongruency-related NoGo impairment otherwise seen without feedback is prevented. Moreover, our dual feedback manipulation also improves goal-directed control when managing newly-learned associations, as evidenced by significant enhancements to NoGo and Go performance in the Novel group. Possibly, cumulative performance feedback may be enhancing intrinsic motivation. The percentage score may provide individuals the opportunity to track task performance improvements, potentially boosting motivation to improve task-competence [35]. Paired with the extrinsic reward of a monetary bonus, the dual feedback provided in our experiment may be producing a global increase in motivation, resulting in more deliberate control of otherwise inflexible behaviors.

Importantly, the beneficial effect of such feedback generalizes to more flexible goal-directed performance, as we observe a significant improvement in NoGo and Go accuracy scores to novel blue–go and purple–stop contingencies when participants are provided dual feedback. Without feedback, we find no mapping-related difference in accuracy to novel stimuli, serving as support for the flexible nature of these newly-learned associations that can readily be reassigned per changes in one’s environment. These findings identify dual feedback as a powerful predictor of motivational control enhancement.

### General Discussion

In a three-experiment study, we introduce a novel Go/NoGo task that capitalizes on familiar, well stamped-in associations of red–stop and green–go to elicit habitual control, and establish dual feedback (i.e., monetary reward paired with cumulative performance tracking) as an intervention to break these well-learned habits to restore goal-directed control. The familiar stimuli in our task evoke a color-response habit that is evident in our participants’ difficulty overriding the well-established red-stop and green-go associations. We found that the familiar stimuli yield persistent instrumental responses even when these contingencies are manipulated to render green-go and red-stop color-responses disadvantageous for task performance. We also report enhanced goal-directed control (i.e., a disruption of the color-response habits) due to dual feedback, lending support to the effectiveness and scope of our performance enhancing feedback manipulation.

Accordingly, an important goal of our study was to establish our paradigm as a tool that captures real-world habits. In Experiment 1, we demonstrated the rigidity of the familiar green– go and red–stop contingencies compared to the newly-learned, flexible associations. The outcome-insensitive responses elicited by the familiar stimuli were reflected by a significant mapping-related impairment not observed when participants managed novel stimuli. Specifically, participants had more difficulty with the green-NoGo association in relation to red-NoGo, whereas variations in color-response mappings did not produce significant differences when managing novel associations (e.g., blue-NoGo or purple-NoGo). It is worth mentioning that the habits demonstrated here are not effector specific, in that we do not assert whether red and green light stimuli trigger actions that are alike those that may be triggered in a driving context (e.g., a foot-press response at red, or foot-release at green). Rather, the familiar stimuli used in our task may be evoking a general approach and avoid response, which, in the context of the task, is mapped onto Go and NoGo responses.

If these familiar red and green stimuli elicit outcome-insensitive habits, it may be argued the color-response mapping that is incongruent with daily experiences should display the lowest accuracy rates. However, in Experiment 1, green-NoGo accuracy was comparable to those of blue or purple-NoGo mappings. This pattern may be due to between-subject designs requiring more power than within-subject designs [36], thus making it more difficult to detect a potential decrement in green-NoGo accuracy. To test this hypothesis, we combined the data from the No-Feedback groups in Experiments 2 and 3, where participants underwent identical procedures. We found that the green-NoGo mapping produced significantly lower accuracy rates than either novel color-response mapping despite the between-subjects design. Furthermore, in a version of this task that employs a within-subject design in which all participants manage familiar and novel Go/NoGo contingencies, we indeed report significantly lower accuracy rates to green as a NoGo stimulus compared to all other colors (Ceceli et al., in press).

We then tested the strength of the habits evoked in our paradigm by introducing a motivation-based intervention: cumulative performance feedback. This type of feedback was not successful in preventing habitual control, supporting the notion that these existing habits are rigid enough to prevail even in the face of a motivational intervention. Nonetheless, performance feedback was able to produce promising results via secondary assays of behavioral flexibility. Namely, the prevention of habitual “Go” actions motivated the augmentation of our feedback manipulation to amplify its effect on motivational control. In Experiment 3, our combined delivery of performance and monetary feedback prevented the mapping-related impairment that is the result of a habit-dominated action control system, possibly improving goal-directed control by enhancing the salience of the outcome. In sum, we demonstrated well-existing habits, tested the limits of their associative strength, and provided the foundation for better understanding the restoration of goal-directed control.

Many habit paradigms that emulate the outcome-insensitive nature of habits have in common a shortcoming that limits generalizability to the typical habit experience: difficulty capturing well-learned habits in the lab that can provide a platform for studying habit disruption. Habit strength is limited by the participants’ brief exposure to experimental paradigms, and targeting these behaviors that are rendered inflexible in the lab may not be representative of habits encountered in the real world [18]. Perhaps due to these difficulties, well-learned habits and habit disruption research have been relatively better-represented in field experiments compared to the laboratory setting. For example, several field studies have examined the efficacy of interventions to change various presentations of daily habits, such as recycling and snacking habits [37–39]. However, recent efforts to bridge lab and field experiments have shown promising results. Although not an experiment of habit disruption, in a recent report, the slips-of-action task in the lab was examined alongside a more ecologically-relevant representation of habits—namely the habit of using one’s house keys. In this study, participants demonstrated an outcome-insensitive habit by making key choice errors, such that they persisted in choosing the incorrect key following a change in key covers. The attentional underpinnings of this behavior significantly correlated with slips of action performance, underlining the importance of focusing on well-established behaviors for an improved empirical approach to habit research [40].

One strategy that has proven beneficial in tackling habit change is implementation intentions, which provides individuals with an if-then plan (i.e., “if X happens, I will do Y”; or in a lab task, “if I see stimulus X, I will press Y”)—an aid to override unwanted or inflexible behaviors [41]. In the lab, implementation intentions have produced promising results, albeit with limited efficacy in disrupting strong habits. For instance, Webb and colleagues trained participants for five days on a target detection task, and successfully disrupted this lab-automated association using implementation intentions. However, this planning strategy did not break unwanted smoking habits, lending credence to the idea that the experimental resources at our disposal may not be sufficient in effectively stopping well-established habits [42]. Although this study approached habitual control from an attentional rather than a value-driven perspective, paralleling evidence from the motivational control literature has recently been reported. In another lab study, Verhoeven et al. employed planning strategies within a single experimental session to reduce action slips in an outcome-devaluation task [43]. Implementation intentions were more effective than goal-intentions (an outcome-based planning strategy, such as “I will not press for outcome X”) in reducing action slips when managing abstract images as outcomes, suggesting that implementation intentions may serve as a promising strategy in studying habit disruption—however, effective paradigms to demonstrate well-learned, outcome-insensitive habits, and an intervention to disrupt them are needed. In our study, we developed a task that allowed us to directly capture ecologically significant, well-established habits via the familiar green–go and red–stop associations. We present our Go/NoGo task with familiar and novel stimuli as a strong candidate for demonstrating habitual behaviors—bridging the success of field studies with the rigor and controllability of lab experimentation. We also illustrate that a salient feedback-based intervention may be utilized to shift cue-driven performance to become value-driven, laying the foundation to translational applications.

Our work also asserts that the use of familiar stimuli may circumvent the obstacles of training length and stimulus–response strength in habit research—an important step in improving paradigms to foster effective habit disruption strategies. A few prior studies have considered a similar approach. In a study investigating habits in substance use disorder, McKim and colleagues induced stimulus familiarity by pre-training a set of stimuli, and tested the strength of the familiar versus novel stimulus sets on a subsequent day via the reversal of a sub-set of these contingencies [17]. They found that compared to healthy controls, individuals with substance use disorder performed better in well-learned stimulus–response execution, yet exhibited impairments in managing contingency reversal. In accord with these findings, our study reveals that when managing contingencies that have been well-established throughout development— beyond an experimental pre-training stage—the recruitment of the habit system may also be evident in healthy individuals. Similarly, developmental and clinical researchers have used familiar green and red stimuli in Go/NoGo tasks with children suffering from attention deficit/hyperactivity disorder, as well as healthy adults to reduce task demands, and justified their decision by identifying these colors as having developmental relevance [19,20]. These prior reports highlight the utility of capitalizing on existing associations when examining habits, especially for clinical examinations of behavioral rigidity. Thus, we further contribute to the literature by introducing a task that requires minimal familiarity training, and by the inclusion of a motivational strategy to disrupt the familiarity-driven outcome-insensitivity. These contributions may be especially useful for optimizing costly fMRI designs, and benefit future translational neuroscience work that aims to reveal the neural bases of habit disruption.

The science of habits is a domain with direct clinical applications. The treatment of habit-based pathologies (e.g., obsessive-compulsive disorder) are within the scope of the habit literature, yet our field’s disproportionate focus on the formation of rigid behaviors, rather than overcoming well-formed habits, limits the translational impact of our research [44]. Indeed, several studies have highlighted the habitual aspects of various clinical disorders, as well as their underlying neural mechanisms [e.g., 7,8,17,45–50]. Researchers have further employed neurotransmitter depletion to emulate the biochemical profiles of psychopathologies to detect action control deficits [9,51,52]. Sub-clinical symptom presentation has also been investigated from the perspective of action control [53–56]. Furthermore, the multi-faceted role of stress in dictating motivated behaviors has been extensively demonstrated under acute, chronic, interaction of acute and chronic, and pharmacologically induced stress hormone reactivity [57–64]. Therefore, although researchers have characterized numerous contexts in which habits are prevalent, interventions that restore goal-directed motivational control have not been examined with similar vigor. As we demonstrate the habit-breaking effects of pairing monetary reward with cumulative performance feedback to amplify the salience of goals, we highlight the need for research avenues that not only identify goal-directed control deficits in clinical disorders, but work toward restoring these deficits to improve treatment strategies and quality of life.

### Conclusions

The disproportionate focus on habit formation and expression in the literature motivated us to direct our efforts to an area of habit research that has been less-explored: habit disruption. Although much research now confirms the habitual aspects of various pathologies, studies examining the restoration of these behavioral rigidities are relatively scarce. Here, we introduce a task that allows us to examine a more complete signature of motivational control by capturing well-learned habits and newly-learned goal-directed behaviors, as well as the possibility to test manipulations that may restore deliberate control. This method may be especially beneficial for understanding the neural markers of motivational control in healthy and compromised populations, as it capitalizes on existing associations that do not require extended lab-training. We also underline the efficacy of feedback in disrupting well-learned habits and promoting outcome-driven, goal-directed behaviors. This motivation-based manipulation may further inform the mechanisms underlying the habit disruption process—a translationally valuable research domain with direct clinical relevance.

## Supporting information

Datasets and plotting scripts

## Acknowledgements

This work was supported by a grant from the National Science Foundation (BCS1150708) awarded to Elizabeth Tricomi. We thank Zana Hariri, Sarah Ramirez, Christine Oti, and Charlie Ndouli for their assistance in data collection. We appreciate the helpful feedback from John O’Doherty and Omar D. Perez on a previous draft of this manuscript.

## Supporting information

**S1 Table. Summary of the Hierarchical Multiple Regression Model for Outcome-Insensitivity as Assayed by ΔNoGo_Accuracy.** Top layer of table depicts all regressors included in the hierarchical model and their respective statistics. Bottom layer of table, Model Summary Statistics, depicts the predictive strength of each model. Delta R^2^ (ΔR^2^) and corresponding F_change_ values denote the specific improvement of Model 2 over Model 1 in predicting the dependent variable. Toler. = Tolerance; VIF = Variance Inflation Factor. Significant p-values (alpha = .05) depicted in bold typeface.

**S2 Table. Summary of the Hierarchical Multiple Regression Model for Outcome-Insensitivity as Assayed by ΔGo_Accuracy.** Top layer of table depicts all regressors included in the hierarchical model and their respective statistics. Bottom layer of table, Model Summary Statistics, depicts the predictive strength of each model. Delta R2 (ΔR2) and corresponding F_change_ values denote the specific improvement of Model 2 over Model 1 in predicting the dependent variable. Toler. = Tolerance; VIF = Variance Inflation Factor. Significant p-values (alpha = .05) depicted in bold typeface.

**S3 Table. Summary of the Hierarchical Multiple Regression Model for Outcome-Insensitivity as Assayed by ΔNoGo_Accuracy.** Top layer of table depicts all regressors included in the hierarchical model and their respective statistics. Bottom layer of table, Model Summary Statistics, depicts the predictive strength of each model. Delta R^2^ (ΔR^2^) and corresponding F_change_ values denote the specific improvement of Model 2 over Model 1 in predicting the dependent variable. Toler. = Tolerance; VIF = Variance Inflation Factor. Significant p-values (alpha = .05) depicted in bold typeface.

**S4 Table. Summary of the Hierarchical Multiple Regression Model for Outcome-Insensitivity as Assayed by ΔGo_Accuracy.** Top layer of table depicts all regressors included in the hierarchical model and their respective statistics. Bottom layer of table, Model Summary Statistics, depicts the predictive strength of each model. Delta R^2^ (ΔR^2^) and corresponding F_change_ values denote the specific improvement of Model 2 over Model 1 in predicting the dependent variable. Toler. = Tolerance; VIF = Variance Inflation Factor. Significant p-values (alpha = .05) depicted in bold typeface.

**S5 Table. Summary of the Hierarchical Multiple Regression Model for Outcome-Insensitivity as Assayed by ΔNoGo_Accuracy.** Top layer of table depicts all regressors included in the hierarchical model and their respective statistics. Bottom layer of table, Model Summary Statistics, depicts the predictive strength of each model. Delta R^2^ (ΔR^2^) and corresponding F_change_ values denote the specific improvement of Model 2 over Model 1 in predicting the dependent variable. Toler. = Tolerance; VIF = Variance Inflation Factor. Significant p-values (alpha = .05) depicted in bold typeface.

**S6 Table. Summary of the Hierarchical Multiple Regression Model for Outcome-Insensitivity as Assayed by ΔGo_Accuracy.** Top layer of table depicts all regressors included in the hierarchical model and their respective statistics. Bottom layer of table, Model Summary Statistics, depicts the predictive strength of each model. Delta (Δ) values denote the specific improvement of Model 2 over Model 1 in predicting the dependent variable. Toler. = Tolerance; VIF = Variance Inflation Factor. Significant p-values (alpha = .05) depicted in bold typeface.

