## Supplementary material for "Demonstrating and disrupting well-learned habits": Datasets and plotting scripts: Experiment1_Python_html_Output.html

Experiment1\_Final\_Script


Experiment 1 Data. Stack data to facilitate plotting. Report NoGo and Go graphs. Exp 2 and 3 data found in their respective folders.

In [1]:

```
# import packages
import pandas as pd
import matplotlib as mpl
import matplotlib.pyplot as plt
import seaborn as sns
import numpy as np
from scipy import stats
from matplotlib.patches import Patch
%matplotlib inline
```

In [2]:

```
xls=pd.ExcelFile("Experiment1_Data_Final.xlsx")
df=pd.read_excel(xls, "Data_Clean")
df
```

Out[2]:

|  | Subj\_ID | Stim\_Cond | Order | Go\_RT | Go\_Rev\_RT | Go\_ACC | Go\_Rev\_ACC | NoGo\_ACC | NoGo\_Rev\_ACC | NoGo\_Diff | Go\_Diff | BIS |
| --- | --- | --- | --- | --- | --- | --- | --- | --- | --- | --- | --- | --- |
| 0 | 1 | Familiar | RegFirst | 335.397727 | 315.988889 | 88 | 90 | 95.0 | 75.0 | -20.0 | 2 | 63 |
| 1 | 5 | Familiar | RegFirst | 322.927083 | 318.043956 | 96 | 91 | 90.0 | 90.0 | 0.0 | -5 | 65 |
| 2 | 9 | Familiar | RegFirst | 319.389474 | 312.139785 | 95 | 93 | 90.0 | 95.0 | 5.0 | -2 | 68 |
| 3 | 13 | Familiar | RegFirst | 325.272727 | 310.247191 | 88 | 89 | 60.0 | 55.0 | -5.0 | 1 | 73 |
| 4 | 17 | Familiar | RegFirst | 304.448718 | 308.848837 | 78 | 86 | 55.0 | 35.0 | -20.0 | 8 | 77 |
| 5 | 21 | Familiar | RegFirst | 284.697917 | 280.489796 | 96 | 98 | 75.0 | 75.0 | 0.0 | 2 | 67 |
| 6 | 25 | Familiar | RegFirst | 319.821429 | 319.074074 | 84 | 81 | 60.0 | 70.0 | 10.0 | -3 | 49 |
| 7 | 29 | Familiar | RegFirst | 320.340909 | 312.430380 | 88 | 79 | 70.0 | 35.0 | -35.0 | -9 | 65 |
| 8 | 31 | Familiar | RegFirst | 302.266667 | 291.416667 | 90 | 84 | 45.0 | 35.0 | -10.0 | -6 | 78 |
| 9 | 37 | Familiar | RegFirst | 315.151515 | 314.479167 | 99 | 96 | 95.0 | 90.0 | -5.0 | -3 | 72 |
| 10 | 41 | Familiar | RegFirst | 330.670213 | 318.045977 | 94 | 87 | 80.0 | 60.0 | -20.0 | -7 | 65 |
| 11 | 45 | Familiar | RegFirst | 323.515789 | 318.260870 | 95 | 92 | 85.0 | 70.0 | -15.0 | -3 | 75 |
| 12 | 47 | Familiar | RegFirst | 315.720930 | 311.197531 | 86 | 81 | 60.0 | 75.0 | 15.0 | -5 | 66 |
| 13 | 2 | Familiar | RevFirst | 324.261364 | 318.534091 | 88 | 88 | 70.0 | 75.0 | 5.0 | 0 | 66 |
| 14 | 6 | Familiar | RevFirst | 320.415730 | 313.954545 | 89 | 88 | 95.0 | 60.0 | -35.0 | -1 | 62 |
| 15 | 10 | Familiar | RevFirst | 312.532609 | 322.576471 | 92 | 85 | 95.0 | 75.0 | -20.0 | -7 | 83 |
| 16 | 14 | Familiar | RevFirst | 325.065789 | 327.197368 | 76 | 76 | 70.0 | 25.0 | -45.0 | 0 | 65 |
| 17 | 18 | Familiar | RevFirst | 301.536082 | 315.111111 | 97 | 90 | 70.0 | 55.0 | -15.0 | -7 | 69 |
| 18 | 22 | Familiar | RevFirst | 311.752577 | 311.462366 | 97 | 93 | 95.0 | 90.0 | -5.0 | -4 | 65 |
| 19 | 26 | Familiar | RevFirst | 307.989583 | 337.404762 | 96 | 84 | 90.0 | 85.0 | -5.0 | -12 | 82 |
| 20 | 28 | Familiar | RevFirst | 324.937500 | 322.682353 | 80 | 85 | 80.0 | 65.0 | -15.0 | 5 | 62 |
| 21 | 34 | Familiar | RevFirst | 288.178947 | 302.098901 | 95 | 91 | 85.0 | 70.0 | -15.0 | -4 | 81 |
| 22 | 38 | Familiar | RevFirst | 317.450549 | 311.735632 | 91 | 87 | 60.0 | 70.0 | 10.0 | -4 | 74 |
| 23 | 42 | Familiar | RevFirst | 304.563830 | 324.241758 | 94 | 91 | 75.0 | 60.0 | -15.0 | -3 | 82 |
| 24 | 46 | Familiar | RevFirst | 297.709677 | 301.695652 | 93 | 92 | 75.0 | 70.0 | -5.0 | -1 | 54 |
| 25 | 3 | Novel | RegFirst | 307.747368 | 314.226804 | 95 | 97 | 100.0 | 95.0 | -5.0 | 2 | 62 |
| 26 | 7 | Novel | RegFirst | 313.717391 | 312.092784 | 92 | 97 | 85.0 | 85.0 | 0.0 | 5 | 70 |
| 27 | 11 | Novel | RegFirst | 313.516484 | 302.934783 | 91 | 92 | 85.0 | 75.0 | -10.0 | 1 | 66 |
| 28 | 15 | Novel | RegFirst | 305.589474 | 294.894737 | 95 | 95 | 70.0 | 70.0 | 0.0 | 0 | 67 |
| 29 | 19 | Novel | RegFirst | 339.372093 | 339.723404 | 86 | 94 | 75.0 | 85.0 | 10.0 | 8 | 77 |
| 30 | 23 | Novel | RegFirst | 307.958763 | 284.927835 | 97 | 97 | 85.0 | 85.0 | 0.0 | 0 | 72 |
| 31 | 27 | Novel | RegFirst | 297.061224 | 306.042553 | 98 | 94 | 45.0 | 50.0 | 5.0 | -4 | 66 |
| 32 | 33 | Novel | RegFirst | 309.034483 | 290.370787 | 87 | 89 | 40.0 | 35.0 | -5.0 | 2 | 66 |
| 33 | 36 | Novel | RegFirst | 322.098901 | 314.074468 | 91 | 94 | 65.0 | 85.0 | 20.0 | 3 | 64 |
| 34 | 39 | Novel | RegFirst | 309.382022 | 308.000000 | 89 | 87 | 70.0 | 50.0 | -20.0 | -2 | 75 |
| 35 | 43 | Novel | RegFirst | 326.743590 | 311.309524 | 78 | 84 | 50.0 | 35.0 | -15.0 | 6 | 71 |
| 36 | 49 | Novel | RegFirst | 329.752809 | 314.168539 | 89 | 89 | 85.0 | 75.0 | -10.0 | 0 | 70 |
| 37 | 4 | Novel | RevFirst | 302.597826 | 310.720430 | 92 | 93 | 60.0 | 75.0 | 15.0 | 1 | 73 |
| 38 | 8 | Novel | RevFirst | 307.348837 | 324.373626 | 86 | 91 | 65.0 | 70.0 | 5.0 | 5 | 63 |
| 39 | 12 | Novel | RevFirst | 327.000000 | 326.029851 | 69 | 67 | 50.0 | 70.0 | 20.0 | -2 | 78 |
| 40 | 16 | Novel | RevFirst | 312.894737 | 312.195652 | 95 | 92 | 40.0 | 60.0 | 20.0 | -3 | 70 |
| 41 | 20 | Novel | RevFirst | 337.395349 | 335.913580 | 86 | 81 | 60.0 | 85.0 | 25.0 | -5 | 68 |
| 42 | 24 | Novel | RevFirst | 308.150000 | 312.487500 | 80 | 80 | 45.0 | 70.0 | 25.0 | 0 | 73 |
| 43 | 30 | Novel | RevFirst | 299.382979 | 321.681818 | 94 | 88 | 70.0 | 65.0 | -5.0 | -6 | 71 |
| 44 | 32 | Novel | RevFirst | 346.345238 | 343.510870 | 84 | 92 | 80.0 | 75.0 | -5.0 | 8 | 70 |
| 45 | 35 | Novel | RevFirst | 225.488889 | 305.528736 | 90 | 87 | 10.0 | 45.0 | 35.0 | -3 | 68 |
| 46 | 40 | Novel | RevFirst | 302.000000 | 295.826531 | 93 | 98 | 95.0 | 90.0 | -5.0 | 5 | 70 |
| 47 | 44 | Novel | RevFirst | 294.182927 | 254.381579 | 82 | 76 | 55.0 | 35.0 | -20.0 | -6 | 70 |
| 48 | 48 | Novel | RevFirst | 322.569767 | 303.126437 | 86 | 87 | 70.0 | 45.0 | -25.0 | 1 | 86 |
| 49 | 50 | Novel | RevFirst | 315.971831 | 336.304348 | 71 | 46 | 65.0 | 80.0 | 15.0 | -25 | 63 |

Congruency refers to the within-subject Mapping factor. In the Familiar condition, Congruent means Red:NoGo and Green:Go--congruent with daily experiences, whereas in the Novel condition, these congruency mappings are arbitrary: Purple:Go, Blue:NoGo.

In [3]:

```
stacked_data=pd.melt(df, id_vars=["Subj_ID", "Stim_Cond"], value_vars=["NoGo_ACC", "NoGo_Rev_ACC"], 
        var_name="Signal", value_name="Accuracy")
def conditions_phase(x):
    if x == "NoGo_ACC":
        return "Congruent"
    elif x == "NoGo_Rev_ACC":
        return "Incongruent"
func = np.vectorize(conditions_phase)
stacked_data["Congruency"] = func(stacked_data["Signal"])

stacked_data
```

Out[3]:

|  | Subj\_ID | Stim\_Cond | Signal | Accuracy | Congruency |
| --- | --- | --- | --- | --- | --- |
| 0 | 1 | Familiar | NoGo\_ACC | 95.0 | Congruent |
| 1 | 5 | Familiar | NoGo\_ACC | 90.0 | Congruent |
| 2 | 9 | Familiar | NoGo\_ACC | 90.0 | Congruent |
| 3 | 13 | Familiar | NoGo\_ACC | 60.0 | Congruent |
| 4 | 17 | Familiar | NoGo\_ACC | 55.0 | Congruent |
| 5 | 21 | Familiar | NoGo\_ACC | 75.0 | Congruent |
| 6 | 25 | Familiar | NoGo\_ACC | 60.0 | Congruent |
| 7 | 29 | Familiar | NoGo\_ACC | 70.0 | Congruent |
| 8 | 31 | Familiar | NoGo\_ACC | 45.0 | Congruent |
| 9 | 37 | Familiar | NoGo\_ACC | 95.0 | Congruent |
| 10 | 41 | Familiar | NoGo\_ACC | 80.0 | Congruent |
| 11 | 45 | Familiar | NoGo\_ACC | 85.0 | Congruent |
| 12 | 47 | Familiar | NoGo\_ACC | 60.0 | Congruent |
| 13 | 2 | Familiar | NoGo\_ACC | 70.0 | Congruent |
| 14 | 6 | Familiar | NoGo\_ACC | 95.0 | Congruent |
| 15 | 10 | Familiar | NoGo\_ACC | 95.0 | Congruent |
| 16 | 14 | Familiar | NoGo\_ACC | 70.0 | Congruent |
| 17 | 18 | Familiar | NoGo\_ACC | 70.0 | Congruent |
| 18 | 22 | Familiar | NoGo\_ACC | 95.0 | Congruent |
| 19 | 26 | Familiar | NoGo\_ACC | 90.0 | Congruent |
| 20 | 28 | Familiar | NoGo\_ACC | 80.0 | Congruent |
| 21 | 34 | Familiar | NoGo\_ACC | 85.0 | Congruent |
| 22 | 38 | Familiar | NoGo\_ACC | 60.0 | Congruent |
| 23 | 42 | Familiar | NoGo\_ACC | 75.0 | Congruent |
| 24 | 46 | Familiar | NoGo\_ACC | 75.0 | Congruent |
| 25 | 3 | Novel | NoGo\_ACC | 100.0 | Congruent |
| 26 | 7 | Novel | NoGo\_ACC | 85.0 | Congruent |
| 27 | 11 | Novel | NoGo\_ACC | 85.0 | Congruent |
| 28 | 15 | Novel | NoGo\_ACC | 70.0 | Congruent |
| 29 | 19 | Novel | NoGo\_ACC | 75.0 | Congruent |
| ... | ... | ... | ... | ... | ... |
| 70 | 28 | Familiar | NoGo\_Rev\_ACC | 65.0 | Incongruent |
| 71 | 34 | Familiar | NoGo\_Rev\_ACC | 70.0 | Incongruent |
| 72 | 38 | Familiar | NoGo\_Rev\_ACC | 70.0 | Incongruent |
| 73 | 42 | Familiar | NoGo\_Rev\_ACC | 60.0 | Incongruent |
| 74 | 46 | Familiar | NoGo\_Rev\_ACC | 70.0 | Incongruent |
| 75 | 3 | Novel | NoGo\_Rev\_ACC | 95.0 | Incongruent |
| 76 | 7 | Novel | NoGo\_Rev\_ACC | 85.0 | Incongruent |
| 77 | 11 | Novel | NoGo\_Rev\_ACC | 75.0 | Incongruent |
| 78 | 15 | Novel | NoGo\_Rev\_ACC | 70.0 | Incongruent |
| 79 | 19 | Novel | NoGo\_Rev\_ACC | 85.0 | Incongruent |
| 80 | 23 | Novel | NoGo\_Rev\_ACC | 85.0 | Incongruent |
| 81 | 27 | Novel | NoGo\_Rev\_ACC | 50.0 | Incongruent |
| 82 | 33 | Novel | NoGo\_Rev\_ACC | 35.0 | Incongruent |
| 83 | 36 | Novel | NoGo\_Rev\_ACC | 85.0 | Incongruent |
| 84 | 39 | Novel | NoGo\_Rev\_ACC | 50.0 | Incongruent |
| 85 | 43 | Novel | NoGo\_Rev\_ACC | 35.0 | Incongruent |
| 86 | 49 | Novel | NoGo\_Rev\_ACC | 75.0 | Incongruent |
| 87 | 4 | Novel | NoGo\_Rev\_ACC | 75.0 | Incongruent |
| 88 | 8 | Novel | NoGo\_Rev\_ACC | 70.0 | Incongruent |
| 89 | 12 | Novel | NoGo\_Rev\_ACC | 70.0 | Incongruent |
| 90 | 16 | Novel | NoGo\_Rev\_ACC | 60.0 | Incongruent |
| 91 | 20 | Novel | NoGo\_Rev\_ACC | 85.0 | Incongruent |
| 92 | 24 | Novel | NoGo\_Rev\_ACC | 70.0 | Incongruent |
| 93 | 30 | Novel | NoGo\_Rev\_ACC | 65.0 | Incongruent |
| 94 | 32 | Novel | NoGo\_Rev\_ACC | 75.0 | Incongruent |
| 95 | 35 | Novel | NoGo\_Rev\_ACC | 45.0 | Incongruent |
| 96 | 40 | Novel | NoGo\_Rev\_ACC | 90.0 | Incongruent |
| 97 | 44 | Novel | NoGo\_Rev\_ACC | 35.0 | Incongruent |
| 98 | 48 | Novel | NoGo\_Rev\_ACC | 45.0 | Incongruent |
| 99 | 50 | Novel | NoGo\_Rev\_ACC | 80.0 | Incongruent |

100 rows × 5 columns

In [4]:

```
sns.set(style="white", font="Times New Roman", context="notebook", font_scale=1.3)
ax = sns.barplot(x="Stim_Cond", y="Accuracy", hue="Congruency", palette=["#ff0000", "#03d547"], ci=68, capsize=0.02, data=stacked_data)
ax.patches[1].set_facecolor("#d12fdf")
ax.patches[3].set_facecolor("#1d47f5")
plt.title("Familiar stimuli elicit mapping-related \nimpairments in NoGo accuracy", weight="bold", y=1.08, fontsize=20)
plt.ylabel("NoGo Accuracy (%)", weight="bold", labelpad=5, fontsize=20)
plt.xticks(weight="bold", fontsize=20)
sns.despine(bottom=False)
ax.set_xlabel("")
ax.set_ylim(50,85)
ax.legend_.remove()
#add significance asterisk and line
x1, x2 = -0.20, 0.18   # only two columns, so they would be 0, 1
y, h, col = stacked_data['Accuracy'].mean()+15, 0.5, "k" #y will be the height of the line and star(2 points above the mean), h will be the height of the two lines
                                          #pointing down--0.2, and col is the color--black coded as k 
plt.plot([x1, x1, x2, x2], [y, y+h, y+h, y], lw=1, c=col) #here we plot this line and star on top of our barplot
plt.text((x1+x2)*0.5, y+h-0.1, "*", ha='center', va='bottom', color=col)
#plt.savefig("Exp1_NoGo_graph.tiff", bbox_inches="tight", dpi=1000)
plt.show()
```

In [5]:

```
stacked_data_go=pd.melt(df, id_vars=["Subj_ID", "Stim_Cond"], value_vars=["Go_ACC", "Go_Rev_ACC"], 
        var_name="Signal", value_name="Accuracy")
def conditions_phase(x):
    if x == "Go_ACC":
        return "Congruent"
    elif x == "Go_Rev_ACC":
        return "Incongruent"
func = np.vectorize(conditions_phase)
stacked_data_go["Congruency"] = func(stacked_data_go["Signal"])

stacked_data_go
```

Out[5]:

|  | Subj\_ID | Stim\_Cond | Signal | Accuracy | Congruency |
| --- | --- | --- | --- | --- | --- |
| 0 | 1 | Familiar | Go\_ACC | 88 | Congruent |
| 1 | 5 | Familiar | Go\_ACC | 96 | Congruent |
| 2 | 9 | Familiar | Go\_ACC | 95 | Congruent |
| 3 | 13 | Familiar | Go\_ACC | 88 | Congruent |
| 4 | 17 | Familiar | Go\_ACC | 78 | Congruent |
| 5 | 21 | Familiar | Go\_ACC | 96 | Congruent |
| 6 | 25 | Familiar | Go\_ACC | 84 | Congruent |
| 7 | 29 | Familiar | Go\_ACC | 88 | Congruent |
| 8 | 31 | Familiar | Go\_ACC | 90 | Congruent |
| 9 | 37 | Familiar | Go\_ACC | 99 | Congruent |
| 10 | 41 | Familiar | Go\_ACC | 94 | Congruent |
| 11 | 45 | Familiar | Go\_ACC | 95 | Congruent |
| 12 | 47 | Familiar | Go\_ACC | 86 | Congruent |
| 13 | 2 | Familiar | Go\_ACC | 88 | Congruent |
| 14 | 6 | Familiar | Go\_ACC | 89 | Congruent |
| 15 | 10 | Familiar | Go\_ACC | 92 | Congruent |
| 16 | 14 | Familiar | Go\_ACC | 76 | Congruent |
| 17 | 18 | Familiar | Go\_ACC | 97 | Congruent |
| 18 | 22 | Familiar | Go\_ACC | 97 | Congruent |
| 19 | 26 | Familiar | Go\_ACC | 96 | Congruent |
| 20 | 28 | Familiar | Go\_ACC | 80 | Congruent |
| 21 | 34 | Familiar | Go\_ACC | 95 | Congruent |
| 22 | 38 | Familiar | Go\_ACC | 91 | Congruent |
| 23 | 42 | Familiar | Go\_ACC | 94 | Congruent |
| 24 | 46 | Familiar | Go\_ACC | 93 | Congruent |
| 25 | 3 | Novel | Go\_ACC | 95 | Congruent |
| 26 | 7 | Novel | Go\_ACC | 92 | Congruent |
| 27 | 11 | Novel | Go\_ACC | 91 | Congruent |
| 28 | 15 | Novel | Go\_ACC | 95 | Congruent |
| 29 | 19 | Novel | Go\_ACC | 86 | Congruent |
| ... | ... | ... | ... | ... | ... |
| 70 | 28 | Familiar | Go\_Rev\_ACC | 85 | Incongruent |
| 71 | 34 | Familiar | Go\_Rev\_ACC | 91 | Incongruent |
| 72 | 38 | Familiar | Go\_Rev\_ACC | 87 | Incongruent |
| 73 | 42 | Familiar | Go\_Rev\_ACC | 91 | Incongruent |
| 74 | 46 | Familiar | Go\_Rev\_ACC | 92 | Incongruent |
| 75 | 3 | Novel | Go\_Rev\_ACC | 97 | Incongruent |
| 76 | 7 | Novel | Go\_Rev\_ACC | 97 | Incongruent |
| 77 | 11 | Novel | Go\_Rev\_ACC | 92 | Incongruent |
| 78 | 15 | Novel | Go\_Rev\_ACC | 95 | Incongruent |
| 79 | 19 | Novel | Go\_Rev\_ACC | 94 | Incongruent |
| 80 | 23 | Novel | Go\_Rev\_ACC | 97 | Incongruent |
| 81 | 27 | Novel | Go\_Rev\_ACC | 94 | Incongruent |
| 82 | 33 | Novel | Go\_Rev\_ACC | 89 | Incongruent |
| 83 | 36 | Novel | Go\_Rev\_ACC | 94 | Incongruent |
| 84 | 39 | Novel | Go\_Rev\_ACC | 87 | Incongruent |
| 85 | 43 | Novel | Go\_Rev\_ACC | 84 | Incongruent |
| 86 | 49 | Novel | Go\_Rev\_ACC | 89 | Incongruent |
| 87 | 4 | Novel | Go\_Rev\_ACC | 93 | Incongruent |
| 88 | 8 | Novel | Go\_Rev\_ACC | 91 | Incongruent |
| 89 | 12 | Novel | Go\_Rev\_ACC | 67 | Incongruent |
| 90 | 16 | Novel | Go\_Rev\_ACC | 92 | Incongruent |
| 91 | 20 | Novel | Go\_Rev\_ACC | 81 | Incongruent |
| 92 | 24 | Novel | Go\_Rev\_ACC | 80 | Incongruent |
| 93 | 30 | Novel | Go\_Rev\_ACC | 88 | Incongruent |
| 94 | 32 | Novel | Go\_Rev\_ACC | 92 | Incongruent |
| 95 | 35 | Novel | Go\_Rev\_ACC | 87 | Incongruent |
| 96 | 40 | Novel | Go\_Rev\_ACC | 98 | Incongruent |
| 97 | 44 | Novel | Go\_Rev\_ACC | 76 | Incongruent |
| 98 | 48 | Novel | Go\_Rev\_ACC | 87 | Incongruent |
| 99 | 50 | Novel | Go\_Rev\_ACC | 46 | Incongruent |

100 rows × 5 columns

In [6]:

```
sns.set(style="white", font="Times New Roman", context="notebook", font_scale=1.3)
ax = sns.barplot(x="Stim_Cond", y="Accuracy", hue="Congruency", palette=["#03d547", "#ff0000"], ci=68, capsize=0.02, data=stacked_data_go)
ax.patches[1].set_facecolor("#1d47f5")
ax.patches[3].set_facecolor("#d12fdf")
plt.title("Familiar stimuli elicit mapping-related \nimpairments in Go accuracy", weight="bold", y=1.08, fontsize=20)
plt.ylabel("Go Accuracy (%)", weight="bold", labelpad=5, fontsize=20)
plt.xticks(weight="bold", fontsize=20)
sns.despine(bottom=False)
ax.set_xlabel("")
ax.set_ylim(50,100)
ax.legend_.remove()
#add significance asterisk and line
x1, x2 = -0.20, 0.18   # only two columns, so they would be 0, 1
y, h, col = stacked_data['Accuracy'].mean()+30, 0.5, "k" #y will be the height of the line and star(2 points above the mean), h will be the height of the two lines
                                          #pointing down--0.2, and col is the color--black coded as k 
plt.plot([x1, x1, x2, x2], [y, y+h, y+h, y], lw=1, c=col) #here we plot this line and star on top of our barplot
plt.text((x1+x2)*0.5, y+h-0.1, "*", ha='center', va='bottom', color=col)
#plt.savefig("Exp1_Go_graph.tiff", bbox_inches="tight", dpi=1000)
plt.show()
```
