## Supplementary material for "Demonstrating and disrupting well-learned habits": Datasets and plotting scripts: Experiment2_Python_html_Output.html

Experiment2\_Final\_Script


Experiment 2 data script. Melt data for graph creation.

In [1]:

```
#import all packages
import pandas as pd
import matplotlib as mpl
import matplotlib.pyplot as plt
import seaborn as sns
import numpy as np
from scipy import stats
from matplotlib.patches import Patch
%matplotlib inline
```

In [2]:

```
xls=pd.ExcelFile("Experiment2_Data_Final.xlsx")
df=pd.read_excel(xls, "Data_Clean")
pd.set_option("display.max_rows", 500)
df
```

Out[2]:

|  | Subj\_ID | Stim\_Cond | FB\_Cond | Go\_RT | Go\_Rev\_RT | Go\_ACC | Go\_Rev\_ACC | NoGo\_ACC | NoGo\_Rev\_ACC | NoGo\_Diff | Go\_Diff | BIS |
| --- | --- | --- | --- | --- | --- | --- | --- | --- | --- | --- | --- | --- |
| 0 | 1 | Familiar | PerfFB | 290.241379 | 287.908046 | 87 | 87 | 55 | 35 | -20 | 0 | 64 |
| 1 | 2 | Familiar | PerfFB | 289.876289 | 310.360825 | 97 | 97 | 90 | 90 | 0 | 0 | 67 |
| 2 | 3 | Familiar | PerfFB | 299.677419 | 299.947368 | 93 | 95 | 60 | 70 | 10 | 2 | 87 |
| 3 | 4 | Familiar | PerfFB | 309.978261 | 296.200000 | 92 | 90 | 70 | 65 | -5 | -2 | 74 |
| 4 | 5 | Familiar | PerfFB | 320.234043 | 298.547368 | 94 | 95 | 80 | 70 | -10 | 1 | 74 |
| 5 | 6 | Familiar | PerfFB | 317.938144 | 304.147727 | 97 | 88 | 80 | 80 | 0 | -9 | 75 |
| 6 | 7 | Familiar | PerfFB | 310.106383 | 312.934783 | 94 | 92 | 90 | 70 | -20 | -2 | 69 |
| 7 | 8 | Familiar | PerfFB | 313.693333 | 328.303797 | 75 | 79 | 85 | 70 | -15 | 4 | 80 |
| 8 | 9 | Familiar | PerfFB | 328.238636 | 324.897436 | 88 | 78 | 65 | 75 | 10 | -10 | 63 |
| 9 | 10 | Familiar | PerfFB | 309.263158 | 294.917526 | 95 | 97 | 75 | 85 | 10 | 2 | 76 |
| 10 | 11 | Familiar | PerfFB | 282.278351 | 291.978723 | 97 | 93 | 55 | 50 | -5 | -4 | 72 |
| 11 | 12 | Familiar | PerfFB | 292.236559 | 267.179775 | 93 | 89 | 75 | 45 | -30 | -4 | 73 |
| 12 | 13 | Familiar | PerfFB | 302.432990 | 289.294737 | 97 | 95 | 90 | 75 | -15 | -2 | 62 |
| 13 | 15 | Familiar | PerfFB | 298.189474 | 293.041667 | 95 | 96 | 65 | 50 | -15 | 1 | 77 |
| 14 | 16 | Familiar | PerfFB | 330.032609 | 299.521277 | 92 | 94 | 90 | 100 | 10 | 2 | 71 |
| 15 | 18 | Familiar | PerfFB | 298.159574 | 277.422222 | 94 | 90 | 60 | 35 | -25 | -4 | 81 |
| 16 | 19 | Familiar | PerfFB | 316.433333 | 311.541667 | 90 | 96 | 95 | 90 | -5 | 6 | 66 |
| 17 | 20 | Familiar | PerfFB | 313.076923 | 302.655556 | 91 | 90 | 60 | 70 | 10 | -1 | 78 |
| 18 | 23 | Familiar | PerfFB | 313.850575 | 291.934783 | 87 | 92 | 80 | 70 | -10 | 5 | 60 |
| 19 | 25 | Familiar | PerfFB | 319.630952 | 324.898876 | 84 | 89 | 80 | 55 | -25 | 5 | 64 |
| 20 | 115 | Familiar | PerfFB | 307.054348 | 317.054348 | 92 | 92 | 100 | 85 | -15 | 0 | 71 |
| 21 | 116 | Familiar | PerfFB | 279.282353 | 283.937500 | 82 | 80 | 65 | 30 | -35 | -2 | 68 |
| 22 | 118 | Familiar | PerfFB | 296.648352 | 295.897727 | 91 | 88 | 80 | 95 | 15 | -3 | 75 |
| 23 | 119 | Familiar | PerfFB | 284.072917 | 278.117021 | 96 | 94 | 95 | 70 | -25 | -2 | 66 |
| 24 | 120 | Familiar | PerfFB | 274.744444 | 276.785714 | 90 | 84 | 10 | 20 | 10 | -6 | 71 |
| 25 | 65 | Familiar | noPerfFB | 315.402439 | 308.977528 | 82 | 89 | 55 | 80 | 25 | 7 | 77 |
| 26 | 66 | Familiar | noPerfFB | 316.882979 | 328.850575 | 94 | 87 | 85 | 70 | -15 | -7 | 68 |
| 27 | 68 | Familiar | noPerfFB | 307.163265 | 283.413793 | 98 | 87 | 100 | 70 | -30 | -11 | 68 |
| 28 | 69 | Familiar | noPerfFB | 318.822222 | 313.369231 | 89 | 65 | 50 | 60 | 10 | -24 | 74 |
| 29 | 71 | Familiar | noPerfFB | 312.957447 | 309.563830 | 94 | 94 | 75 | 85 | 10 | 0 | 57 |
| 30 | 72 | Familiar | noPerfFB | 282.302083 | 300.333333 | 96 | 93 | 75 | 50 | -25 | -3 | 71 |
| 31 | 73 | Familiar | noPerfFB | 316.494624 | 311.710843 | 93 | 83 | 80 | 45 | -35 | -10 | 67 |
| 32 | 74 | Familiar | noPerfFB | 322.136364 | 322.268817 | 87 | 93 | 75 | 50 | -25 | 6 | 66 |
| 33 | 75 | Familiar | noPerfFB | 317.666667 | 314.494505 | 96 | 91 | 95 | 95 | 0 | -5 | 73 |
| 34 | 76 | Familiar | noPerfFB | 295.783505 | 304.069767 | 97 | 86 | 80 | 50 | -30 | -11 | 58 |
| 35 | 77 | Familiar | noPerfFB | 329.651163 | 352.450980 | 86 | 51 | 90 | 90 | 0 | -35 | 65 |
| 36 | 78 | Familiar | noPerfFB | 282.457447 | 272.109890 | 94 | 91 | 45 | 25 | -20 | -3 | 73 |
| 37 | 79 | Familiar | noPerfFB | 307.391753 | 299.729167 | 97 | 96 | 95 | 90 | -5 | -1 | 62 |
| 38 | 80 | Familiar | noPerfFB | 291.739583 | 294.903226 | 96 | 93 | 80 | 70 | -10 | -3 | 69 |
| 39 | 82 | Familiar | noPerfFB | 309.879121 | 303.662921 | 91 | 89 | 65 | 35 | -30 | -2 | 75 |
| 40 | 83 | Familiar | noPerfFB | 312.095745 | 303.043011 | 94 | 93 | 90 | 60 | -30 | -1 | 67 |
| 41 | 84 | Familiar | noPerfFB | 308.936842 | 297.149425 | 95 | 87 | 85 | 45 | -40 | -8 | 61 |
| 42 | 85 | Familiar | noPerfFB | 275.969072 | 301.291667 | 97 | 96 | 70 | 90 | 20 | -1 | 68 |
| 43 | 86 | Familiar | noPerfFB | 308.235955 | 319.024096 | 89 | 83 | 85 | 70 | -15 | -6 | 79 |
| 44 | 87 | Familiar | noPerfFB | 329.833333 | 340.528090 | 90 | 89 | 70 | 65 | -5 | -1 | 68 |
| 45 | 88 | Familiar | noPerfFB | 331.510204 | 317.840426 | 98 | 94 | 100 | 100 | 0 | -4 | 65 |
| 46 | 109 | Familiar | noPerfFB | 299.427083 | 303.265957 | 96 | 94 | 75 | 65 | -10 | -2 | 67 |
| 47 | 110 | Familiar | noPerfFB | 288.989474 | 293.440860 | 95 | 93 | 100 | 85 | -15 | -2 | 74 |
| 48 | 111 | Familiar | noPerfFB | 314.010753 | 327.080460 | 93 | 87 | 70 | 75 | 5 | -6 | 70 |
| 49 | 112 | Familiar | noPerfFB | 312.937500 | 300.373626 | 96 | 91 | 85 | 80 | -5 | -5 | 58 |
| 50 | 21 | Novel | PerfFB | 328.148936 | 324.726316 | 94 | 95 | 100 | 100 | 0 | 1 | 66 |
| 51 | 22 | Novel | PerfFB | 314.144444 | 309.597938 | 90 | 97 | 45 | 95 | 50 | 7 | 64 |
| 52 | 24 | Novel | PerfFB | 326.084337 | 329.318182 | 83 | 88 | 40 | 75 | 35 | 5 | 82 |
| 53 | 26 | Novel | PerfFB | 308.291667 | 296.947917 | 96 | 96 | 90 | 85 | -5 | 0 | 64 |
| 54 | 27 | Novel | PerfFB | 313.113402 | 329.515464 | 97 | 97 | 90 | 95 | 5 | 0 | 71 |
| 55 | 28 | Novel | PerfFB | 307.284091 | 310.090909 | 88 | 88 | 75 | 80 | 5 | 0 | 70 |
| 56 | 29 | Novel | PerfFB | 287.629213 | 304.553191 | 89 | 94 | 45 | 70 | 25 | 5 | 68 |
| 57 | 30 | Novel | PerfFB | 312.694118 | 291.375000 | 85 | 88 | 70 | 75 | 5 | 3 | 57 |
| 58 | 31 | Novel | PerfFB | 305.623656 | 310.020833 | 93 | 96 | 85 | 85 | 0 | 3 | 60 |
| 59 | 32 | Novel | PerfFB | 316.636364 | 318.208791 | 88 | 91 | 85 | 75 | -10 | 3 | 65 |
| 60 | 33 | Novel | PerfFB | 309.011236 | 294.322581 | 89 | 93 | 55 | 65 | 10 | 4 | 68 |
| 61 | 34 | Novel | PerfFB | 324.185185 | 304.612903 | 81 | 93 | 40 | 60 | 20 | 12 | 66 |
| 62 | 35 | Novel | PerfFB | 303.000000 | 306.646341 | 77 | 82 | 50 | 75 | 25 | 5 | 62 |
| 63 | 36 | Novel | PerfFB | 300.229885 | 300.397727 | 87 | 88 | 85 | 85 | 0 | 1 | 65 |
| 64 | 37 | Novel | PerfFB | 331.433333 | 301.366667 | 90 | 90 | 70 | 60 | -10 | 0 | 62 |
| 65 | 38 | Novel | PerfFB | 322.241379 | 296.952381 | 87 | 84 | 95 | 55 | -40 | -3 | 70 |
| 66 | 39 | Novel | PerfFB | 314.035714 | 306.869048 | 84 | 84 | 60 | 50 | -10 | 0 | 85 |
| 67 | 40 | Novel | PerfFB | 288.042105 | 286.864583 | 95 | 96 | 90 | 95 | 5 | 1 | 68 |
| 68 | 41 | Novel | PerfFB | 293.364706 | 293.325843 | 85 | 89 | 60 | 85 | 25 | 4 | 72 |
| 69 | 104 | Novel | PerfFB | 327.300000 | 331.868132 | 90 | 91 | 80 | 95 | 15 | 1 | 73 |
| 70 | 117 | Novel | PerfFB | 320.323944 | 308.352113 | 71 | 71 | 55 | 45 | -10 | 0 | 64 |
| 71 | 121 | Novel | PerfFB | 291.329670 | 317.821429 | 91 | 84 | 70 | 75 | 5 | -7 | 65 |
| 72 | 124 | Novel | PerfFB | 313.386364 | 299.937500 | 88 | 96 | 40 | 60 | 20 | 8 | 66 |
| 73 | 125 | Novel | PerfFB | 301.957447 | 304.138298 | 94 | 94 | 90 | 90 | 0 | 0 | 72 |
| 74 | 127 | Novel | PerfFB | 306.808989 | 314.321429 | 89 | 84 | 85 | 70 | -15 | -5 | 66 |
| 75 | 67 | Novel | noPerfFB | 319.607595 | 338.317460 | 79 | 63 | 70 | 80 | 10 | -16 | 85 |
| 76 | 70 | Novel | noPerfFB | 284.885417 | 301.979167 | 96 | 96 | 50 | 65 | 15 | 0 | 60 |
| 77 | 81 | Novel | noPerfFB | 302.628866 | 291.115789 | 97 | 95 | 80 | 75 | -5 | -2 | 67 |
| 78 | 89 | Novel | noPerfFB | 300.416667 | 294.010204 | 96 | 98 | 90 | 90 | 0 | 2 | 69 |
| 79 | 90 | Novel | noPerfFB | 304.056180 | 285.787234 | 89 | 94 | 40 | 35 | -5 | 5 | 71 |
| 80 | 91 | Novel | noPerfFB | 300.184783 | 289.845361 | 92 | 97 | 80 | 75 | -5 | 5 | 81 |
| 81 | 92 | Novel | noPerfFB | 303.860215 | 305.101010 | 93 | 99 | 100 | 95 | -5 | 6 | 69 |
| 82 | 94 | Novel | noPerfFB | 308.597826 | 324.211111 | 92 | 90 | 80 | 90 | 10 | -2 | 66 |
| 83 | 95 | Novel | noPerfFB | 325.388889 | 312.977011 | 90 | 87 | 75 | 55 | -20 | -3 | 65 |
| 84 | 96 | Novel | noPerfFB | 299.537634 | 286.694737 | 93 | 95 | 70 | 50 | -20 | 2 | 83 |
| 85 | 97 | Novel | noPerfFB | 317.989130 | 312.956989 | 92 | 93 | 70 | 65 | -5 | 1 | 77 |
| 86 | 98 | Novel | noPerfFB | 324.726190 | 307.376344 | 84 | 93 | 75 | 85 | 10 | 9 | 66 |
| 87 | 99 | Novel | noPerfFB | 324.634409 | 320.645833 | 93 | 96 | 85 | 95 | 10 | 3 | 80 |
| 88 | 100 | Novel | noPerfFB | 311.222222 | 306.276596 | 90 | 94 | 65 | 60 | -5 | 4 | 59 |
| 89 | 101 | Novel | noPerfFB | 286.670103 | 290.715789 | 97 | 95 | 75 | 50 | -25 | -2 | 67 |
| 90 | 102 | Novel | noPerfFB | 323.693182 | 317.602151 | 88 | 93 | 80 | 65 | -15 | 5 | 69 |
| 91 | 103 | Novel | noPerfFB | 310.630435 | 303.488889 | 92 | 90 | 70 | 60 | -10 | -2 | 78 |
| 92 | 105 | Novel | noPerfFB | 333.219780 | 316.022989 | 91 | 87 | 100 | 85 | -15 | -4 | 75 |
| 93 | 106 | Novel | noPerfFB | 314.390805 | 304.136986 | 87 | 73 | 80 | 60 | -20 | -14 | 74 |
| 94 | 107 | Novel | noPerfFB | 282.010309 | 275.268041 | 97 | 97 | 60 | 60 | 0 | 0 | 68 |
| 95 | 108 | Novel | noPerfFB | 309.443299 | 309.170213 | 97 | 94 | 75 | 100 | 25 | -3 | 72 |
| 96 | 113 | Novel | noPerfFB | 326.215054 | 304.372340 | 93 | 94 | 85 | 65 | -20 | 1 | 64 |
| 97 | 122 | Novel | noPerfFB | 312.553191 | 302.663158 | 94 | 95 | 80 | 95 | 15 | 1 | 62 |
| 98 | 123 | Novel | noPerfFB | 282.391753 | 289.638298 | 97 | 94 | 60 | 75 | 15 | -3 | 72 |
| 99 | 126 | Novel | noPerfFB | 300.440860 | 296.329670 | 93 | 91 | 65 | 70 | 5 | -2 | 68 |

Congruency refers to the within-subject Mapping factor. In the Familiar condition, Congruent means Red:NoGo and Green:Go--congruent with daily experiences, whereas in the Novel condition, these congruency mappings are arbitrary: Purple:Go, Blue:NoGo.

In [3]:

```
stacked_data=pd.melt(df, id_vars=["Subj_ID", "Stim_Cond", "FB_Cond"], value_vars=["NoGo_ACC", "NoGo_Rev_ACC"], 
                     var_name="Signal", value_name="Accuracy")
def condition_phase(x):
    if x == "NoGo_ACC":
        return "Congruent"
    elif x == "NoGo_Rev_ACC":
        return "Incongruent"
func = np.vectorize(condition_phase)
stacked_data["Congruency"] = func(stacked_data["Signal"])

stacked_data_fam = stacked_data.iloc[np.r_[0:50, 100:150]]

stacked_data_fam
```

Out[3]:

|  | Subj\_ID | Stim\_Cond | FB\_Cond | Signal | Accuracy | Congruency |
| --- | --- | --- | --- | --- | --- | --- |
| 0 | 1 | Familiar | PerfFB | NoGo\_ACC | 55 | Congruent |
| 1 | 2 | Familiar | PerfFB | NoGo\_ACC | 90 | Congruent |
| 2 | 3 | Familiar | PerfFB | NoGo\_ACC | 60 | Congruent |
| 3 | 4 | Familiar | PerfFB | NoGo\_ACC | 70 | Congruent |
| 4 | 5 | Familiar | PerfFB | NoGo\_ACC | 80 | Congruent |
| 5 | 6 | Familiar | PerfFB | NoGo\_ACC | 80 | Congruent |
| 6 | 7 | Familiar | PerfFB | NoGo\_ACC | 90 | Congruent |
| 7 | 8 | Familiar | PerfFB | NoGo\_ACC | 85 | Congruent |
| 8 | 9 | Familiar | PerfFB | NoGo\_ACC | 65 | Congruent |
| 9 | 10 | Familiar | PerfFB | NoGo\_ACC | 75 | Congruent |
| 10 | 11 | Familiar | PerfFB | NoGo\_ACC | 55 | Congruent |
| 11 | 12 | Familiar | PerfFB | NoGo\_ACC | 75 | Congruent |
| 12 | 13 | Familiar | PerfFB | NoGo\_ACC | 90 | Congruent |
| 13 | 15 | Familiar | PerfFB | NoGo\_ACC | 65 | Congruent |
| 14 | 16 | Familiar | PerfFB | NoGo\_ACC | 90 | Congruent |
| 15 | 18 | Familiar | PerfFB | NoGo\_ACC | 60 | Congruent |
| 16 | 19 | Familiar | PerfFB | NoGo\_ACC | 95 | Congruent |
| 17 | 20 | Familiar | PerfFB | NoGo\_ACC | 60 | Congruent |
| 18 | 23 | Familiar | PerfFB | NoGo\_ACC | 80 | Congruent |
| 19 | 25 | Familiar | PerfFB | NoGo\_ACC | 80 | Congruent |
| 20 | 115 | Familiar | PerfFB | NoGo\_ACC | 100 | Congruent |
| 21 | 116 | Familiar | PerfFB | NoGo\_ACC | 65 | Congruent |
| 22 | 118 | Familiar | PerfFB | NoGo\_ACC | 80 | Congruent |
| 23 | 119 | Familiar | PerfFB | NoGo\_ACC | 95 | Congruent |
| 24 | 120 | Familiar | PerfFB | NoGo\_ACC | 10 | Congruent |
| 25 | 65 | Familiar | noPerfFB | NoGo\_ACC | 55 | Congruent |
| 26 | 66 | Familiar | noPerfFB | NoGo\_ACC | 85 | Congruent |
| 27 | 68 | Familiar | noPerfFB | NoGo\_ACC | 100 | Congruent |
| 28 | 69 | Familiar | noPerfFB | NoGo\_ACC | 50 | Congruent |
| 29 | 71 | Familiar | noPerfFB | NoGo\_ACC | 75 | Congruent |
| 30 | 72 | Familiar | noPerfFB | NoGo\_ACC | 75 | Congruent |
| 31 | 73 | Familiar | noPerfFB | NoGo\_ACC | 80 | Congruent |
| 32 | 74 | Familiar | noPerfFB | NoGo\_ACC | 75 | Congruent |
| 33 | 75 | Familiar | noPerfFB | NoGo\_ACC | 95 | Congruent |
| 34 | 76 | Familiar | noPerfFB | NoGo\_ACC | 80 | Congruent |
| 35 | 77 | Familiar | noPerfFB | NoGo\_ACC | 90 | Congruent |
| 36 | 78 | Familiar | noPerfFB | NoGo\_ACC | 45 | Congruent |
| 37 | 79 | Familiar | noPerfFB | NoGo\_ACC | 95 | Congruent |
| 38 | 80 | Familiar | noPerfFB | NoGo\_ACC | 80 | Congruent |
| 39 | 82 | Familiar | noPerfFB | NoGo\_ACC | 65 | Congruent |
| 40 | 83 | Familiar | noPerfFB | NoGo\_ACC | 90 | Congruent |
| 41 | 84 | Familiar | noPerfFB | NoGo\_ACC | 85 | Congruent |
| 42 | 85 | Familiar | noPerfFB | NoGo\_ACC | 70 | Congruent |
| 43 | 86 | Familiar | noPerfFB | NoGo\_ACC | 85 | Congruent |
| 44 | 87 | Familiar | noPerfFB | NoGo\_ACC | 70 | Congruent |
| 45 | 88 | Familiar | noPerfFB | NoGo\_ACC | 100 | Congruent |
| 46 | 109 | Familiar | noPerfFB | NoGo\_ACC | 75 | Congruent |
| 47 | 110 | Familiar | noPerfFB | NoGo\_ACC | 100 | Congruent |
| 48 | 111 | Familiar | noPerfFB | NoGo\_ACC | 70 | Congruent |
| 49 | 112 | Familiar | noPerfFB | NoGo\_ACC | 85 | Congruent |
| 100 | 1 | Familiar | PerfFB | NoGo\_Rev\_ACC | 35 | Incongruent |
| 101 | 2 | Familiar | PerfFB | NoGo\_Rev\_ACC | 90 | Incongruent |
| 102 | 3 | Familiar | PerfFB | NoGo\_Rev\_ACC | 70 | Incongruent |
| 103 | 4 | Familiar | PerfFB | NoGo\_Rev\_ACC | 65 | Incongruent |
| 104 | 5 | Familiar | PerfFB | NoGo\_Rev\_ACC | 70 | Incongruent |
| 105 | 6 | Familiar | PerfFB | NoGo\_Rev\_ACC | 80 | Incongruent |
| 106 | 7 | Familiar | PerfFB | NoGo\_Rev\_ACC | 70 | Incongruent |
| 107 | 8 | Familiar | PerfFB | NoGo\_Rev\_ACC | 70 | Incongruent |
| 108 | 9 | Familiar | PerfFB | NoGo\_Rev\_ACC | 75 | Incongruent |
| 109 | 10 | Familiar | PerfFB | NoGo\_Rev\_ACC | 85 | Incongruent |
| 110 | 11 | Familiar | PerfFB | NoGo\_Rev\_ACC | 50 | Incongruent |
| 111 | 12 | Familiar | PerfFB | NoGo\_Rev\_ACC | 45 | Incongruent |
| 112 | 13 | Familiar | PerfFB | NoGo\_Rev\_ACC | 75 | Incongruent |
| 113 | 15 | Familiar | PerfFB | NoGo\_Rev\_ACC | 50 | Incongruent |
| 114 | 16 | Familiar | PerfFB | NoGo\_Rev\_ACC | 100 | Incongruent |
| 115 | 18 | Familiar | PerfFB | NoGo\_Rev\_ACC | 35 | Incongruent |
| 116 | 19 | Familiar | PerfFB | NoGo\_Rev\_ACC | 90 | Incongruent |
| 117 | 20 | Familiar | PerfFB | NoGo\_Rev\_ACC | 70 | Incongruent |
| 118 | 23 | Familiar | PerfFB | NoGo\_Rev\_ACC | 70 | Incongruent |
| 119 | 25 | Familiar | PerfFB | NoGo\_Rev\_ACC | 55 | Incongruent |
| 120 | 115 | Familiar | PerfFB | NoGo\_Rev\_ACC | 85 | Incongruent |
| 121 | 116 | Familiar | PerfFB | NoGo\_Rev\_ACC | 30 | Incongruent |
| 122 | 118 | Familiar | PerfFB | NoGo\_Rev\_ACC | 95 | Incongruent |
| 123 | 119 | Familiar | PerfFB | NoGo\_Rev\_ACC | 70 | Incongruent |
| 124 | 120 | Familiar | PerfFB | NoGo\_Rev\_ACC | 20 | Incongruent |
| 125 | 65 | Familiar | noPerfFB | NoGo\_Rev\_ACC | 80 | Incongruent |
| 126 | 66 | Familiar | noPerfFB | NoGo\_Rev\_ACC | 70 | Incongruent |
| 127 | 68 | Familiar | noPerfFB | NoGo\_Rev\_ACC | 70 | Incongruent |
| 128 | 69 | Familiar | noPerfFB | NoGo\_Rev\_ACC | 60 | Incongruent |
| 129 | 71 | Familiar | noPerfFB | NoGo\_Rev\_ACC | 85 | Incongruent |
| 130 | 72 | Familiar | noPerfFB | NoGo\_Rev\_ACC | 50 | Incongruent |
| 131 | 73 | Familiar | noPerfFB | NoGo\_Rev\_ACC | 45 | Incongruent |
| 132 | 74 | Familiar | noPerfFB | NoGo\_Rev\_ACC | 50 | Incongruent |
| 133 | 75 | Familiar | noPerfFB | NoGo\_Rev\_ACC | 95 | Incongruent |
| 134 | 76 | Familiar | noPerfFB | NoGo\_Rev\_ACC | 50 | Incongruent |
| 135 | 77 | Familiar | noPerfFB | NoGo\_Rev\_ACC | 90 | Incongruent |
| 136 | 78 | Familiar | noPerfFB | NoGo\_Rev\_ACC | 25 | Incongruent |
| 137 | 79 | Familiar | noPerfFB | NoGo\_Rev\_ACC | 90 | Incongruent |
| 138 | 80 | Familiar | noPerfFB | NoGo\_Rev\_ACC | 70 | Incongruent |
| 139 | 82 | Familiar | noPerfFB | NoGo\_Rev\_ACC | 35 | Incongruent |
| 140 | 83 | Familiar | noPerfFB | NoGo\_Rev\_ACC | 60 | Incongruent |
| 141 | 84 | Familiar | noPerfFB | NoGo\_Rev\_ACC | 45 | Incongruent |
| 142 | 85 | Familiar | noPerfFB | NoGo\_Rev\_ACC | 90 | Incongruent |
| 143 | 86 | Familiar | noPerfFB | NoGo\_Rev\_ACC | 70 | Incongruent |
| 144 | 87 | Familiar | noPerfFB | NoGo\_Rev\_ACC | 65 | Incongruent |
| 145 | 88 | Familiar | noPerfFB | NoGo\_Rev\_ACC | 100 | Incongruent |
| 146 | 109 | Familiar | noPerfFB | NoGo\_Rev\_ACC | 65 | Incongruent |
| 147 | 110 | Familiar | noPerfFB | NoGo\_Rev\_ACC | 85 | Incongruent |
| 148 | 111 | Familiar | noPerfFB | NoGo\_Rev\_ACC | 75 | Incongruent |
| 149 | 112 | Familiar | noPerfFB | NoGo\_Rev\_ACC | 80 | Incongruent |

In [4]:

```
sns.set(style="white", context="notebook", font="Times New Roman", font_scale=1.3)
ax = sns.barplot(x="FB_Cond", y="Accuracy", hue="Congruency", palette=["#ff0000", "#03d547"], ci=68, capsize=0.02, data=stacked_data_fam)
plt.title("Familiar", weight="bold", y=1.08, fontsize=20)
ax.set_ylim(50,85)
ax.set_xlabel("")
ax.set_ylabel("NoGo Accuracy (%)", weight="bold", labelpad=5, fontsize=20)
ax.set_xticklabels(["Feedback", "No Feedback"], fontsize=20)
ax.legend_.remove()
sns.despine(bottom=False)
#add sig asterisks and line
#first
x1, x2 = -0.20, 0.18
y, h, col = stacked_data_fam["Accuracy"].mean()+12.2, 0.5, "k"
plt.plot([x1, x1, x2, x2], [y, y+h, y+h, y], lw=1, c=col)
plt.text((x1+x2)*0.5, y+h-0.1, "*", ha="center", va="bottom", color=col)
#second
x3, x4 = 0.79, 1.17
y, h, col = stacked_data_fam["Accuracy"].mean()+12.2, 0.5, "k"
plt.plot([x3, x3, x4, x4], [y, y+h, y+h, y], lw=1, c=col)
plt.text((x3+x4)*0.5, y+h-0.1, "*", ha="center", va="bottom", color=col)
plt.text(-0.1, 1.1, "A", weight="bold", fontsize=25, ha="left", va="bottom", transform=ax.transAxes)
#plt.savefig("Exp4_NoGo_graph.tiff", bbox_inches="tight", dpi=1000)
plt.show()
```

In [5]:

```
stacked_data_nov = stacked_data.iloc[np.r_[50:100, 150:200]]

stacked_data_nov
```

Out[5]:

|  | Subj\_ID | Stim\_Cond | FB\_Cond | Signal | Accuracy | Congruency |
| --- | --- | --- | --- | --- | --- | --- |
| 50 | 21 | Novel | PerfFB | NoGo\_ACC | 100 | Congruent |
| 51 | 22 | Novel | PerfFB | NoGo\_ACC | 45 | Congruent |
| 52 | 24 | Novel | PerfFB | NoGo\_ACC | 40 | Congruent |
| 53 | 26 | Novel | PerfFB | NoGo\_ACC | 90 | Congruent |
| 54 | 27 | Novel | PerfFB | NoGo\_ACC | 90 | Congruent |
| 55 | 28 | Novel | PerfFB | NoGo\_ACC | 75 | Congruent |
| 56 | 29 | Novel | PerfFB | NoGo\_ACC | 45 | Congruent |
| 57 | 30 | Novel | PerfFB | NoGo\_ACC | 70 | Congruent |
| 58 | 31 | Novel | PerfFB | NoGo\_ACC | 85 | Congruent |
| 59 | 32 | Novel | PerfFB | NoGo\_ACC | 85 | Congruent |
| 60 | 33 | Novel | PerfFB | NoGo\_ACC | 55 | Congruent |
| 61 | 34 | Novel | PerfFB | NoGo\_ACC | 40 | Congruent |
| 62 | 35 | Novel | PerfFB | NoGo\_ACC | 50 | Congruent |
| 63 | 36 | Novel | PerfFB | NoGo\_ACC | 85 | Congruent |
| 64 | 37 | Novel | PerfFB | NoGo\_ACC | 70 | Congruent |
| 65 | 38 | Novel | PerfFB | NoGo\_ACC | 95 | Congruent |
| 66 | 39 | Novel | PerfFB | NoGo\_ACC | 60 | Congruent |
| 67 | 40 | Novel | PerfFB | NoGo\_ACC | 90 | Congruent |
| 68 | 41 | Novel | PerfFB | NoGo\_ACC | 60 | Congruent |
| 69 | 104 | Novel | PerfFB | NoGo\_ACC | 80 | Congruent |
| 70 | 117 | Novel | PerfFB | NoGo\_ACC | 55 | Congruent |
| 71 | 121 | Novel | PerfFB | NoGo\_ACC | 70 | Congruent |
| 72 | 124 | Novel | PerfFB | NoGo\_ACC | 40 | Congruent |
| 73 | 125 | Novel | PerfFB | NoGo\_ACC | 90 | Congruent |
| 74 | 127 | Novel | PerfFB | NoGo\_ACC | 85 | Congruent |
| 75 | 67 | Novel | noPerfFB | NoGo\_ACC | 70 | Congruent |
| 76 | 70 | Novel | noPerfFB | NoGo\_ACC | 50 | Congruent |
| 77 | 81 | Novel | noPerfFB | NoGo\_ACC | 80 | Congruent |
| 78 | 89 | Novel | noPerfFB | NoGo\_ACC | 90 | Congruent |
| 79 | 90 | Novel | noPerfFB | NoGo\_ACC | 40 | Congruent |
| 80 | 91 | Novel | noPerfFB | NoGo\_ACC | 80 | Congruent |
| 81 | 92 | Novel | noPerfFB | NoGo\_ACC | 100 | Congruent |
| 82 | 94 | Novel | noPerfFB | NoGo\_ACC | 80 | Congruent |
| 83 | 95 | Novel | noPerfFB | NoGo\_ACC | 75 | Congruent |
| 84 | 96 | Novel | noPerfFB | NoGo\_ACC | 70 | Congruent |
| 85 | 97 | Novel | noPerfFB | NoGo\_ACC | 70 | Congruent |
| 86 | 98 | Novel | noPerfFB | NoGo\_ACC | 75 | Congruent |
| 87 | 99 | Novel | noPerfFB | NoGo\_ACC | 85 | Congruent |
| 88 | 100 | Novel | noPerfFB | NoGo\_ACC | 65 | Congruent |
| 89 | 101 | Novel | noPerfFB | NoGo\_ACC | 75 | Congruent |
| 90 | 102 | Novel | noPerfFB | NoGo\_ACC | 80 | Congruent |
| 91 | 103 | Novel | noPerfFB | NoGo\_ACC | 70 | Congruent |
| 92 | 105 | Novel | noPerfFB | NoGo\_ACC | 100 | Congruent |
| 93 | 106 | Novel | noPerfFB | NoGo\_ACC | 80 | Congruent |
| 94 | 107 | Novel | noPerfFB | NoGo\_ACC | 60 | Congruent |
| 95 | 108 | Novel | noPerfFB | NoGo\_ACC | 75 | Congruent |
| 96 | 113 | Novel | noPerfFB | NoGo\_ACC | 85 | Congruent |
| 97 | 122 | Novel | noPerfFB | NoGo\_ACC | 80 | Congruent |
| 98 | 123 | Novel | noPerfFB | NoGo\_ACC | 60 | Congruent |
| 99 | 126 | Novel | noPerfFB | NoGo\_ACC | 65 | Congruent |
| 150 | 21 | Novel | PerfFB | NoGo\_Rev\_ACC | 100 | Incongruent |
| 151 | 22 | Novel | PerfFB | NoGo\_Rev\_ACC | 95 | Incongruent |
| 152 | 24 | Novel | PerfFB | NoGo\_Rev\_ACC | 75 | Incongruent |
| 153 | 26 | Novel | PerfFB | NoGo\_Rev\_ACC | 85 | Incongruent |
| 154 | 27 | Novel | PerfFB | NoGo\_Rev\_ACC | 95 | Incongruent |
| 155 | 28 | Novel | PerfFB | NoGo\_Rev\_ACC | 80 | Incongruent |
| 156 | 29 | Novel | PerfFB | NoGo\_Rev\_ACC | 70 | Incongruent |
| 157 | 30 | Novel | PerfFB | NoGo\_Rev\_ACC | 75 | Incongruent |
| 158 | 31 | Novel | PerfFB | NoGo\_Rev\_ACC | 85 | Incongruent |
| 159 | 32 | Novel | PerfFB | NoGo\_Rev\_ACC | 75 | Incongruent |
| 160 | 33 | Novel | PerfFB | NoGo\_Rev\_ACC | 65 | Incongruent |
| 161 | 34 | Novel | PerfFB | NoGo\_Rev\_ACC | 60 | Incongruent |
| 162 | 35 | Novel | PerfFB | NoGo\_Rev\_ACC | 75 | Incongruent |
| 163 | 36 | Novel | PerfFB | NoGo\_Rev\_ACC | 85 | Incongruent |
| 164 | 37 | Novel | PerfFB | NoGo\_Rev\_ACC | 60 | Incongruent |
| 165 | 38 | Novel | PerfFB | NoGo\_Rev\_ACC | 55 | Incongruent |
| 166 | 39 | Novel | PerfFB | NoGo\_Rev\_ACC | 50 | Incongruent |
| 167 | 40 | Novel | PerfFB | NoGo\_Rev\_ACC | 95 | Incongruent |
| 168 | 41 | Novel | PerfFB | NoGo\_Rev\_ACC | 85 | Incongruent |
| 169 | 104 | Novel | PerfFB | NoGo\_Rev\_ACC | 95 | Incongruent |
| 170 | 117 | Novel | PerfFB | NoGo\_Rev\_ACC | 45 | Incongruent |
| 171 | 121 | Novel | PerfFB | NoGo\_Rev\_ACC | 75 | Incongruent |
| 172 | 124 | Novel | PerfFB | NoGo\_Rev\_ACC | 60 | Incongruent |
| 173 | 125 | Novel | PerfFB | NoGo\_Rev\_ACC | 90 | Incongruent |
| 174 | 127 | Novel | PerfFB | NoGo\_Rev\_ACC | 70 | Incongruent |
| 175 | 67 | Novel | noPerfFB | NoGo\_Rev\_ACC | 80 | Incongruent |
| 176 | 70 | Novel | noPerfFB | NoGo\_Rev\_ACC | 65 | Incongruent |
| 177 | 81 | Novel | noPerfFB | NoGo\_Rev\_ACC | 75 | Incongruent |
| 178 | 89 | Novel | noPerfFB | NoGo\_Rev\_ACC | 90 | Incongruent |
| 179 | 90 | Novel | noPerfFB | NoGo\_Rev\_ACC | 35 | Incongruent |
| 180 | 91 | Novel | noPerfFB | NoGo\_Rev\_ACC | 75 | Incongruent |
| 181 | 92 | Novel | noPerfFB | NoGo\_Rev\_ACC | 95 | Incongruent |
| 182 | 94 | Novel | noPerfFB | NoGo\_Rev\_ACC | 90 | Incongruent |
| 183 | 95 | Novel | noPerfFB | NoGo\_Rev\_ACC | 55 | Incongruent |
| 184 | 96 | Novel | noPerfFB | NoGo\_Rev\_ACC | 50 | Incongruent |
| 185 | 97 | Novel | noPerfFB | NoGo\_Rev\_ACC | 65 | Incongruent |
| 186 | 98 | Novel | noPerfFB | NoGo\_Rev\_ACC | 85 | Incongruent |
| 187 | 99 | Novel | noPerfFB | NoGo\_Rev\_ACC | 95 | Incongruent |
| 188 | 100 | Novel | noPerfFB | NoGo\_Rev\_ACC | 60 | Incongruent |
| 189 | 101 | Novel | noPerfFB | NoGo\_Rev\_ACC | 50 | Incongruent |
| 190 | 102 | Novel | noPerfFB | NoGo\_Rev\_ACC | 65 | Incongruent |
| 191 | 103 | Novel | noPerfFB | NoGo\_Rev\_ACC | 60 | Incongruent |
| 192 | 105 | Novel | noPerfFB | NoGo\_Rev\_ACC | 85 | Incongruent |
| 193 | 106 | Novel | noPerfFB | NoGo\_Rev\_ACC | 60 | Incongruent |
| 194 | 107 | Novel | noPerfFB | NoGo\_Rev\_ACC | 60 | Incongruent |
| 195 | 108 | Novel | noPerfFB | NoGo\_Rev\_ACC | 100 | Incongruent |
| 196 | 113 | Novel | noPerfFB | NoGo\_Rev\_ACC | 65 | Incongruent |
| 197 | 122 | Novel | noPerfFB | NoGo\_Rev\_ACC | 95 | Incongruent |
| 198 | 123 | Novel | noPerfFB | NoGo\_Rev\_ACC | 75 | Incongruent |
| 199 | 126 | Novel | noPerfFB | NoGo\_Rev\_ACC | 70 | Incongruent |

In [6]:

```
sns.set(style="white", context="notebook", font="Times New Roman", font_scale=1.3)
ax = sns.barplot(x="FB_Cond", y="Accuracy", hue="Congruency", palette=["#1d47f5", "#d12fdf"], ci=68, capsize=0.02, data=stacked_data_nov)
plt.title("Novel", weight="bold", y=1.08, fontsize=20)
ax.set_ylim(50,85)
ax.set_xlabel("")
ax.set_ylabel("NoGo Accuracy (%)", weight="bold", labelpad=5, fontsize=20)
ax.set_xticklabels(["Feedback", "No Feedback"], fontsize=20)
ax.legend_.remove()
sns.despine(bottom=False)
plt.text(-0.1, 1.1, "B", weight="bold", fontsize=25, ha="left", va="bottom", transform=ax.transAxes)
#plt.savefig("Exp4_NoGo_graph_Nov.tiff", bbox_inches="tight", dpi=1000)
plt.show()
```

Go Accuracy graphs

In [7]:

```
stacked_data_go=pd.melt(df, id_vars=["Subj_ID", "Stim_Cond", "FB_Cond"], value_vars=["Go_ACC", "Go_Rev_ACC"], 
                     var_name="Signal", value_name="Accuracy")
def condition_phase(x):
    if x == "Go_ACC":
        return "Congruent"
    elif x == "Go_Rev_ACC":
        return "Incongruent"
func = np.vectorize(condition_phase)
stacked_data_go["Congruency"] = func(stacked_data_go["Signal"])

stacked_data_fam_go = stacked_data_go.iloc[np.r_[0:50, 100:150]]

stacked_data_fam_go
```

Out[7]:

|  | Subj\_ID | Stim\_Cond | FB\_Cond | Signal | Accuracy | Congruency |
| --- | --- | --- | --- | --- | --- | --- |
| 0 | 1 | Familiar | PerfFB | Go\_ACC | 87 | Congruent |
| 1 | 2 | Familiar | PerfFB | Go\_ACC | 97 | Congruent |
| 2 | 3 | Familiar | PerfFB | Go\_ACC | 93 | Congruent |
| 3 | 4 | Familiar | PerfFB | Go\_ACC | 92 | Congruent |
| 4 | 5 | Familiar | PerfFB | Go\_ACC | 94 | Congruent |
| 5 | 6 | Familiar | PerfFB | Go\_ACC | 97 | Congruent |
| 6 | 7 | Familiar | PerfFB | Go\_ACC | 94 | Congruent |
| 7 | 8 | Familiar | PerfFB | Go\_ACC | 75 | Congruent |
| 8 | 9 | Familiar | PerfFB | Go\_ACC | 88 | Congruent |
| 9 | 10 | Familiar | PerfFB | Go\_ACC | 95 | Congruent |
| 10 | 11 | Familiar | PerfFB | Go\_ACC | 97 | Congruent |
| 11 | 12 | Familiar | PerfFB | Go\_ACC | 93 | Congruent |
| 12 | 13 | Familiar | PerfFB | Go\_ACC | 97 | Congruent |
| 13 | 15 | Familiar | PerfFB | Go\_ACC | 95 | Congruent |
| 14 | 16 | Familiar | PerfFB | Go\_ACC | 92 | Congruent |
| 15 | 18 | Familiar | PerfFB | Go\_ACC | 94 | Congruent |
| 16 | 19 | Familiar | PerfFB | Go\_ACC | 90 | Congruent |
| 17 | 20 | Familiar | PerfFB | Go\_ACC | 91 | Congruent |
| 18 | 23 | Familiar | PerfFB | Go\_ACC | 87 | Congruent |
| 19 | 25 | Familiar | PerfFB | Go\_ACC | 84 | Congruent |
| 20 | 115 | Familiar | PerfFB | Go\_ACC | 92 | Congruent |
| 21 | 116 | Familiar | PerfFB | Go\_ACC | 82 | Congruent |
| 22 | 118 | Familiar | PerfFB | Go\_ACC | 91 | Congruent |
| 23 | 119 | Familiar | PerfFB | Go\_ACC | 96 | Congruent |
| 24 | 120 | Familiar | PerfFB | Go\_ACC | 90 | Congruent |
| 25 | 65 | Familiar | noPerfFB | Go\_ACC | 82 | Congruent |
| 26 | 66 | Familiar | noPerfFB | Go\_ACC | 94 | Congruent |
| 27 | 68 | Familiar | noPerfFB | Go\_ACC | 98 | Congruent |
| 28 | 69 | Familiar | noPerfFB | Go\_ACC | 89 | Congruent |
| 29 | 71 | Familiar | noPerfFB | Go\_ACC | 94 | Congruent |
| 30 | 72 | Familiar | noPerfFB | Go\_ACC | 96 | Congruent |
| 31 | 73 | Familiar | noPerfFB | Go\_ACC | 93 | Congruent |
| 32 | 74 | Familiar | noPerfFB | Go\_ACC | 87 | Congruent |
| 33 | 75 | Familiar | noPerfFB | Go\_ACC | 96 | Congruent |
| 34 | 76 | Familiar | noPerfFB | Go\_ACC | 97 | Congruent |
| 35 | 77 | Familiar | noPerfFB | Go\_ACC | 86 | Congruent |
| 36 | 78 | Familiar | noPerfFB | Go\_ACC | 94 | Congruent |
| 37 | 79 | Familiar | noPerfFB | Go\_ACC | 97 | Congruent |
| 38 | 80 | Familiar | noPerfFB | Go\_ACC | 96 | Congruent |
| 39 | 82 | Familiar | noPerfFB | Go\_ACC | 91 | Congruent |
| 40 | 83 | Familiar | noPerfFB | Go\_ACC | 94 | Congruent |
| 41 | 84 | Familiar | noPerfFB | Go\_ACC | 95 | Congruent |
| 42 | 85 | Familiar | noPerfFB | Go\_ACC | 97 | Congruent |
| 43 | 86 | Familiar | noPerfFB | Go\_ACC | 89 | Congruent |
| 44 | 87 | Familiar | noPerfFB | Go\_ACC | 90 | Congruent |
| 45 | 88 | Familiar | noPerfFB | Go\_ACC | 98 | Congruent |
| 46 | 109 | Familiar | noPerfFB | Go\_ACC | 96 | Congruent |
| 47 | 110 | Familiar | noPerfFB | Go\_ACC | 95 | Congruent |
| 48 | 111 | Familiar | noPerfFB | Go\_ACC | 93 | Congruent |
| 49 | 112 | Familiar | noPerfFB | Go\_ACC | 96 | Congruent |
| 100 | 1 | Familiar | PerfFB | Go\_Rev\_ACC | 87 | Incongruent |
| 101 | 2 | Familiar | PerfFB | Go\_Rev\_ACC | 97 | Incongruent |
| 102 | 3 | Familiar | PerfFB | Go\_Rev\_ACC | 95 | Incongruent |
| 103 | 4 | Familiar | PerfFB | Go\_Rev\_ACC | 90 | Incongruent |
| 104 | 5 | Familiar | PerfFB | Go\_Rev\_ACC | 95 | Incongruent |
| 105 | 6 | Familiar | PerfFB | Go\_Rev\_ACC | 88 | Incongruent |
| 106 | 7 | Familiar | PerfFB | Go\_Rev\_ACC | 92 | Incongruent |
| 107 | 8 | Familiar | PerfFB | Go\_Rev\_ACC | 79 | Incongruent |
| 108 | 9 | Familiar | PerfFB | Go\_Rev\_ACC | 78 | Incongruent |
| 109 | 10 | Familiar | PerfFB | Go\_Rev\_ACC | 97 | Incongruent |
| 110 | 11 | Familiar | PerfFB | Go\_Rev\_ACC | 93 | Incongruent |
| 111 | 12 | Familiar | PerfFB | Go\_Rev\_ACC | 89 | Incongruent |
| 112 | 13 | Familiar | PerfFB | Go\_Rev\_ACC | 95 | Incongruent |
| 113 | 15 | Familiar | PerfFB | Go\_Rev\_ACC | 96 | Incongruent |
| 114 | 16 | Familiar | PerfFB | Go\_Rev\_ACC | 94 | Incongruent |
| 115 | 18 | Familiar | PerfFB | Go\_Rev\_ACC | 90 | Incongruent |
| 116 | 19 | Familiar | PerfFB | Go\_Rev\_ACC | 96 | Incongruent |
| 117 | 20 | Familiar | PerfFB | Go\_Rev\_ACC | 90 | Incongruent |
| 118 | 23 | Familiar | PerfFB | Go\_Rev\_ACC | 92 | Incongruent |
| 119 | 25 | Familiar | PerfFB | Go\_Rev\_ACC | 89 | Incongruent |
| 120 | 115 | Familiar | PerfFB | Go\_Rev\_ACC | 92 | Incongruent |
| 121 | 116 | Familiar | PerfFB | Go\_Rev\_ACC | 80 | Incongruent |
| 122 | 118 | Familiar | PerfFB | Go\_Rev\_ACC | 88 | Incongruent |
| 123 | 119 | Familiar | PerfFB | Go\_Rev\_ACC | 94 | Incongruent |
| 124 | 120 | Familiar | PerfFB | Go\_Rev\_ACC | 84 | Incongruent |
| 125 | 65 | Familiar | noPerfFB | Go\_Rev\_ACC | 89 | Incongruent |
| 126 | 66 | Familiar | noPerfFB | Go\_Rev\_ACC | 87 | Incongruent |
| 127 | 68 | Familiar | noPerfFB | Go\_Rev\_ACC | 87 | Incongruent |
| 128 | 69 | Familiar | noPerfFB | Go\_Rev\_ACC | 65 | Incongruent |
| 129 | 71 | Familiar | noPerfFB | Go\_Rev\_ACC | 94 | Incongruent |
| 130 | 72 | Familiar | noPerfFB | Go\_Rev\_ACC | 93 | Incongruent |
| 131 | 73 | Familiar | noPerfFB | Go\_Rev\_ACC | 83 | Incongruent |
| 132 | 74 | Familiar | noPerfFB | Go\_Rev\_ACC | 93 | Incongruent |
| 133 | 75 | Familiar | noPerfFB | Go\_Rev\_ACC | 91 | Incongruent |
| 134 | 76 | Familiar | noPerfFB | Go\_Rev\_ACC | 86 | Incongruent |
| 135 | 77 | Familiar | noPerfFB | Go\_Rev\_ACC | 51 | Incongruent |
| 136 | 78 | Familiar | noPerfFB | Go\_Rev\_ACC | 91 | Incongruent |
| 137 | 79 | Familiar | noPerfFB | Go\_Rev\_ACC | 96 | Incongruent |
| 138 | 80 | Familiar | noPerfFB | Go\_Rev\_ACC | 93 | Incongruent |
| 139 | 82 | Familiar | noPerfFB | Go\_Rev\_ACC | 89 | Incongruent |
| 140 | 83 | Familiar | noPerfFB | Go\_Rev\_ACC | 93 | Incongruent |
| 141 | 84 | Familiar | noPerfFB | Go\_Rev\_ACC | 87 | Incongruent |
| 142 | 85 | Familiar | noPerfFB | Go\_Rev\_ACC | 96 | Incongruent |
| 143 | 86 | Familiar | noPerfFB | Go\_Rev\_ACC | 83 | Incongruent |
| 144 | 87 | Familiar | noPerfFB | Go\_Rev\_ACC | 89 | Incongruent |
| 145 | 88 | Familiar | noPerfFB | Go\_Rev\_ACC | 94 | Incongruent |
| 146 | 109 | Familiar | noPerfFB | Go\_Rev\_ACC | 94 | Incongruent |
| 147 | 110 | Familiar | noPerfFB | Go\_Rev\_ACC | 93 | Incongruent |
| 148 | 111 | Familiar | noPerfFB | Go\_Rev\_ACC | 87 | Incongruent |
| 149 | 112 | Familiar | noPerfFB | Go\_Rev\_ACC | 91 | Incongruent |

In [8]:

```
sns.set(style="white", context="notebook", font="Times New Roman", font_scale=1.3)
ax = sns.barplot(x="FB_Cond", y="Accuracy", hue="Congruency", palette=["#03d547", "#ff0000"], ci=68, capsize=0.02, data=stacked_data_fam_go)
plt.title("Familiar", weight="bold", y=1.08, fontsize=20)
ax.set_ylim(50,100)
ax.set_xlabel("")
ax.set_ylabel("Go Accuracy (%)", weight="bold", labelpad=5, fontsize=20)
ax.set_xticklabels(["Feedback", "No Feedback"], fontsize=20)
ax.legend_.remove()
sns.despine(bottom=False)
#add sig asterisks and line
x3, x4 = 0.79, 1.17
y, h, col = stacked_data_fam_go["Accuracy"].mean()+8, 0.5, "k"
plt.plot([x3, x3, x4, x4], [y, y+h, y+h, y], lw=1, c=col)
plt.text((x3+x4)*0.5, y+h-0.1, "*", ha="center", va="bottom", color=col)
plt.text(-0.1, 1.1, "A", weight="bold", fontsize=25, ha="left", va="bottom", transform=ax.transAxes)
#plt.savefig("Exp4_NoGo_graph_Fam_Go.tiff", bbox_inches="tight", dpi=1000)
plt.show()
```

In [9]:

```
stacked_data_go=pd.melt(df, id_vars=["Subj_ID", "Stim_Cond", "FB_Cond"], value_vars=["Go_ACC", "Go_Rev_ACC"], 
                     var_name="Signal", value_name="Accuracy")
def condition_phase(x):
    if x == "Go_ACC":
        return "Congruent"
    elif x == "Go_Rev_ACC":
        return "Incongruent"
func = np.vectorize(condition_phase)
stacked_data_go["Congruency"] = func(stacked_data_go["Signal"])

stacked_data_nov_go = stacked_data_go.iloc[np.r_[50:100, 150:200]]

stacked_data_nov_go
```

Out[9]:

|  | Subj\_ID | Stim\_Cond | FB\_Cond | Signal | Accuracy | Congruency |
| --- | --- | --- | --- | --- | --- | --- |
| 50 | 21 | Novel | PerfFB | Go\_ACC | 94 | Congruent |
| 51 | 22 | Novel | PerfFB | Go\_ACC | 90 | Congruent |
| 52 | 24 | Novel | PerfFB | Go\_ACC | 83 | Congruent |
| 53 | 26 | Novel | PerfFB | Go\_ACC | 96 | Congruent |
| 54 | 27 | Novel | PerfFB | Go\_ACC | 97 | Congruent |
| 55 | 28 | Novel | PerfFB | Go\_ACC | 88 | Congruent |
| 56 | 29 | Novel | PerfFB | Go\_ACC | 89 | Congruent |
| 57 | 30 | Novel | PerfFB | Go\_ACC | 85 | Congruent |
| 58 | 31 | Novel | PerfFB | Go\_ACC | 93 | Congruent |
| 59 | 32 | Novel | PerfFB | Go\_ACC | 88 | Congruent |
| 60 | 33 | Novel | PerfFB | Go\_ACC | 89 | Congruent |
| 61 | 34 | Novel | PerfFB | Go\_ACC | 81 | Congruent |
| 62 | 35 | Novel | PerfFB | Go\_ACC | 77 | Congruent |
| 63 | 36 | Novel | PerfFB | Go\_ACC | 87 | Congruent |
| 64 | 37 | Novel | PerfFB | Go\_ACC | 90 | Congruent |
| 65 | 38 | Novel | PerfFB | Go\_ACC | 87 | Congruent |
| 66 | 39 | Novel | PerfFB | Go\_ACC | 84 | Congruent |
| 67 | 40 | Novel | PerfFB | Go\_ACC | 95 | Congruent |
| 68 | 41 | Novel | PerfFB | Go\_ACC | 85 | Congruent |
| 69 | 104 | Novel | PerfFB | Go\_ACC | 90 | Congruent |
| 70 | 117 | Novel | PerfFB | Go\_ACC | 71 | Congruent |
| 71 | 121 | Novel | PerfFB | Go\_ACC | 91 | Congruent |
| 72 | 124 | Novel | PerfFB | Go\_ACC | 88 | Congruent |
| 73 | 125 | Novel | PerfFB | Go\_ACC | 94 | Congruent |
| 74 | 127 | Novel | PerfFB | Go\_ACC | 89 | Congruent |
| 75 | 67 | Novel | noPerfFB | Go\_ACC | 79 | Congruent |
| 76 | 70 | Novel | noPerfFB | Go\_ACC | 96 | Congruent |
| 77 | 81 | Novel | noPerfFB | Go\_ACC | 97 | Congruent |
| 78 | 89 | Novel | noPerfFB | Go\_ACC | 96 | Congruent |
| 79 | 90 | Novel | noPerfFB | Go\_ACC | 89 | Congruent |
| 80 | 91 | Novel | noPerfFB | Go\_ACC | 92 | Congruent |
| 81 | 92 | Novel | noPerfFB | Go\_ACC | 93 | Congruent |
| 82 | 94 | Novel | noPerfFB | Go\_ACC | 92 | Congruent |
| 83 | 95 | Novel | noPerfFB | Go\_ACC | 90 | Congruent |
| 84 | 96 | Novel | noPerfFB | Go\_ACC | 93 | Congruent |
| 85 | 97 | Novel | noPerfFB | Go\_ACC | 92 | Congruent |
| 86 | 98 | Novel | noPerfFB | Go\_ACC | 84 | Congruent |
| 87 | 99 | Novel | noPerfFB | Go\_ACC | 93 | Congruent |
| 88 | 100 | Novel | noPerfFB | Go\_ACC | 90 | Congruent |
| 89 | 101 | Novel | noPerfFB | Go\_ACC | 97 | Congruent |
| 90 | 102 | Novel | noPerfFB | Go\_ACC | 88 | Congruent |
| 91 | 103 | Novel | noPerfFB | Go\_ACC | 92 | Congruent |
| 92 | 105 | Novel | noPerfFB | Go\_ACC | 91 | Congruent |
| 93 | 106 | Novel | noPerfFB | Go\_ACC | 87 | Congruent |
| 94 | 107 | Novel | noPerfFB | Go\_ACC | 97 | Congruent |
| 95 | 108 | Novel | noPerfFB | Go\_ACC | 97 | Congruent |
| 96 | 113 | Novel | noPerfFB | Go\_ACC | 93 | Congruent |
| 97 | 122 | Novel | noPerfFB | Go\_ACC | 94 | Congruent |
| 98 | 123 | Novel | noPerfFB | Go\_ACC | 97 | Congruent |
| 99 | 126 | Novel | noPerfFB | Go\_ACC | 93 | Congruent |
| 150 | 21 | Novel | PerfFB | Go\_Rev\_ACC | 95 | Incongruent |
| 151 | 22 | Novel | PerfFB | Go\_Rev\_ACC | 97 | Incongruent |
| 152 | 24 | Novel | PerfFB | Go\_Rev\_ACC | 88 | Incongruent |
| 153 | 26 | Novel | PerfFB | Go\_Rev\_ACC | 96 | Incongruent |
| 154 | 27 | Novel | PerfFB | Go\_Rev\_ACC | 97 | Incongruent |
| 155 | 28 | Novel | PerfFB | Go\_Rev\_ACC | 88 | Incongruent |
| 156 | 29 | Novel | PerfFB | Go\_Rev\_ACC | 94 | Incongruent |
| 157 | 30 | Novel | PerfFB | Go\_Rev\_ACC | 88 | Incongruent |
| 158 | 31 | Novel | PerfFB | Go\_Rev\_ACC | 96 | Incongruent |
| 159 | 32 | Novel | PerfFB | Go\_Rev\_ACC | 91 | Incongruent |
| 160 | 33 | Novel | PerfFB | Go\_Rev\_ACC | 93 | Incongruent |
| 161 | 34 | Novel | PerfFB | Go\_Rev\_ACC | 93 | Incongruent |
| 162 | 35 | Novel | PerfFB | Go\_Rev\_ACC | 82 | Incongruent |
| 163 | 36 | Novel | PerfFB | Go\_Rev\_ACC | 88 | Incongruent |
| 164 | 37 | Novel | PerfFB | Go\_Rev\_ACC | 90 | Incongruent |
| 165 | 38 | Novel | PerfFB | Go\_Rev\_ACC | 84 | Incongruent |
| 166 | 39 | Novel | PerfFB | Go\_Rev\_ACC | 84 | Incongruent |
| 167 | 40 | Novel | PerfFB | Go\_Rev\_ACC | 96 | Incongruent |
| 168 | 41 | Novel | PerfFB | Go\_Rev\_ACC | 89 | Incongruent |
| 169 | 104 | Novel | PerfFB | Go\_Rev\_ACC | 91 | Incongruent |
| 170 | 117 | Novel | PerfFB | Go\_Rev\_ACC | 71 | Incongruent |
| 171 | 121 | Novel | PerfFB | Go\_Rev\_ACC | 84 | Incongruent |
| 172 | 124 | Novel | PerfFB | Go\_Rev\_ACC | 96 | Incongruent |
| 173 | 125 | Novel | PerfFB | Go\_Rev\_ACC | 94 | Incongruent |
| 174 | 127 | Novel | PerfFB | Go\_Rev\_ACC | 84 | Incongruent |
| 175 | 67 | Novel | noPerfFB | Go\_Rev\_ACC | 63 | Incongruent |
| 176 | 70 | Novel | noPerfFB | Go\_Rev\_ACC | 96 | Incongruent |
| 177 | 81 | Novel | noPerfFB | Go\_Rev\_ACC | 95 | Incongruent |
| 178 | 89 | Novel | noPerfFB | Go\_Rev\_ACC | 98 | Incongruent |
| 179 | 90 | Novel | noPerfFB | Go\_Rev\_ACC | 94 | Incongruent |
| 180 | 91 | Novel | noPerfFB | Go\_Rev\_ACC | 97 | Incongruent |
| 181 | 92 | Novel | noPerfFB | Go\_Rev\_ACC | 99 | Incongruent |
| 182 | 94 | Novel | noPerfFB | Go\_Rev\_ACC | 90 | Incongruent |
| 183 | 95 | Novel | noPerfFB | Go\_Rev\_ACC | 87 | Incongruent |
| 184 | 96 | Novel | noPerfFB | Go\_Rev\_ACC | 95 | Incongruent |
| 185 | 97 | Novel | noPerfFB | Go\_Rev\_ACC | 93 | Incongruent |
| 186 | 98 | Novel | noPerfFB | Go\_Rev\_ACC | 93 | Incongruent |
| 187 | 99 | Novel | noPerfFB | Go\_Rev\_ACC | 96 | Incongruent |
| 188 | 100 | Novel | noPerfFB | Go\_Rev\_ACC | 94 | Incongruent |
| 189 | 101 | Novel | noPerfFB | Go\_Rev\_ACC | 95 | Incongruent |
| 190 | 102 | Novel | noPerfFB | Go\_Rev\_ACC | 93 | Incongruent |
| 191 | 103 | Novel | noPerfFB | Go\_Rev\_ACC | 90 | Incongruent |
| 192 | 105 | Novel | noPerfFB | Go\_Rev\_ACC | 87 | Incongruent |
| 193 | 106 | Novel | noPerfFB | Go\_Rev\_ACC | 73 | Incongruent |
| 194 | 107 | Novel | noPerfFB | Go\_Rev\_ACC | 97 | Incongruent |
| 195 | 108 | Novel | noPerfFB | Go\_Rev\_ACC | 94 | Incongruent |
| 196 | 113 | Novel | noPerfFB | Go\_Rev\_ACC | 94 | Incongruent |
| 197 | 122 | Novel | noPerfFB | Go\_Rev\_ACC | 95 | Incongruent |
| 198 | 123 | Novel | noPerfFB | Go\_Rev\_ACC | 94 | Incongruent |
| 199 | 126 | Novel | noPerfFB | Go\_Rev\_ACC | 91 | Incongruent |

In [10]:

```
sns.set(style="white", context="notebook", font="Times New Roman", font_scale=1.3)
ax = sns.barplot(x="FB_Cond", y="Accuracy", hue="Congruency", palette=["#d12fdf", "#1d47f5"], ci=68, capsize=0.02, data=stacked_data_nov_go)
plt.title("Novel", weight="bold", y=1.08, fontsize=20)
ax.set_ylim(50,100)
ax.set_xlabel("")
ax.set_ylabel("Go Accuracy (%)", weight="bold", labelpad=5, fontsize=20)
ax.set_xticklabels(["Feedback", "No Feedback"], fontsize=20)
ax.legend_.remove()
sns.despine(bottom=False)
plt.text(-0.1, 1.1, "B", weight="bold", fontsize=25, ha="left", va="bottom", transform=ax.transAxes)
#plt.savefig("Exp4_NoGo_graph_Nov_Go.tiff", bbox_inches="tight", dpi=1000)
plt.show()
```
