## Supplementary material for "Demonstrating and disrupting well-learned habits": Datasets and plotting scripts: Experiment3_Python_html_Output_corr.html

Exp7\_Graphs


In [1]:

```
#import all packages
import pandas as pd
import matplotlib as mpl
import matplotlib.pyplot as plt
import seaborn as sns
import numpy as np
from scipy import stats
from matplotlib.patches import Patch
%matplotlib inline
```

In [2]:

```
xls=pd.ExcelFile("MonPerfFB_Data.xlsx")
df=pd.read_excel(xls, "Exp7Data")
pd.set_option("display.max_rows", 500)
df
```

Out[2]:

|  | Subj\_ID | Stim\_Cond | FB\_Cond | Go\_RT | Go\_Rev\_RT | Go\_ACC | Go\_Rev\_ACC | NoGo\_ACC | NoGo\_Rev\_ACC | NoGo\_Diff | Go\_Diff | BIS |
| --- | --- | --- | --- | --- | --- | --- | --- | --- | --- | --- | --- | --- |
| 0 | 1 | Familiar | MonPerfFB | 286.824742 | 284.242105 | 97 | 95 | 90 | 95 | 5 | -2 | 78.0 |
| 1 | 2 | Familiar | MonPerfFB | 295.541667 | 283.655914 | 96 | 93 | 100 | 75 | -25 | -3 | 67.0 |
| 2 | 3 | Familiar | MonPerfFB | 307.096774 | 310.365591 | 93 | 93 | 65 | 85 | 20 | 0 | 67.0 |
| 3 | 4 | Familiar | MonPerfFB | 277.136842 | 277.041667 | 95 | 96 | 70 | 60 | -10 | 1 | 73.0 |
| 4 | 5 | Familiar | MonPerfFB | 299.291667 | 295.156250 | 96 | 96 | 95 | 90 | -5 | 0 | 70.0 |
| 5 | 6 | Familiar | MonPerfFB | 303.882979 | 286.051546 | 94 | 97 | 85 | 85 | 0 | 3 | 79.0 |
| 6 | 7 | Familiar | MonPerfFB | 304.055556 | 283.477778 | 90 | 90 | 50 | 35 | -15 | 0 | 71.0 |
| 7 | 8 | Familiar | MonPerfFB | 270.040816 | 287.893617 | 98 | 94 | 100 | 95 | -5 | -4 | 72.0 |
| 8 | 9 | Familiar | MonPerfFB | 316.455556 | 308.741935 | 90 | 93 | 80 | 85 | 5 | 3 | 70.0 |
| 9 | 10 | Familiar | MonPerfFB | 297.546392 | 298.390000 | 97 | 100 | 100 | 95 | -5 | 3 | 80.0 |
| 10 | 11 | Familiar | MonPerfFB | 283.688172 | 292.378947 | 93 | 95 | 50 | 65 | 15 | 2 | 72.0 |
| 11 | 12 | Familiar | MonPerfFB | 313.100000 | 289.054348 | 90 | 92 | 60 | 40 | -20 | 2 | 85.0 |
| 12 | 13 | Familiar | MonPerfFB | 319.422680 | 307.880435 | 97 | 92 | 100 | 75 | -25 | -5 | 78.0 |
| 13 | 14 | Familiar | MonPerfFB | 285.864583 | 282.336957 | 96 | 92 | 75 | 70 | -5 | -4 | 76.0 |
| 14 | 15 | Familiar | MonPerfFB | 326.104651 | 337.407407 | 86 | 81 | 70 | 85 | 15 | -5 | 68.0 |
| 15 | 16 | Familiar | MonPerfFB | 312.484211 | 286.717391 | 95 | 92 | 85 | 65 | -20 | -3 | 70.0 |
| 16 | 17 | Familiar | MonPerfFB | 284.000000 | 277.275000 | 85 | 80 | 60 | 50 | -10 | -5 | 64.0 |
| 17 | 18 | Familiar | MonPerfFB | 302.711111 | 304.372340 | 90 | 94 | 60 | 50 | -10 | 4 | 58.0 |
| 18 | 19 | Familiar | MonPerfFB | 338.712329 | 325.590361 | 73 | 83 | 85 | 80 | -5 | 10 | 74.0 |
| 19 | 20 | Familiar | MonPerfFB | 300.451613 | 312.373626 | 93 | 91 | 85 | 85 | 0 | -2 | 73.0 |
| 20 | 21 | Familiar | MonPerfFB | 297.693878 | 290.775510 | 98 | 98 | 85 | 95 | 10 | 0 | 67.0 |
| 21 | 85 | Familiar | MonPerfFB | 284.894737 | 277.343750 | 95 | 96 | 85 | 85 | 0 | 1 | 68.0 |
| 22 | 86 | Familiar | MonPerfFB | 277.614583 | 292.362637 | 96 | 91 | 45 | 40 | -5 | -5 | 69.0 |
| 23 | 87 | Familiar | MonPerfFB | 294.000000 | 291.526882 | 96 | 93 | 90 | 75 | -15 | -3 | 70.0 |
| 24 | 88 | Familiar | MonPerfFB | 332.868421 | 320.767442 | 76 | 86 | 65 | 60 | -5 | 10 | 69.0 |
| 25 | 43 | Familiar | noMonPerfFB | 280.273684 | 299.527473 | 95 | 91 | 75 | 75 | 0 | -4 | 87.0 |
| 26 | 44 | Familiar | noMonPerfFB | 295.178947 | 285.913978 | 95 | 93 | 75 | 60 | -15 | -2 | 67.0 |
| 27 | 45 | Familiar | noMonPerfFB | 313.359375 | 302.558824 | 64 | 68 | 60 | 35 | -25 | 4 | 66.0 |
| 28 | 46 | Familiar | noMonPerfFB | 313.255319 | 315.022472 | 94 | 89 | 60 | 45 | -15 | -5 | 55.0 |
| 29 | 47 | Familiar | noMonPerfFB | 302.116279 | 297.097561 | 86 | 82 | 55 | 25 | -30 | -4 | 67.0 |
| 30 | 48 | Familiar | noMonPerfFB | 310.763441 | 289.533333 | 93 | 90 | 50 | 60 | 10 | -3 | 76.0 |
| 31 | 49 | Familiar | noMonPerfFB | 284.557895 | 276.119565 | 95 | 92 | 60 | 35 | -25 | -3 | 74.0 |
| 32 | 50 | Familiar | noMonPerfFB | 292.131313 | 279.437500 | 99 | 96 | 95 | 95 | 0 | -3 | 72.0 |
| 33 | 51 | Familiar | noMonPerfFB | 302.021739 | 299.443038 | 92 | 79 | 70 | 60 | -10 | -13 | 70.0 |
| 34 | 52 | Familiar | noMonPerfFB | 287.553191 | 296.106383 | 94 | 94 | 75 | 60 | -15 | 0 | 68.0 |
| 35 | 53 | Familiar | noMonPerfFB | 303.322581 | 315.551724 | 93 | 87 | 90 | 75 | -15 | -6 | 65.0 |
| 36 | 54 | Familiar | noMonPerfFB | 319.168831 | 306.734177 | 77 | 79 | 60 | 40 | -20 | 2 | 69.0 |
| 37 | 55 | Familiar | noMonPerfFB | 293.896552 | 307.790698 | 87 | 86 | 95 | 55 | -40 | -1 | 69.0 |
| 38 | 56 | Familiar | noMonPerfFB | 315.948454 | 307.989583 | 97 | 96 | 95 | 95 | 0 | -1 | 71.0 |
| 39 | 57 | Familiar | noMonPerfFB | 322.941176 | 324.022989 | 85 | 87 | 80 | 75 | -5 | 2 | 62.0 |
| 40 | 58 | Familiar | noMonPerfFB | 305.685393 | 295.434783 | 89 | 92 | 55 | 55 | 0 | 3 | 83.0 |
| 41 | 59 | Familiar | noMonPerfFB | 326.045455 | 321.525641 | 88 | 78 | 90 | 60 | -30 | -10 | 60.0 |
| 42 | 60 | Familiar | noMonPerfFB | 305.802198 | 301.825000 | 91 | 80 | 90 | 45 | -45 | -11 | 69.0 |
| 43 | 61 | Familiar | noMonPerfFB | 348.828947 | 349.239130 | 76 | 46 | 100 | 100 | 0 | -30 | 55.0 |
| 44 | 62 | Familiar | noMonPerfFB | 316.967391 | 287.845238 | 92 | 84 | 80 | 30 | -50 | -8 | 58.0 |
| 45 | 63 | Familiar | noMonPerfFB | 278.755102 | 308.584270 | 98 | 89 | 70 | 65 | -5 | -9 | 69.0 |
| 46 | 93 | Familiar | noMonPerfFB | 325.975610 | 333.025000 | 82 | 80 | 95 | 75 | -20 | -2 | 81.0 |
| 47 | 98 | Familiar | noMonPerfFB | 294.223404 | 281.360825 | 94 | 97 | 75 | 60 | -15 | 3 | 63.0 |
| 48 | 99 | Familiar | noMonPerfFB | 312.946809 | 333.577320 | 94 | 97 | 100 | 90 | -10 | 3 | 70.0 |
| 49 | 100 | Familiar | noMonPerfFB | 306.527473 | 288.031579 | 91 | 95 | 60 | 45 | -15 | 4 | NaN |
| 50 | 22 | Novel | MonPerfFB | 315.384615 | 322.741935 | 78 | 93 | 60 | 80 | 20 | 15 | 89.0 |
| 51 | 23 | Novel | MonPerfFB | 322.190476 | 320.989011 | 84 | 91 | 80 | 90 | 10 | 7 | 68.0 |
| 52 | 24 | Novel | MonPerfFB | 306.829787 | 325.326316 | 94 | 95 | 80 | 95 | 15 | 1 | 72.0 |
| 53 | 25 | Novel | MonPerfFB | 319.449438 | 311.750000 | 89 | 96 | 55 | 85 | 30 | 7 | 68.0 |
| 54 | 26 | Novel | MonPerfFB | 314.988636 | 295.148936 | 88 | 94 | 50 | 75 | 25 | 6 | 84.0 |
| 55 | 27 | Novel | MonPerfFB | 309.863158 | 302.610526 | 95 | 95 | 90 | 65 | -25 | 0 | 89.0 |
| 56 | 28 | Novel | MonPerfFB | 290.903226 | 302.701031 | 93 | 97 | 70 | 60 | -10 | 4 | 75.0 |
| 57 | 29 | Novel | MonPerfFB | 313.571429 | 297.315217 | 91 | 92 | 70 | 60 | -10 | 1 | 78.0 |
| 58 | 30 | Novel | MonPerfFB | 276.257732 | 304.333333 | 97 | 93 | 85 | 95 | 10 | -4 | 69.0 |
| 59 | 31 | Novel | MonPerfFB | 310.677778 | 324.426966 | 90 | 89 | 85 | 90 | 5 | -1 | 65.0 |
| 60 | 32 | Novel | MonPerfFB | 290.166667 | 300.375000 | 96 | 96 | 75 | 85 | 10 | 0 | 67.0 |
| 61 | 33 | Novel | MonPerfFB | 301.185567 | 275.690722 | 97 | 97 | 60 | 60 | 0 | 0 | 65.0 |
| 62 | 34 | Novel | MonPerfFB | 303.956522 | 291.970000 | 92 | 100 | 75 | 95 | 20 | 8 | 55.0 |
| 63 | 35 | Novel | MonPerfFB | 311.708333 | 298.050000 | 96 | 100 | 100 | 95 | -5 | 4 | 65.0 |
| 64 | 36 | Novel | MonPerfFB | 331.454545 | 332.563830 | 88 | 94 | 70 | 95 | 25 | 6 | 69.0 |
| 65 | 37 | Novel | MonPerfFB | 287.130435 | 294.416667 | 92 | 96 | 85 | 95 | 10 | 4 | 66.0 |
| 66 | 38 | Novel | MonPerfFB | 298.691489 | 313.458333 | 94 | 96 | 70 | 90 | 20 | 2 | 70.0 |
| 67 | 39 | Novel | MonPerfFB | 315.645833 | 301.142857 | 96 | 98 | 85 | 90 | 5 | 2 | 63.0 |
| 68 | 40 | Novel | MonPerfFB | 288.195122 | 302.861702 | 82 | 94 | 85 | 90 | 5 | 12 | 72.0 |
| 69 | 41 | Novel | MonPerfFB | 301.078652 | 300.819149 | 89 | 94 | 70 | 85 | 15 | 5 | 62.0 |
| 70 | 42 | Novel | MonPerfFB | 299.087912 | 274.442105 | 91 | 95 | 40 | 45 | 5 | 4 | 72.0 |
| 71 | 89 | Novel | MonPerfFB | 306.776596 | 312.906250 | 94 | 96 | 80 | 70 | -10 | 2 | 66.0 |
| 72 | 90 | Novel | MonPerfFB | 289.516854 | 292.904255 | 89 | 94 | 75 | 85 | 10 | 5 | 69.0 |
| 73 | 91 | Novel | MonPerfFB | 325.297619 | 327.252747 | 84 | 91 | 100 | 85 | -15 | 7 | 68.0 |
| 74 | 92 | Novel | MonPerfFB | 308.851064 | 306.587629 | 94 | 97 | 85 | 80 | -5 | 3 | 72.0 |
| 75 | 64 | Novel | noMonPerfFB | 334.978723 | 329.648936 | 94 | 94 | 90 | 60 | -30 | 0 | 68.0 |
| 76 | 65 | Novel | noMonPerfFB | 287.969388 | 289.191919 | 98 | 99 | 80 | 95 | 15 | 1 | 50.0 |
| 77 | 66 | Novel | noMonPerfFB | 303.847826 | 303.197802 | 92 | 91 | 85 | 85 | 0 | -1 | 71.0 |
| 78 | 67 | Novel | noMonPerfFB | 306.453608 | 306.551020 | 97 | 98 | 80 | 80 | 0 | 1 | 79.0 |
| 79 | 68 | Novel | noMonPerfFB | 314.726027 | 307.528736 | 73 | 87 | 65 | 55 | -10 | 14 | 61.0 |
| 80 | 69 | Novel | noMonPerfFB | 323.447059 | 322.681818 | 85 | 88 | 80 | 85 | 5 | 3 | 66.0 |
| 81 | 70 | Novel | noMonPerfFB | 307.166667 | 299.967742 | 96 | 93 | 85 | 80 | -5 | -3 | 76.0 |
| 82 | 71 | Novel | noMonPerfFB | 296.247312 | 300.239583 | 93 | 96 | 60 | 60 | 0 | 3 | 69.0 |
| 83 | 72 | Novel | noMonPerfFB | 302.921348 | 269.500000 | 89 | 90 | 25 | 40 | 15 | 1 | 70.0 |
| 84 | 73 | Novel | noMonPerfFB | 304.702128 | 301.950617 | 94 | 81 | 85 | 75 | -10 | -13 | 67.0 |
| 85 | 74 | Novel | noMonPerfFB | 280.936170 | 289.063830 | 94 | 94 | 65 | 80 | 15 | 0 | 63.0 |
| 86 | 75 | Novel | noMonPerfFB | 290.840426 | 295.344444 | 94 | 90 | 50 | 55 | 5 | -4 | 60.0 |
| 87 | 76 | Novel | noMonPerfFB | 302.540816 | 313.021053 | 98 | 95 | 95 | 90 | -5 | -3 | 58.0 |
| 88 | 77 | Novel | noMonPerfFB | 299.534091 | 280.163043 | 88 | 92 | 40 | 60 | 20 | 4 | 75.0 |
| 89 | 78 | Novel | noMonPerfFB | 304.367347 | 309.616162 | 98 | 99 | 85 | 100 | 15 | 1 | 76.0 |
| 90 | 79 | Novel | noMonPerfFB | 310.525641 | 312.487500 | 78 | 80 | 80 | 65 | -15 | 2 | 69.0 |
| 91 | 80 | Novel | noMonPerfFB | 302.956989 | 295.822222 | 93 | 90 | 90 | 75 | -15 | -3 | 70.0 |
| 92 | 81 | Novel | noMonPerfFB | 323.372093 | 315.788235 | 86 | 85 | 95 | 90 | -5 | -1 | 69.0 |
| 93 | 82 | Novel | noMonPerfFB | 321.645161 | 309.923913 | 93 | 92 | 65 | 90 | 25 | -1 | 64.0 |
| 94 | 83 | Novel | noMonPerfFB | 308.421687 | 311.241758 | 83 | 91 | 75 | 80 | 5 | 8 | 72.0 |
| 95 | 84 | Novel | noMonPerfFB | 305.277778 | 318.870968 | 90 | 93 | 80 | 80 | 0 | 3 | 73.0 |
| 96 | 94 | Novel | noMonPerfFB | 313.907216 | 318.114583 | 97 | 96 | 100 | 90 | -10 | -1 | 64.0 |
| 97 | 95 | Novel | noMonPerfFB | 304.851064 | 298.391753 | 94 | 97 | 75 | 65 | -10 | 3 | 75.0 |
| 98 | 96 | Novel | noMonPerfFB | 311.000000 | 314.119565 | 92 | 92 | 80 | 75 | -5 | 0 | 73.0 |
| 99 | 97 | Novel | noMonPerfFB | 365.159091 | 362.190476 | 44 | 42 | 100 | 95 | -5 | -2 | 76.0 |

```
C:\Users\ahmet\Anaconda3\lib\site-packages\scipy\stats\stats.py:1713: FutureWarning: Using a non-tuple sequence for multidimensional indexing is deprecated; use `arr[tuple(seq)]` instead of `arr[seq]`. In the future this will be interpreted as an array index, `arr[np.array(seq)]`, which will result either in an error or a different result.
  return np.add.reduce(sorted[indexer] * weights, axis=axis) / sumval
```

In [6]:

```
sns.set(style="white", context="notebook", font="Times New Roman", font_scale=1.3)
ax = sns.barplot(x="FB_Cond", y="Accuracy", hue="Congruency", palette=["#1d47f5", "#d12fdf"], ci=68, capsize=0.02, data=stacked_data_nov)
plt.title("Novel", weight="bold", y=1.08, fontsize=20)
ax.set_ylim(50,88)
ax.set_xlabel("")
ymajor = np.arange(50, 86, 5)
ax.set_yticks(ymajor)
ax.set_ylabel("NoGo Accuracy (%)", weight="bold", labelpad=5, fontsize=20)
ax.set_xticklabels(["Feedback", "No Feedback"], fontsize=20)
ax.legend_.remove()
sns.despine(bottom=False)
#add sig asterisks and line
#first
x1, x2 = -0.20, 0.18
y, h, col = stacked_data_fam["Accuracy"].mean()+13.3, 0.5, "k"
plt.plot([x1, x1, x2, x2], [y, y+h, y+h, y], lw=1, c=col)
plt.text((x1+x2)*0.5, y+h-0.1, "*", ha="center", va="bottom", color=col)
#second
#x3, x4 = 0.79, 1.17
#y, h, col = stacked_data_fam["Accuracy"].mean()+14, 0.5, "k"
#plt.plot([x3, x3, x4, x4], [y, y+h, y+h, y], lw=1, c=col)
#plt.text((x3+x4)*0.5, y+h-0.1, "*", ha="center", va="bottom", color=col)
plt.text(-0.1, 1.1, "B", weight="bold", fontsize=25, ha="left", va="bottom", transform=ax.transAxes)
#plt.savefig("Exp4_NoGo_graph_Nov.tiff", bbox_inches="tight", dpi=500)
plt.show()
```

```
C:\Users\ahmet\Anaconda3\lib\site-packages\scipy\stats\stats.py:1713: FutureWarning: Using a non-tuple sequence for multidimensional indexing is deprecated; use `arr[tuple(seq)]` instead of `arr[seq]`. In the future this will be interpreted as an array index, `arr[np.array(seq)]`, which will result either in an error or a different result.
  return np.add.reduce(sorted[indexer] * weights, axis=axis) / sumval
```

```
C:\Users\ahmet\Anaconda3\lib\site-packages\scipy\stats\stats.py:1713: FutureWarning: Using a non-tuple sequence for multidimensional indexing is deprecated; use `arr[tuple(seq)]` instead of `arr[seq]`. In the future this will be interpreted as an array index, `arr[np.array(seq)]`, which will result either in an error or a different result.
  return np.add.reduce(sorted[indexer] * weights, axis=axis) / sumval
```

```
C:\Users\ahmet\Anaconda3\lib\site-packages\scipy\stats\stats.py:1713: FutureWarning: Using a non-tuple sequence for multidimensional indexing is deprecated; use `arr[tuple(seq)]` instead of `arr[seq]`. In the future this will be interpreted as an array index, `arr[np.array(seq)]`, which will result either in an error or a different result.
  return np.add.reduce(sorted[indexer] * weights, axis=axis) / sumval
```

In [ ]:
