## Supplement for "Demonstrating and disrupting well-learned habits"

**Experiment 1 – Omnibus regression to confirm mapping-related accuracy impairment**

**Primary measure of outcome-sensitivity: NoGo accuracy**

We derived a ΔNoGo_Accuracy (i.e., change in NoGo accuracy scores across mappings) DV to quantify the mapping-related impairment for each subject. This ΔNoGo_Accuracy variable serves as the primary measure of outcome-sensitivity, in that a greater impairment represents greater outcome-insensitivity. Specifically, difficulty overriding the Familiar Red–NoGo association for the Green–NoGo association indicates a cue-driven habit. In contrast, we would not expect a pronounced ΔNoGo_Accuracy value when participants manage Novel NoGo contingencies (i.e., blue–NoGo and purple–NoGo should yield similar accuracy scores). Participants with DV standardized residual values below -3.3 and above +3.3 were identified as outliers [1]. In such cases, we performed identical analyses without outlier participants to verify robustness of findings, but only report these excluded analyses if outliers produced substantial changes in statistical significance.

We employed a hierarchical multiple regression model to extract the predictive strength of the between-group Condition variable while controlling for age, gender, order of mapping phase (i.e., whether a subject completed a particular color-response mapping first), and impulsivity. We entered the controlled Age, Gender, Order, and Impulsivity regressors into the first, and the Condition regressor of interest into the second step of the model. Therefore, our hierarchical multiple regression model yielded an R^2^ change value (ΔR^2^) for Condition, determining whether mapping-related impairments are predicted specifically by Condition (i.e., contingency change in Familiar versus Novel stimuli). We also derived the corresponding *F*_change_ value, which compares the predictive strength of the variables in the second step of the model with those in the first step (i.e., confirming whether ΔR^2^ reflects a significant change in the model’s predictive strength).

The regression model met the assumptions of normality and homoscedasticity. Multicollinearity tests produced negligible Variance Inflation Factors (VIF), confirming linearity assumptions of the regression (VIF for all variables < 1.09). Model 1, a linear combination of the controlled variables of Age, Gender, Order, and Impulsivity, did not significantly predict outcome-sensitivity: *F*(4,45) = 0.46, *p* = .767, and only explained 4% of the variance in ΔNoGo_Accuracy (R^2^ = .04). Additionally, no controlled regressor independently predicted a change in ΔNoGo_Accuracy (all β coefficient *p’*s. > .05). In the second step of the regression, the inclusion of Condition as a regressor explained an additional 15.5% of the variance in outcome-sensitivity: β_Condition_ = -.40, ΔR^2^ = .15, *F*_change_ (1,44) = 8.47, *p* = .006—a significant contribution. Thus, the addition of the significant Condition regressor rendered the entirety of Model 2 a near-significant predictor of outcome-sensitivity, despite the null contributions from Age, Gender, Order, and Impulsivity: *F*(5,44) = 2.12, *p* = .081 (see S1 Table).

**S1 Table. Summary of the Hierarchical Multiple Regression Model for Outcome-Insensitivity as Assayed by ΔNoGo_Accuracy.**

| Variable | *Toler.* | *VIF* | *B* | *SE* | *β* | *t* | *sig.* |
| --- | --- | --- | --- | --- | --- | --- | --- |
| **Model 1** |  |  |  |  |  |  |  |
| Age | .98 | 1.02 | 0.57 | 0.82 | .10 | 0.69 | .493 |
| Gender | .97 | 1.04 | -2.69 | 5.08 | -.08 | -0.53 | .599 |
| Impulsivity | .93 | 1.07 | -0.25 | 0.35 | -.11 | -0.71 | .483 |
| Order | .96 | 1.04 | 3.12 | 4.88 | .95 | 0.64 | .527 |
| **Model 2** |  |  |  |  |  |  |  |
| Age | .97 | 1.03 | 0.34 | 0.77 | .06 | 0.44 | .662 |
| Gender | .96 | 1.05 | -1.30 | 4.73 | -.04 | -0.27 | .784 |
| Impulsivity | .93 | 1.08 | -0.33 | 0.33 | -.14 | -0.99 | .326 |
| Order | .96 | 1.04 | 2.88 | 4.52 | .09 | 0.64 | .527 |
| Condition | .98 | 1.02 | -13.06 | 4.50 | -.40^**^ | -2.91 | **.006** |

| Model Summary Statistics | | | | | | |
| --- | --- | --- | --- | --- | --- | --- |
| Model | *R^2^* | *F* | *F sig.* | *ΔR^2^* | *F*_change_ | *F*_change_ *sig.* |
| **Model 1** | .04 | 0.46 | .767 |  |  |  |
| **Model 2** | .19 | 2.12 | .081 | .15 | 8.47 | **.006** |

*Note: Top layer of table depicts all regressors included in the hierarchical model and their respective statistics. Bottom layer of table, Model Summary Statistics, depicts the predictive strength of each model. Delta R^2^ (ΔR^2^) and corresponding F_change_ values denote the specific improvement of Model 2 over Model 1 in predicting the dependent variable. Toler. = Tolerance; VIF = Variance Inflation Factor. Significant p-values (alpha = .05) depicted in bold typeface.*

These results also suggest that the differential mapping-related impairment observed across Familiar and Novel conditions is not due to the order in which participants managed color-response mappings. The Order variable did not significantly predict ΔNoGo_Accuracy in our model (β = .09, *p* = .527). We found no interaction between factors of Order and Mapping in NoGo accuracy as a result of the repeated measures ANOVA: *F*(1,48) = 0.35, *p* = .555, η_p_^2^ < .01. We performed the same ANOVA separately in Familiar and Novel conditions and observed no significant interactions in either group (*p*’s > .05).

**Secondary measure of outcome-sensitivity: Go accuracy**

We performed an identical omnibus regression using ΔGo_Accuracy as DV—the secondary assay of outcome-sensitivity. The regression model met the assumptions of normality and homoscedasticity. Multicollinearity tests produced negligible Variance Inflation Factors (VIF), confirming the assumption of linearity in the regression model (VIF for all variables < 1.09).

Collectively, the linear combination of Age, Gender, Order, and Impulsivity did not significantly predict mapping-related impairments in Go accuracy: *F*(4,45) = 2.18, *p* = .087. As depicted in S2 Table, closer examination of the individual regressors revealed a significant role played by Age, such that older participants suffered a greater mapping-related Go accuracy impairment: β_Age_ = -.31, *p* = .027. The inclusion of the Condition regressor significantly improved the predictive strength of the model in step 2: β_Condition_ = -.27, ΔR^2^ = .07, *F*_change_(1,44) = 4.10, *p* = .049, and the significant contribution of Age remained: β_Age_ = -.34, *p* = .014. Although age was a significant predictor of change in Go accuracy, because we had no *a priori* hypothesis, and the correlational direction of this relationship varied across conditions (Familiar Condition Pearson’s *r* = .43; Novel Condition Pearson’s *r* = -.70), we refrain from further age-related speculation.

**S2 Table. Summary of the Hierarchical Multiple Regression Model for Outcome-Insensitivity as Assayed by ΔGo_Accuracy.**

| Variable | *Toler.* | *VIF* | *B* | *SE* | *β* | *t* | *sig.* |
| --- | --- | --- | --- | --- | --- | --- | --- |
| **Model 1** |  |  |  |  |  |  |  |
| Age | .98 | 1.02 | -0.60 | 0.26 | -.31 | -2.29^*^ | **.027** |
| Gender | .97 | 1.04 | 1.83 | 1.61 | .16 | 1.14 | .261 |
| Impulsivity | .93 | 1.07 | -0.02 | 0.11 | -.03 | -0.20 | .843 |
| Order | .96 | 1.04 | -2.02 | 1.55 | -.18 | -1.30 | .199 |
| **Model 2** |  |  |  |  |  |  |  |
| Age | .97 | 1.03 | -0.65 | 0.25 | -.34^*^ | -2.56 | **.014** |
| Gender | .96 | 1.05 | 2.15 | 1.56 | .19 | 1.38 | .176 |
| Impulsivity | .93 | 1.08 | -0.04 | 0.11 | -.05 | -0.37 | .716 |
| Order | .96 | 1.04 | -2.07 | 1.50 | -.19 | -1.38 | .174 |
| Condition | .98 | 1.02 | -3.01 | 1.49 | -.27^**^ | -2.03 | **.049** |

| Model Summary Statistics | | | | | | |
| --- | --- | --- | --- | --- | --- | --- |
| Model | *R^2^* | *F* | *F sig.* | *ΔR^2^* | *F*_change_ | *F*_change_ *sig.* |
| **Model 1** | .16 | 2.18 | .087 |  |  |  |
| **Model 2** | .23 | 2.68 | **.034** | .07 | 4.10 | **.049** |

*Note: Top layer of table depicts all regressors included in the hierarchical model and their respective statistics. Bottom layer of table, Model Summary Statistics, depicts the predictive strength of each model. Delta R^2^ (ΔR^2^) and corresponding F_change_ values denote the specific improvement of Model 2 over Model 1 in predicting the dependent variable. Toler. = Tolerance; VIF = Variance Inflation Factor. Significant p-values (alpha = .05) depicted in bold typeface.*

Similar to our primary assay of outcome-sensitivity, change in Go accuracy was not due to the order in which participants managed color-response mappings (β_Order_ = -.19, *p* = .174). There was no significant Order x Mapping interaction in Go accuracy: *F*(1,48) = 2.26, *p* = .140, η_p_^2^ = .04. We performed the same ANOVA separately in Familiar and Novel conditions and observed no significant interactions in either group (*p*’s > .05).

**Experiment 2 – Omnibus regression to illustrate mapping-related impairment while testing the effect of cumulative performance feedback**

**Primary measure of outcome-sensitivity: NoGo accuracy**

Similar to Experiment 1, we derived a ΔNoGo_Accuracy DV to quantify the mapping-related impairment for each participant as the primary assay of outcome-sensitivity. We employed a hierarchical multiple regression model to extract the predictive strengths of the between-group regressors, Condition and Feedback, while controlling for age, gender, and impulsivity. The resulting ΔR^2^ value for the contributions of Condition and Feedback determined whether mapping-related impairments are predicted specifically by Condition (i.e., Familiar versus Novel stimuli), and whether the cumulative performance feedback manipulation plays a role in affecting motivational control. A corresponding *F*_change_ value was derived to confirm whether ΔR^2^ reflects a significant change in the model’s predictive strength. Participants with DV standardized residual values below -3.3 and above 3.3 were identified as outliers [1]. In such cases, we performed identical analyses without outlier participants to verify robustness of findings, but only report these excluded analyses if outliers produced substantial changes in statistical significance.

The regression model met the assumptions of normality and homoscedasticity. Multicollinearity tests produced negligible Variance Inflation Factors (VIF), confirming the assumption of linearity in the regression model (VIF for all variables < 1.04; see S3 Table).

Model 1, a linear combination of the controlled variables Age, Gender, and Impulsivity, did not significantly predict outcome-sensitivity: *F*(3,96) = 0.18, *p* = .91, and only explained 0.6% of the variance in ΔNoGo_Accuracy. Additionally, no controlled regressor independently predicted a change in ΔNoGo_Accuracy (all β coefficient *p’*s. > .05; see S3 Table). In the second step of the regression, the inclusion of the Condition and Feedback regressors explained an additional 14.5% of the variance in outcome-sensitivity: β_Condition_ = -.34, *p* = .001, β_Feedback_ = .18, *p* = .07, ΔR^2^ = .14, *F*_change_ (2,94) = 8.03, *p* = .001—rendering the entirety of Model 2 a significant predictor of outcome-sensitivity: *F*(5,94) = 3.34, *p* = .008 (see S3 Table).

**S3 Table. Summary of the Hierarchical Multiple Regression Model for Outcome-Insensitivity as Assayed by ΔNoGo_Accuracy.**

| Variable | *Toler.* | *VIF* | *B* | *SE* | *β* | *t* | *sig.* |
| --- | --- | --- | --- | --- | --- | --- | --- |
| **Model 1** |  |  |  |  |  |  |  |
| Age | .99 | 1.01 | -0.31 | 0.58 | -.05 | -0.54 | .590 |
| Gender | .98 | 1.02 | -0.31 | 3.75 | -.01 | -0.08 | .935 |
| Impulsivity | .98 | 1.02 | 0.13 | 0.27 | .05 | 0.48 | .633 |
| **Model 2** |  |  |  |  |  |  |  |
| Age | .98 | 1.02 | -0.54 | 0.54 | -.09 | -1.00 | .319 |
| Gender | .96 | 1.04 | 1.10 | 3.53 | .03 | 0.31 | .756 |
| Impulsivity | .98 | 1.02 | 0.17 | 0.25 | .06 | 0.68 | .497 |
| Condition | .98 | 1.02 | -11.74 | 3.29 | -.34^**^ | -3.57 | **.001** |
| Feedback | .99 | 1.01 | 6.06 | 3.28 | .18 | 1.85 | .067 |

| Model Summary Statistics | | | | | | |
| --- | --- | --- | --- | --- | --- | --- |
| Model | *R^2^* | *F* | *F sig.* | *ΔR^2^* | *F_change_* | *F_change_ sig.* |
| **Model 1** | .01 | 0.18 | .910 |  |  |  |
| **Model 2** | .15 | 3.34 | **.008** | .14 | 8.03 | **.001** |

*Note: Top layer of table depicts all regressors included in the hierarchical model and their respective statistics. Bottom layer of table, Model Summary Statistics, depicts the predictive strength of each model. Delta R^2^ (ΔR^2^) and corresponding F_change_ values denote the specific improvement of Model 2 over Model 1 in predicting the dependent variable. Toler. = Tolerance; VIF = Variance Inflation Factor. Significant p-values (alpha = .05) depicted in bold typeface.*

We hypothesized that performance feedback may be a salient factor that can potentially restore goal-directed control when managing these well-established associations. However, cumulative performance feedback did not break the habits elicited by these familiar stimuli. As seen in the hierarchical multiple regression model, although Condition yielded differential mapping-related NoGo accuracy changes across familiar and novel conditions, the Feedback regressor was not a significant predictor of outcome-sensitivity.

*Secondary measure of outcome-insensitivity:* ΔGo_Accuracy

We performed an identical omnibus hierarchical regression using ΔGo_Accuracy, which serves as our secondary measure of outcome-sensitivity. The regression model met the assumptions of normality and homoscedasticity. Multicollinearity tests produced negligible VIFs, confirming the assumption of linearity in the regression model (VIF for all variables < 1.04; see S4 Table). Two participants were identified as outliers due to a standardized residual values falling outside the predetermined range [1]. Identical analyses without outlier data produced no substantial changes in the statistical findings below.

Collectively, the linear combination of Age, Gender, and Impulsivity did not significantly predict mapping-related change in Go accuracy: *F*(3,96) = 0.31, *p* = .81. As depicted in S4 Table, closer examination of the individual regressors revealed no significant role played by any of the controlled variables in Model 1 (all β coefficient *p’*s. > .05; see S4 Table). In the second step of the regression, the inclusion of the Condition and Feedback regressors explained an additional 17.7% of the variance in Go accuracy impairment: β_Condition_ = -.32, *p* = .001, β_Feedback_ = .28, *p* = .003, ΔR^2^ = .18, *F*_change_(2,94) = 10.23, *p* < .001—rendering Model 2 a significant predictor of ΔGo_Accuracy: *F*(5,94) = 4.32, *p* = .001 (see S4 Table).

**S4 Table. Summary of the Hierarchical Multiple Regression Model for Outcome-Insensitivity as Assayed by ΔGo_Accuracy.**

| Variable | *Toler.* | *VIF* | *B* | *SE* | *β* | *t* | *sig.* |
| --- | --- | --- | --- | --- | --- | --- | --- |
| **Model 1** |  |  |  |  |  |  |  |
| Age | .99 | 1.01 | -0.11 | 0.21 | -.54 | -0.54 | .591 |
| Gender | .98 | 1.02 | -1.00 | 1.38 | -.07 | -0.73 | .469 |
| Impulsivity | .98 | 1.02 | -0.05 | 0.10 | -.05 | -0.53 | .600 |
| **Model 2** |  |  |  |  |  |  |  |
| Age | .98 | 1.02 | -0.22 | 0.20 | -.11 | -1.13 | .261 |
| Gender | .96 | 1.04 | -0.51 | 1.27 | -.04 | -0.40 | .692 |
| Impulsivity | .98 | 1.02 | -0.04 | 0.09 | -.04 | -0.43 | .670 |
| Condition | .98 | 1.02 | -4.03 | 1.19 | -.32^**^ | -3.39 | **.001** |
| Feedback | .99 | 1.01 | 3.59 | 1.18 | .28^**^ | 3.03 | **.003** |

| Model Summary Statistics | | | | | | |
| --- | --- | --- | --- | --- | --- | --- |
| Model | *R^2^* | *F* | *F sig.* | *ΔR^2^* | *F*_change_ | *F*_change_ *sig.* |
| **Model 1** | .01 | 0.31 | .815 |  |  |  |
| **Model 2** | .18 | 4.32 | **.001** | .18 | 10.23 | **<.001** |

*Note: Top layer of table depicts all regressors included in the hierarchical model and their respective statistics. Bottom layer of table, Model Summary Statistics, depicts the predictive strength of each model. Delta R^2^ (ΔR^2^) and corresponding F_change_ values denote the specific improvement of Model 2 over Model 1 in predicting the dependent variable. Toler. = Tolerance; VIF = Variance Inflation Factor. Significant p-values (alpha = .05) depicted in bold typeface.*

Although these hierarchical regression results regarding ΔGo_Accuracy suggest that cumulative performance feedback has significant predictive strength, the mixed-design ANOVAs in the main text confirm that cumulative performance feedback has a significant effect on Go actions only in the Familiar condition.

**Experiment 3 – Omnibus regression to illustrate mapping-related impairment while testing the effect of dual feedback—cumulative performance and monetary feedback**

To detect the potential habit-disrupting effects of dual feedback (i.e., paired monetary and cumulative performance feedback), we employed a hierarchical multiple regression model to extract the predictive strengths of the between-group regressors, Condition and Feedback, while controlling for age, gender, and impulsivity. We entered the controlled Age, Gender, and Impulsivity regressors into the first, and Condition and Feedback regressors of interest into the second step of the model to yield an R^2^ change (ΔR^2^) value that quantifies the contributions of Condition and Feedback. This allowed us to confirm whether mapping-related impairments are predicted specifically by Condition (i.e., Familiar versus Novel stimuli), and whether dual feedback affected motivational control. A corresponding *F*_change_ value was derived to confirm whether ΔR^2^ reflects a significant change in the model’s predictive strength. Participants with DV standardized residual values below -3.3 and above 3.3 were identified as outliers [1]. In such cases, we performed identical analyses without outlier participants to verify robustness of findings, but only report these excluded analyses if outliers produced substantial changes in statistical significance.

The regression model met the assumptions of normality and homoscedasticity. Multicollinearity tests produced negligible Variance Inflation Factors (VIF), confirming linearity assumptions in the regression (VIF for all variables < 1.15; see S5 Table).

As hypothesized, Model 1, a linear combination of the controlled variables Age, Gender, and Impulsivity, did not significantly predict outcome-sensitivity: *F*(3,96) = 0.12, *p* = .95, and only explained 0.4% of the variance in ΔNoGo_Accuracy (R^2^ = .004). Additionally, none of these regressors independently predicted ΔNoGo_Accuracy (all β coefficient *p’*s > .05; see S5 Table). In the second step of the regression, the inclusion of the Condition and Feedback regressors explained an additional 26.6% of the variance in outcome-sensitivity: β_Condition_ = -.43, *p* < .001, β_Feedback_ = .28, *p* = .003, ΔR^2^ = .27, *F*_change_(2,94) = 17.16, *p* < .001—rendering Model 2 a significant predictor of outcome-sensitivity: *F*(5,94) = 6.96, *p* < .001 (see S5 Table).

**S5 Table. Summary of the Hierarchical Multiple Regression Model for Outcome-Insensitivity as Assayed by ΔNoGo_Accuracy.**

| Variable | *Toler.* | *VIF* | *B* | *SE* | *β* | *t* | *sig.* |
| --- | --- | --- | --- | --- | --- | --- | --- |
| **Model 1** |  |  |  |  |  |  |  |
| Age | .97 | 1.03 | 0.16 | 0.57 | .03 | 0.29 | .775 |
| Gender | .99 | 1.00 | 1.31 | 3.69 | .04 | 0.35 | .724 |
| Impulsivity | .97 | 1.03 | 0.09 | 0.22 | .04 | 0.40 | .692 |
| **Model 2** |  |  |  |  |  |  |  |
| Age | .94 | 1.06 | 0.24 | 0.50 | .04 | 0.48 | .635 |
| Gender | .87 | 1.14 | 0.14 | 3.41 | .004 | 0.04 | .968 |
| Impulsivity | .91 | 1.09 | 0.01 | 0.20 | .01 | 0.08 | .939 |
| Condition | .92 | 1.08 | -13.42 | 2.83 | -.43^***^ | -4.74 | **<.001** |
| Feedback | .89 | 1.13 | 8.76 | 2.89 | .28^**^ | 3.03 | **.003** |

| Model Summary Statistics | | | | | | |
| --- | --- | --- | --- | --- | --- | --- |
| Model | *R^2^* | *F* | *F sig.* | *ΔR^2^* | *F*_change_ | *F*_change_ *sig.* |
| **Model 1** | .004 | 0.12 | .950 |  |  |  |
| **Model 2** | .52 | 6.96 | **<.001** | .27 | 17.16 | **<.001** |

*Note: Top layer of table depicts all regressors included in the hierarchical model and their respective statistics. Bottom layer of table, Model Summary Statistics, depicts the predictive strength of each model. Delta R^2^ (ΔR^2^) and corresponding F_change_ values denote the specific improvement of Model 2 over Model 1 in predicting the dependent variable. Toler. = Tolerance; VIF = Variance Inflation Factor. Significant p-values (alpha = .05) depicted in bold typeface.*

As hypothesized, the delivery of cumulative performance and monetary feedback disrupted habits (i.e., prevent a significant incongruency-related impairment in NoGo accuracy to familiar lights) and improved goal-directed control (i.e., significantly increase NoGo accuracy to novel stimuli). In other words, these regression data suggest that the differential mapping-related NoGo impairment is replicated in Experiment 3, in that the stimulus condition predicts changes in accuracy, and importantly, dual feedback is able to significantly predict improvements in performance.

*Secondary index of outcome-sensitivity: Go Accuracy*

We performed identical regression analyses using ΔGo_Accuracy as DV. The regression model met the assumptions of normality and homoscedasticity. Multicollinearity tests produced negligible VIFs, confirming the assumption of linearity in the regression model (VIF for all variables < 1.15). One participant was identified as an outlier due to a standardized residual value less than -3.3 [1]. Identical analyses without the outlier data produced no substantial changes in the statistical findings reported below; distinctions are specified where relevant.

Collectively, the linear combination of Age, Gender, and Impulsivity did not significantly predict mapping-related impairments in Go accuracy: *F*(3,96) = 1.67, *p* = .18. As depicted in S6 Table, Model 1 only explained 5% of the variance (R^2^ = .05), and of all the controlled variables in Model 1, only the Impulsivity variable significantly predicted ΔGo_Accuracy: β_Impulsivity_ = .21, *p* = .04 (all other β coefficient *p’*s. > .05). In the second step of our regression model, the inclusion of the Condition and Feedback regressors explained an additional 21.7% of the variance in Go accuracy impairment: β_Condition_ = -.36, *p* < .001, β_Feedback_ = .28, *p* = .004, ΔR^2^ = .21, *F*_change_(2,94) = 13.11, *p* < .001—rendering Model 2 a significant predictor of ΔGo_Accuracy: *F*(5,94) = 6.50, *p* < .001 (see S6 Table). The predictive strength of the Impulsivity variable diminished below significance in Model 2 (β_Impulsivity_ = .17, *p* = .07). When reanalyzed without outlier data, Impulsivity did not predict ΔGo_Accuracy in either model (both *p*’s > .05). Given its lack of significant contributions in Experiments 1 and 2, and sensitivity to outlier correction in Experiment 3, we refrain from speculating further regarding the robustness of Impulsivity as a predictor.

**S6 Table. Summary of the Hierarchical Multiple Regression Model for Outcome-Insensitivity as Assayed by ΔGo_Accuracy.**

| Variable | *Toler.* | *VIF* | *B* | *SE* | *β* | *t* | *sig.* |
| --- | --- | --- | --- | --- | --- | --- | --- |
| **Model 1** |  |  |  |  |  |  |  |
| Age | .97 | 1.03 | 0.18 | 0.21 | .09 | 0.87 | .384 |
| Gender | .99 | 1.00 | 0.78 | 1.36 | .06 | 0.57 | .569 |
| Impulsivity | .97 | 1.03 | 0.17 | 0.08 | .21^*^ | 2.06 | **.042** |
| **Model 2** |  |  |  |  |  |  |  |
| Age | .94 | 1.06 | 0.20 | 0.19 | .09 | 1.03 | .306 |
| Gender | .87 | 1.14 | 0.55 | 1.30 | .04 | 0.42 | .673 |
| Impulsivity | .91 | 1.09 | 0.14 | 0.08 | .17 | 1.84 | .069 |
| Condition | .92 | 1.08 | -4.22 | 1.08 | -.36^***^ | -3.92 | **<.001** |
| Feedback | .89 | 1.13 | 3.25 | 1.10 | .28^**^ | 2.95 | **.004** |

| Model Summary Statistics | | | | | | |
| --- | --- | --- | --- | --- | --- | --- |
| Model | *R^2^* | *F* | *F sig.* | *ΔR^2^* | *F*_change_ | *F*_change_ *sig.* |
| **Model 1** | .05 | 1.67 | .178 |  |  |  |
| **Model 2** | .26 | 6.50 | **<.001** | .21 | 13.11 | **<.001** |

*Note: Top layer of table depicts all regressors included in the hierarchical model and their respective statistics. Bottom layer of table, Model Summary Statistics, depicts the predictive strength of each model. Delta (Δ) values denote the specific improvement of Model 2 over Model 1 in predicting the dependent variable. Toler. = Tolerance; VIF = Variance Inflation Factor. Significant p-values (alpha = .05) depicted in bold typeface.*

The significant Condition and Feedback regressors indicate that like Experiments 1 and 2, Condition (Familiar vs. Novel) differentially yields changes in Go accuracy, and Feedback has a significant improvement effect on Go accuracy. It should be noted that our mixed-design ANOVAs (reported in the main text) reveal a significant feedback effect on Go actions only in the Novel condition. We can thus conclude that dual feedback does not significantly disrupt Go habits, though it does significantly promote goal-directed Go actions.
